# Functional metagenomic screening in microfluidic droplets identifies a β-glucuronidase in an unprecedented sequence neighbourhood

**DOI:** 10.1101/2022.04.25.489410

**Authors:** Stefanie Neun, Paul Brear, Eleanor Campbell, Theodora Tryfona, Kamel El Omari, Armin Wagner, Paul Dupree, Marko Hyvönen, Florian Hollfelder

## Abstract

The abundance of recorded protein sequence data stands in contrast to the small number of experimentally verified functional annotation. Here we screened a million-membered metagenomic library at ultrahigh throughput in microfluidic droplets for β-glucuronidase activity. We identified SN243, a genuine β-glucuronidase with little homology to previously studied enzymes of this type, as a glycoside hydrolase (GH) 3 family member. This GH family had no recorded evidence of β-glucuronidases at the outset of this study, showing that a functional metagenomic approach can shed light on assignments that are currently ‘unpredictable’ by bioinformatics. Kinetic analyses of SN243 characterised it as a promiscuous catalyst and structural analysis suggests regions of divergence from homologous GH3 members creating a wide-open active site. With a screening throughput of >10^7^ library members per day, picolitre volume microfluidic droplets enable functional assignments that complement current enzyme database dictionaries and provide bridgeheads for the annotation of unexplored sequence space.

## INTRODUCTION

Enzymes are the ultimate reagents for efficient, sustainable biocatalytic production lines or, as biologics, for medical and nutritional applications. However, the set of currently known enzymes does not cover all desirable activities, so accelerated discovery of new enzymes is imperative. Activity prediction using sequence homology, boosted by the enormous amounts of metagenomic data becoming available (e.g. in the Mgnify database),^1^ is a powerful tool for characterising closely related enzymes, but difficult when no experimentally characterized precedent exists in nearby sequence space. The functional annotation available in comprehensive databases (such as BRENDA^2^ and CAZy; http://www.cazy.org/)^3^ captures only a fraction of the enzyme thesaurus available to us. Indeed, the amount of sequence information available is growing much faster than the number of characterized enzymes,^1, 4^ raising the question of how ‘bridgeheads’ of functional annotation in unexplored sequence neighbourhoods can be found. Often there will not be just one functional assignment to be made: promiscuous side activities – recognised to be an important engine in interconversion of activities in natural evolution^5, 6^ - are of interest, as they can also provide a springboard for directed evolution^7, 8^ (in analogy to their role in Nature). If assignment of the principal activities by homology is already difficult, then prediction of promiscuous functions will be even harder (or impossible), as the catalytic machinery for native and promiscuous reactions is often similar.^9^

*Functional* metagenomics provides an alternative to *sequence-based* exploration of such hidden activities: experimental screening for enzyme function by detecting actual catalytic turnover as a readout is the most direct way to determine function without reliance on sequence homology.^10, 11^ However, the wet lab work to provide evidence for a correct functional assignment is cumbersome, and the number of ‘hits’ in a given library is generally low (estimated ∼1 in 10^4^-10^5^).^12^ Droplet microfluidics allows a massive scale-up of screening throughput (∼10^8^ droplets in a single day) and the picolitre assay volume of the droplets makes enzyme screening extremely economical.^13, 14^ In this experimental approach, water-in-oil droplets serve as miniature reaction compartments and maintain the genotype-phenotype linkage.^15–19^ Active catalysts are detected via an optical readout followed by on-chip^20, 21^ or flow cytometric sorting^22, 23^ and recovery of the encoding genes.^24^ Enzymes for a non-natural reaction, phosphotriester hydrolysis,^25^ have already been identified in metagenomic libraries by on-chip droplet microfluidics. In this work, we explore its potential to annotate sequences for natural reactions.

Specifically we target enzymes that hydrolyse *O*-linked β-linked glucuronic acids, ubiquitous sugar acids present in various plant polysaccharides,^26^ bacterial capsular polysaccharides,^27^ mammalian glycosaminoglycans,^28^ or conjugates excreted by the liver of vertebrates.^29^ Most currently known bacterial β-glucuronidases (E.C. 3.2.1.31) are members of the glycoside hydrolase family (GH) 2^3, 30^ (Figure S5 and Table S2, SI). Therefore, the question arises whether other bacterial β-glucuronidases exist elsewhere that are not revealed in traditional sequencing-based homology searches, but amenable to discovery by ultrahigh throughput functional metagenomics.

To this end we screen a million-membered metagenomic library for β-glucuronidase activity and uncover an enzyme that no *in silico* classification method would have assigned to this activity; accordingly, no evidence of a combination of this activity and the respective GH in the CAZy database had been recorded at the outset of this study. The genuinely new functional insight from metagenomic droplet screening underlines its potential to provide otherwise inaccessible annotation in the exploration of the enzyme universe.

## RESULTS

### Discovery of New β-Glucuronidases by Functional Metagenomic Screening

Our objective to discover new enzymes with β-glucuronidase activity was addressed by a functional metagenomic screening campaign, based on an ultrahigh throughput screening workflow: picoliter water-in-oil emulsion droplets were produced at kHz rates in microfluidic devices, so that a large library could be screened to identify rare events. In addition to beating the odds of potentially low hit rates, practical challenges had to be overcome: (a) background activity in lysates of the screening host (ascribed to β-glucuronidase A) led to a high number of false positive hits and limited the detection range, and (b) phenotypic variation in single cell lysate experiments made it hard to find genuine hits amongst the high number of false positives. To meet these demands, a layered screening approach was designed, as illustrated in Figure 1, suitable to identify rare catalysts even with activities similar or just above the un- or buffer-catalysed background rates. Fluorescein-di-β-glucuronide (FD-β-GlcA) was used as a bait substrate, making the detection exquisitely sensitive (requiring only 2500 product molecules per droplet).^18, 31^ Rather than using one stringent sorting step, the library size was reduced in a stepwise manner accepting to gradually remove false positive hits (e.g. arising from enzymatic rates just above background) from one layer to another to enrich genuine hits progressively, in a strategy similar to functional metagenomic campaigns in formats that did not involve droplet screening.^32, 33^.

**Figure 1:**
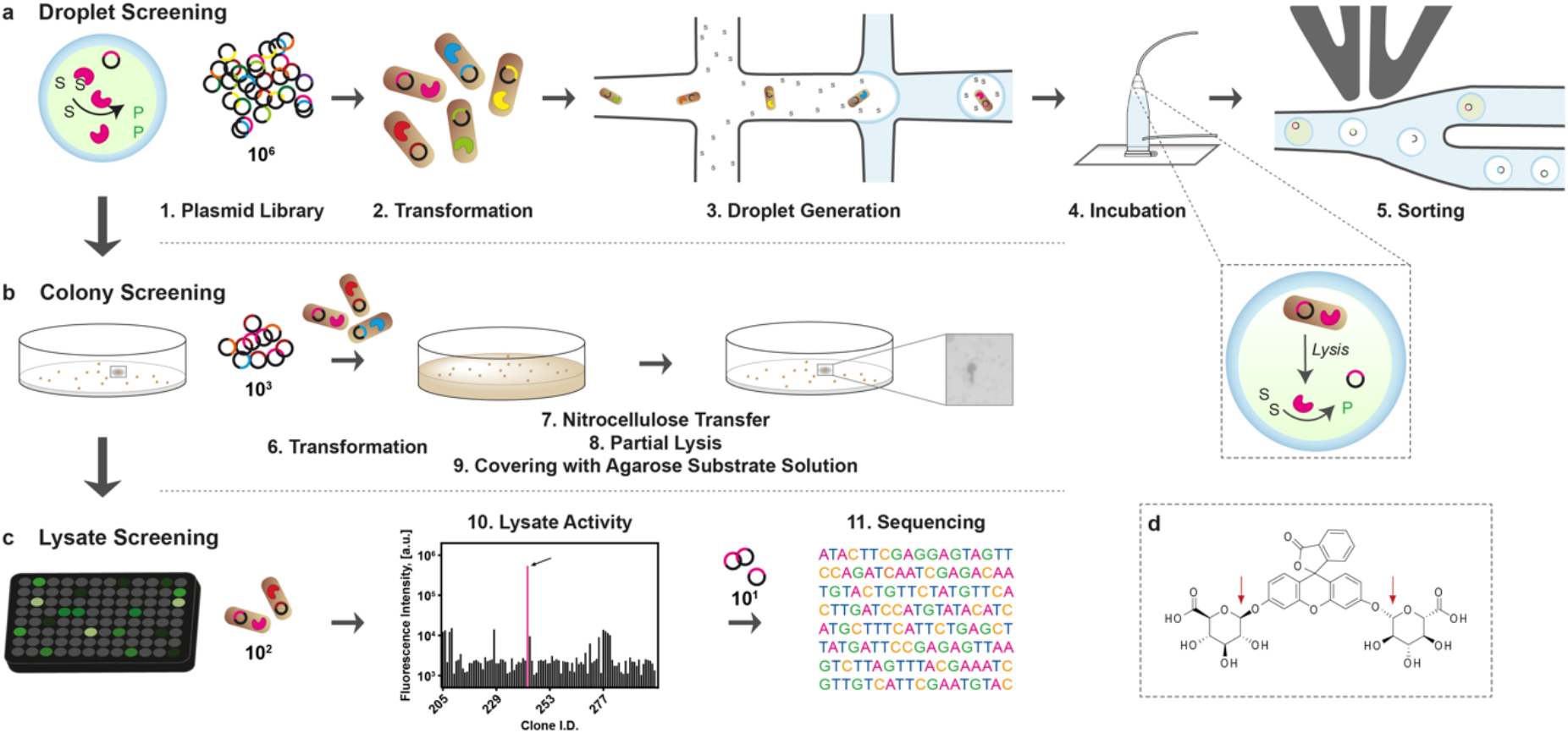
Layered workflow of a functional metagenomic screening campaign for β-glucuronidases. **(a) Droplet Screening.** The plasmid library was transformed into *E. coli* and single bacteria were encapsulated in microfluidic droplets together with lysis agent and fluorogenic substrate. During off-chip incubation lysis took place and expressed β-glucuronidases hydrolysed the substrate setting fluorescein free. Highly fluorescent droplets were sorted on chip in a FADS step. (**b) Colony Screening.** The recovered DNA was transformed into *E. coli* and plated on LB-agar. Colonies were transferred and regrown on a nitrocellulose membrane before partial lysis via cycles of freezing and thawing. To screen for colonies with β-glucuronidase activity the nitrocellulose membrane was covered with an agarose buffer solution containing substrate. Hits were detected as fluorescent spots. **(c) Lysate Screening.** Positive colonies were regrown in liquid culture and their activity was tested in lysate. Plasmids of clones with high activity were sequenced. **(d)** Structure of the substrate used in microdroplet assays (FD-GlcA). (Microfluidic chip designs are shown in Figure S8, SI.)

In the first step, *E. coli* were transformed with the ‘SCV’ library, a large metagenomic plasmid library comprising ∼10^6^ members from various environmental sources.^25, 34, 35^ Single-cell lysates of bacterial library members were screened for β-glucuronidase activity on-chip in microfluidic droplets and potential hits were isolated using fluorescence-activated droplet sorting (FADS) in two separate experiments with varying incubation time (Figure 1a). Seven million droplets with droplet occupancy λ = 0.35 (and therefore about 2.1 million occupied droplets) were screened in each selection, oversampling the library twice in each droplet screening. This lead to the identification of 697 and 645 droplets as potential hits (Figures 2a and S7, SI). Hence, approximately 0.01% of the screened droplets were selected, so that the microfluidic screening reduced the library size by three orders of magnitude to 1,432 sorted FADS hits (Table 1).

**Figure 2:**
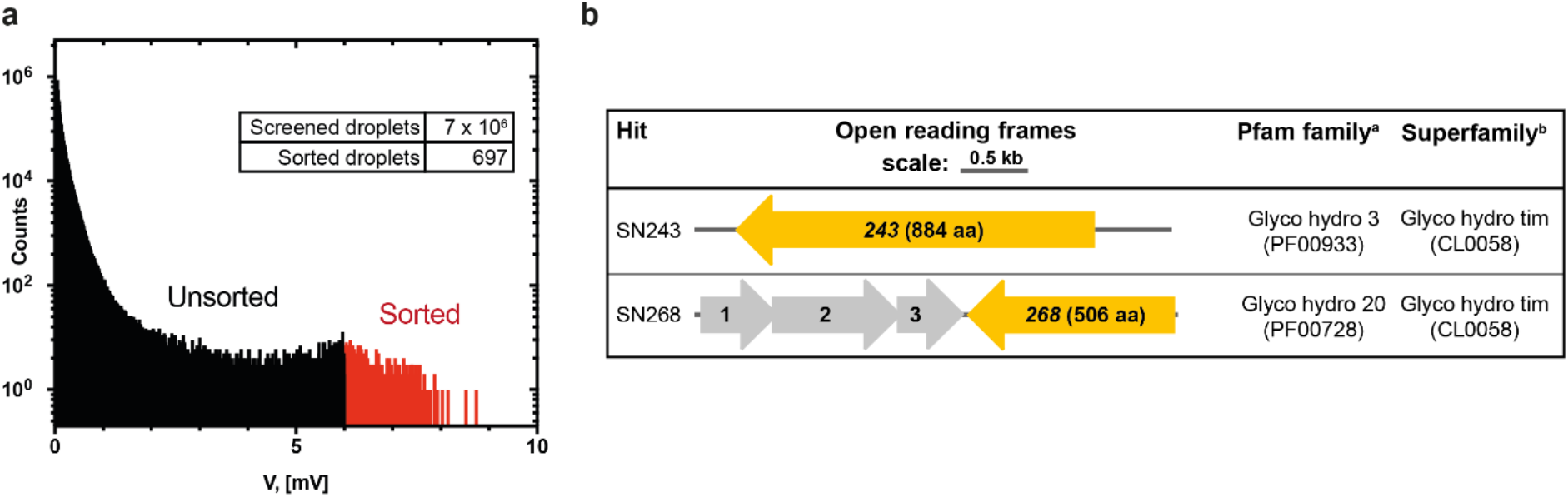
Outcome of the functional metagenomic screening campaign for β-glucuronidases. **(a) Sorting histogram of the functional metagenomic screening for β-glucuronidases in microfluidic droplets.** After an overnight incubation the droplets containing lysed cells and FD-β-GlcA were reinjected into a sorting chip and screened for high fluorescence. Droplets with a detector signal above 6 V (red) and within the chosen size range were sorted. The histogram shows the screening data on day one, when 697 droplets were selected. **(b) Metagenomic inserts harbouring β-glucuronidase hits isolated from the SCV library.** ORFs conferring β-glucuronidase activity are highlighted in orange. Other ORFs (shown in grey) have highest homology to: (1) polysaccharide deacetylases, (2) epoxyqueuosine reductases and (3) disintegrin and metalloprotease domain. Only ORFs of at least 300 bp length and with annotated homologs in the NCBI non-redundant protein sequences database were considered. *Footnotes:* ^a,b^Protein families and superfamilies were identified based on sequence homology in the Pfam database.^78^

**Table 1:** Quantification of hits discovered by the individual steps of the metagenomic screening workflow resulting in two unique and genuine hits.

| Screening step | Library size | Hits | Hit rate |
| --- | --- | --- | --- |
| 1. FADS | $1.25 \times 10^6$ | 1,342<sup>a</sup> | 0.01% |
| 2. Colony screening | 1,342<sup>b</sup> | 25 | 1.86% |
| 3. Lysate screening | 134<sup>c</sup> | 28 | 20.90% |
| 4. Sequencing | 28 | 2<sup>d</sup> (total: 4) | 7.14%<sup>d</sup> (total: 14.29%) |

<sup>a</sup>Total number of sorting events combined from both sorting days (day 1: 697, day 2: 645; Figure 2a and Figure S7, SI).
<sup>b</sup>The actual number of colonies was ~6,000., i.e. each FADS hit was represented multiple times.
<sup>c</sup>The size of the library submitted to the lysate screening (134) was larger than the number of hits (25) detected in the colony screening. Where a signal could not unambiguously be linked to a single colony in the second screening step, more clones were taken to the lysate screening to avoid losing hits.
<sup>d</sup>Number of unique sequences (2). Hit SN243 was found three times. The remaining 24 library members had only short metagenomic inserts or vector deletions.

In a second step, this enriched library was retransformed into *E. coli* and ∼6,000 colonies were screened using nitrocellulose transfer, partial lysis and coverage with FD-β-GlcA in an agarose-buffer solution. Twenty-five fluorescent spots were immediately detected by a laser scanner (see Figure S10a, SI for examples). As the reaction was faster than the solidification of the agarose, fluorescent spots were blurred and 134 colonies were taken forward for further screening.

In step three, a rescreening was performed testing the bacterial lysate of these 134 clones for β-glucuronidase activity in microtiter plates. Lysates from 28 clones showed substantial β-glucuronidase above the background (Figure S10b,c, SI).

Sequencing revealed two unique and genuine hits in this screening campaign (Figure 2b): SN243 and SN268. The remaining clones contained only short metagenomic inserts or partly deleted vector, suggesting that they had been isolated based on the host cell background activity towards FD-β-GlcA (see SI, Section 5.1).

The restriction sites used to insert the metagenomic DNA into the vector allowed the identification of the environmental DNA source, which was either soil or vanilla pod for SN243 and cow rumen for SN268. Nevertheless, the genomic origin of both hits is unknown, and their organisms of origin cannot be identified. Hit SN268 was not further characterized, because the open reading frame (ORF) encoding for a glycoside hydrolase could not be expressed in quantity (details in Section 3, SI).

### SN243 is a Highly Efficient β-Glucuronidase with Substrate Promiscuity

The metagenomic insert of hit SN243 contained one large ORF (for simplicity also named SN243), the sequence of which did not align with known β-glucuronidases. Instead, comparisons via the NCBI non-redundant protein sequence database suggest close homology to the GH3 family (the closest search hit shares a 75% identity with SN243, WP 110852811). The closest hits from database searches are *bona fide* β-glucosidases belonging to the GH3 family (with the most similar characterized bacterial GH3 in the CAZy database sharing 37% identity with SN243; β-glycosidase from *Terrabacter ginsenosidimutans*, ACZ66247.2). The CAZy database^3^ contains 43,222 entries for GH3 family members (release date 22/01/21), none of which were know to have β-glucuronidase activity at the outset of this study. To investigate the mismatch between annotation and screening outcome, we created an SN243 construct by shortening the ORF at the N- and C-termini, respectively (by removing a predicted signalling sequence and a transmembrane domain; Figure S11, SI). A codon-optimized gene coding for a construct with 821 amino acids was recombinantly expressed in *E. coli* BL21(DE3). The purified, soluble form of SN243 showed highest activity between pH 7.5 and pH 9.5 (Figure S12, SI) and was demonstrated to be thermostable (T_m_ = 63 °C). Michaelis-Menten parameters for the β-glucuronidase activity of SN243 at 25 °C towards *p*NP-β-glucuronide (*p*NP-β-GlcA) and FD-β-GlcA indicated K_M_ values of 13 and 8 µM, respectively, and second order rates of ∼10^6^-10^7^ M^-1^s^-1^ (Table 2, Figure S15, SI). To the best of our knowledge this is the β-glucuronidase with the highest substrate affinity of a wild-type enzyme (according to BRENDA^2^ and Table S4, SI). At its optimal reaction temperature of 56 °C (Figure S12c,d, SI), the first order rate for the hydrolysis of *p*NP-β-GlcA by SN243 doubles (compared to 25 °C), with a k_cat_ of 44.6 s^-1^ that places SN243 in the range of values observed for good, *bona fide* β-glucuronidases (see Table S4, SI).^36, 37^ Taken together with the substantial rate accelerations (with a second order rate enhancement of 4.5 x 10^14^; Extended Data Table 1) these data suggest that the screening workflow has been successful in identifying an efficient β-glucuronidase that would not have been recognised as such computationally: we can find no sequence similarity to proteins classified as β-glucuronidase (E.C. 3.2.1.31). Likewise, comparisons to homologous sequences in a blast search did not result in any β-glucuronidases and thus, SN243 would not have been assigned and annotated in databases as an enzyme catalysing this reaction.

**Table 2:** Comparison of Michaelis-Menten parameters determined for the glycoside hydrolase activity of SN243 towards various sugar substrates. Standard errors for Michaelis-Menten parameters are indicated with a 95% confidence level.

| Substrate | $K_M$, [mM] | $k_{cat}$, [s<sup>-1</sup>] | $k_{cat}/K_M$, [M<sup>-1</sup> s<sup>-1</sup>] |
| --- | --- | --- | --- |
| FD-β-GlcA | $8.0 \times 10^{-3} \pm 0.7 \times 10^{-3}$ | $142.0 \pm 6.1$ | $1.8 \times 10^7$ |
| pNP-β-GlcA | $13.7 \times 10^{-3} \pm 0.9 \times 10^{-3}$ | $23.1 \pm 0.2$ | $1.7 \times 10^6$ |
| pNP-β-Glc | $56.5 \pm 4.1$ | $0.7 \pm 0.03$ | $1.3 \times 10^1$ |
| pNP-β-GalA | $1.0 \pm 0.1$ | $16.6 \pm 0.7$ | $1.6 \times 10^4$ |
| pNP-β-Gal<sup>a</sup> | $38.7 \pm 4.6$ | $1.2 \times 10^{-2} \pm 1.1 \times 10^{-3}$ | $3.1 \times 10^{-1}$ |
| pNP-β-Xyl | $23.0 \pm 0.9$ | $0.4 \pm 0.007$ | $1.8 \times 10^1$ |
| pNP-α-L-Araf<sup>a</sup> | $84.5 \pm 9.3$ | $1.4 \times 10^{-2} \pm 1.4 \times 10^{-3}$ | $1.7 \times 10^{-1}$ |

<sup>a</sup>Limited solubility of the substrates in buffer prevented reliable determination of the Michaelis-Menten parameters. The indicated values ( $K_M$ and $k_{cat}$ ) for the reaction with pNP-β-Gal and pNP-α-L-Araf are only estimates resulting from measurements plotting the start of the Michaelis Menten curve.

The demonstration that many enzymes have additional, promiscuous activities^6, 7, 38^ has validated the idea that such side activities can boost adaptive evolution.^5^ To probe this versatility and evolutionary potential for SN243, its promiscuity was analysed with 17 sugar substrates, of which 5 showed measurable hydrolytic turnover (Table S3, SI).

Altering the stereochemistry in the 4-position (to *p*NP-β-galacturonide; *p*NP-β-GalA) lowered the catalytic efficiency k_cat_/K_M_ by two orders of magnitude (compared to the glucuronide substrate with the same leaving group, *p*NP-β-GlcA). Consistent with a misalignment of the 4-OH group, the reduced activity is mainly due to a lower affinity for the substrate (100-fold higher K_M_), while the first order rate constant is in a similar range.

Replacement of the 5-carboxy group with -CH_2_OH (*p*NP-β-D-glucopyranoside, *p*NP-β-Glc) or with a hydrogen (*p*NP-β-D-xylopyranoside; *p*NP-β-Xyl), lowers k_cat_/K_M_ by five orders of magnitude (compared to *p*NP-β-GlcA) as a composite effect of K_M_ (10^3^-10^4^-fold higher) and k_cat_ (10^1^-10^2^-fold lower). Very low activity was also detected towards a substrate with a change in 4-stereochemisty and 5-carboxy replacement (*p*NP-β-D-galactopyranoside’ *p*NP-β-Gal) or even a five-membered ring sugar (*p*NP-α-L-arabinofuranoside; *p*NP-α-L-Ara*f*) (Table 2 and Figure S16, SI). Despite the rate reduction in response to changes in substrate structure, the rate accelerations for promiscuous catalysis are substantial, ranging from 10^4^-10^7^ in first order rate acceleration, between 10^7^-10^9^ in second-order rate enhancement and 10^5^-10^10^ in catalytic proficiency (Figure 4 and Extended Data Table 1). The specificity of SN243 reflects the bait reaction used in the droplets screening: there is a clear preference for β-glucuronidase substrates even over substrates only changed in one sugar position. These promiscuous activities endow SN243 with versatility to give a head start to evolution of new activities, but with moderate trade-off^39, 40^ that may facilitate evolution involving intermediates with multiple activities.

### SN243 Expands the List of Known Activities for the Glycoside Hydrolase Family 3

Carbohydrate-active-enzymes are arguably the class with the most comprehensive classification in the CAZy database,^3^ which can even be used to assign function.^41^ The alignment with the protein sequences of experimentally characterised bacterial GH3 members extracted from the CAZy database (Extended Data Figure 1) indicates that SN243 contains the conserved active site residues of this family: Asp415 can be identified as the catalytic nucleophile. Indeed, the knockout mutant SN243^D415A^ was found to be completely inactive, supporting a GH3 assignment based on homology of functional active site residues. Furthermore, in a phylogenetic tree (Figure S17, SI) with the previously characterized bacterial GH3 members the closest neighbours are all N-acetylglucosaminidases (of the GH3 NagZ family) that share the conserved motif KH(F/I)PG(H/L)GX_4_**D**(S/T)**H,**^42^ with His and Asp in this motif (in bold) constituting a reactive acid/base dyad in the active site,^43^ to which an unusual degree of plasticity had been ascribed.^44^ The β-glucuronidase SN243 shares this highly conserved sequence pattern with slight modifications extending it to: KH(F/I)PG(H/L/G)GX_6_**D**(S/T/P)**H**. Asp331/His333 are very likely to act as a general acid-base dyad. The involvement of these residues was again experimentally tested with alanine mutants: variants SN243^H333A^ (k_cat_ = 0.3 s^-1^) and SN243^D331A^ (k_cat_ = 0.1 s^-1^) showed an 80- and 180-fold reduction in k_cat_, respectively (Figure S18, SI).

Taken together, the experimentally validated classification points to a clear assignment of SN243 as an N-acetylglucosaminidase. Nevertheless, when substrates representing this activity (*p*NP-β-D-glucosaminide, *p*NP-β-GlcNAc and *p*NP-β-D-galactosaminide, *p*NP-β-GalNAc) were tested with SN243, no hydrolysis of these substrates was detected (despite an assay sensitivity that could have detected a k_cat_/K_M_ >10^8^-fold lower than for *p*NP-β-GlcA; Extended Data Figure 2b and Table S3, SI). This observation exemplifies the additional value that hypothesis-free experimental screening provides: despite a ‘clean’, seemingly unambiguous homology model (Extended Data Figure 1), prediction of function is fraught with difficulty. Historically, experimental library screening campaigns aiming at functional enzyme assignment have been impractical due to low hit rates, but the throughput of droplet microfluidics makes such endeavours possible. As a result of a screen that can be conducted in a few days, β-glucuronidase activity can be added as a new genuine function to the description of the GH3 family.

### Dynamic Substrate Recognition at the Bottom of a Wide-Open Active Site Cleft

SN243 crystallised readily into well-diffracting crystals, but attempts to solve the phases via molecular replacement, selenomethionine labelling or heavy metal incubation were unsuccessful. Only sulfur phasing on the specialised beamline I23 allowed the phases to be determined and revealed its *apo*-structure at 1.87 Å resolution (PDB: 7QE1). These phases allowed the determination of a co-crystallised structure of the wild-type enzyme with the product, GlcA, at 2.15 Å (PDB: 7QE2); the inactive mutant D415A with the substrates FD-β-GlcA (PDB: 7QEA) and *p*NP-β-GlcA (PDB: 7QEF); the mutant D415N with the partially hydrolysed substrate *p*NP-β-GlcA (PDB: 7QEE) and an *apo*-structure of the mutant D415N (PDB: 7QG4). Consistent with the high sequence homology diagnostic for GH3 membership, the active site with the product GlcA (Figure 3 and Extended Data Figure 3) is lined by the conserved motif **KH**FP**G**G**G**PQELGL**D**P**H** (highly conserved residues in bold). The presence of catalytic residues within < 4Å is consistent with the classic Koshland mechanism for retaining glycosidases: involving Asp415 as a carboxylate nucleophile with assistance for leaving group departure by His333 (held in place by Asp331 and together forming a catalytic dyad; Figure 3c, d). Several hydrogen bonds (two of them short: < 3Å; between O(C4) and Lys318, and O(C3) and His319, respectively) bind the hydroxy periphery of the sugar ring. Correspondingly, the K_M_ for the enzymatic cleavage of *p*NP-β-GlcA by a SN243^H319A^ mutant is 47-fold increased to 611 μM (Figure S18, SI). These interactions mirror contacts seen in a structure of the GH3 β-N-acetylglucosaminidase from *Bacillus subtilis* (*Bs*NagZ; PDB: 3NVD), where the core sugar is surrounded by Arg57 (Arg136 in SN243), Arg191 (Arg272 in SN243, but interacting with O(C3) instead of O(C2)), Lys221 (Lys318 in SN243), and His222 (His319 in SN243).^43^ Yet in contrast to the enzyme-substrate interactions in *Bs*NagZ, SN243 Arg136 is engaged in two short H-bonds (2.6 Å) that recognise the C_6_-carboxylate on the sugar. Indeed, this functional group distinguishes *p*NP-β-GlcA from known substrates of the GH3 superfamily, consistent with a role for Arg136 in maintaining specificity, which binds in a similar position as the NxK motif in GH2 β-glucuronidases^45, 46^ (Figure S20, SI). Accordingly, mutation of Arg136 to Ala resulted in a 282-fold increase in K_M_ (3.9 mM for *p*NP-β-GlcA; Figure S18, SI). Interestingly, this Arg is conserved across the NagZ family and has been predicted to be involved in binding of natural substrates, yet similar interactions have not been observed and SN243 is the first example of a specificity conferring role of this residue.^33, 43^ Furthermore, the structure of wild-type SN243 co-crystallized with GlcA shows that the −1 binding site is occluded by the surrounding protein (Figure 3b, g), suggesting that this metagenomic hit is an *exo*-acting GH.

**Figure 3:**
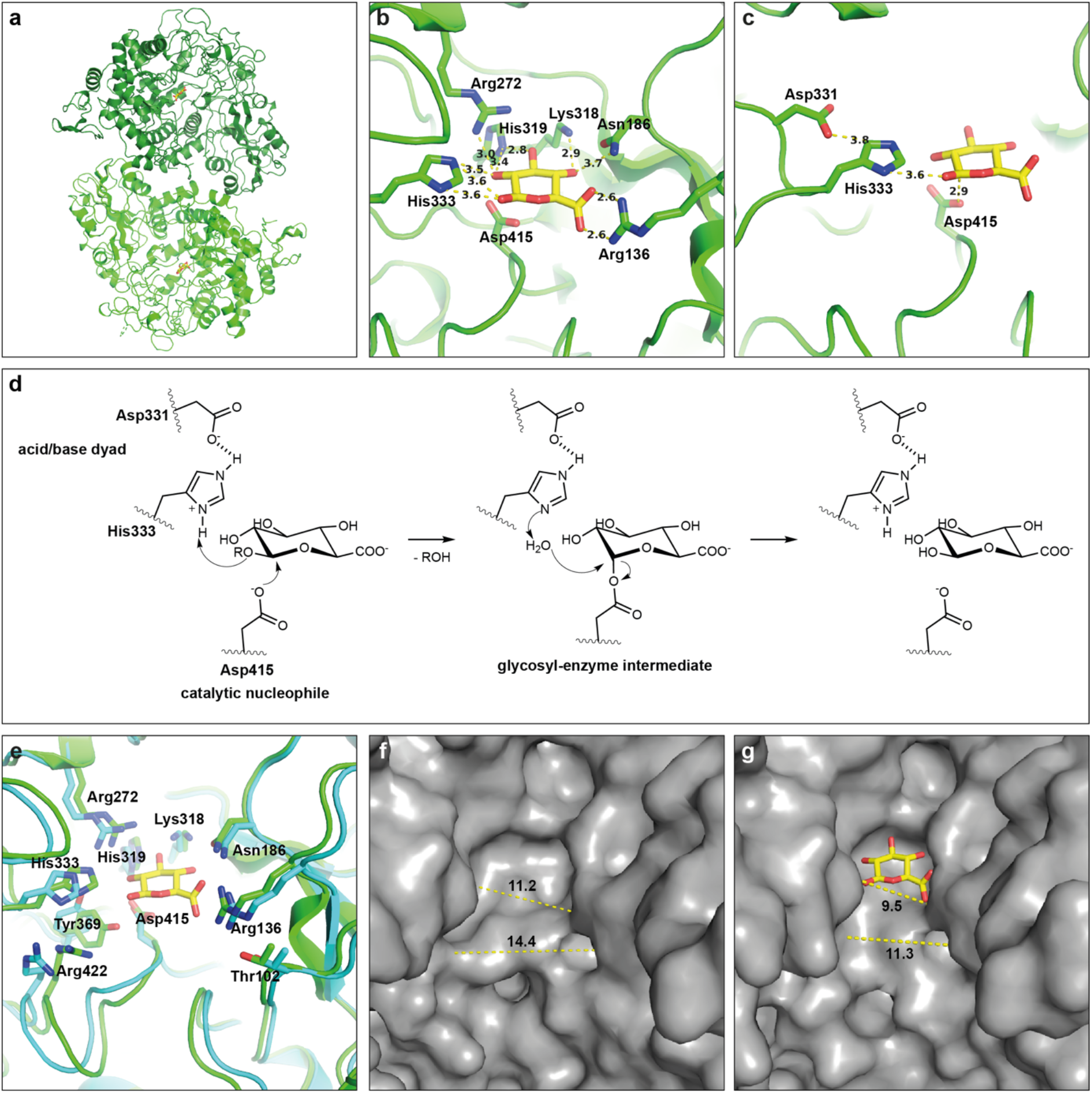
Crystal structure of SN243. **(a) Cartoon of SN243 co-crystallized with the reaction product GlcA (PDB: 7QE2).** While the *apo*-form of SN243 crystallizes as a monomer, structures with product or substrates, crystallize as dimers. **(b) Interactions of GlcA with active site residues.** The reaction product GlcA is bound in the active site via five hydrogen bonds, namely between the carboxyl group and Arg136, O(C4) and Lys 318, and O(C3) and His 319, respectively. **(c) Catalytic residues.** The catalytic nucleophile Asp415 is displayed in close proximity to the anomeric carbon of GlcA. His333 and Asp331 act as acid/base dyad. **(d) Reaction scheme.** SN243 follows a classical retaining Koshland mechanism. **(e-g) Structural rearrangement upon binding of reaction product in the active site.** In the comparative view of the cartoons **(e)** of the *apo*-crystal structure SN243 (cyan, PDB: 7QE1) and SN243 co-crystallized (green, PDB: 7QE2) with GlcA, a structural re-arrangement is observed. Arg422, Tyr369 and His333 undergo especially large shifts. Distances displayed in the wild-type *apo*-structure **(f)** and the wild-type structure co-crystallized with GlcA **(g)** are measured between side-chain atoms C2(H333) and N^*ω*^(Arg136) as well as N^*ω*^(Arg422) and O(Thr102).

Zooming out from the direct contacts with GlcA, an open cleft around the active site becomes apparent, with the small GlcA, occupying the bottom of a much larger crevice. This open cleft provides ready access to the active site of SN243, while other GH3 members possess structural elements that reach over the active site, presumably making active site access more selective. The closest structure in the PDB, a *Saccharopolyspora erythraea* glucan β-1,4-glucosidase (PDB: 5M6G), has a 20-fold smaller active site volume^47^ than SN243 (1293 *vs* 65 Å^3^) despite the structural homology in the rest of the structures (Extended Data Figure 4). We conjecture that SN243 is either intrinsically promiscuous with its tolerant, unimpeded substrate access or that a much larger oligosaccharide molecule is the natural substrate and smaller molecules can also reach the active site. In the initial *apo* and substrate-bound structures no electron density for the loop ranging from His190 to Ala202 was observed, suggesting disorder (Figure 4). Interestingly, when SN243 is modelled by AlphaFold2^48, 49^ this loop was predicted to fold over into the active site effectively reducing the volume of the binding pocket (Figure 4b, d, f). The predicted structure of the loop was consistent for all five of the models generated by AlphaFold2. However, during the screening for co-crystals of substrates with the SN243 mutant D415N a new *apo*-structure was found, with substantial electron density for the His190-Ala202 loop (despite the lack of density corresponding to the substrate) allowing it to be successfully modelled. In this crystal structure the loop folds away from the active site, leaving the active site fully open (Figure 4b, c, e). The difference to the AlphaFold2 prediction highlights the limits of an artificial intelligence (AI) analysis: based upon multiple sequence alignments the predicted position of this loop is likely to be biased by the position of similar structural elements in related proteins. Just as with predictions based on sequence homology, AI predictions can be burdened by previous evidence in the PDB. Although AlphaFold2 successfully predicts most of the structure of SN243, this loop is vital to the understanding of the protein as the position affects the substrate selectivity of SN243. There are two possible explanations for the difference between the AlphaFold2 predictions and the experimentally determined structures. First, in SN243 this loop is always folded back as in the crystal structure, giving a wide-open active site with little substrate selectivity and the AlphaFold2 prediction is wrong. Alternatively, the AlphaFold2 model and the crystal structure represent two different conformations of a mobile loop. This idea is supported by the observation that the loop is disordered in many structures and has higher normalized B-factors than the rest of the catalytic core (Figure 4a). However, the loop is well defined in the *apo*-structure of the mutant D415N. The loop may then fold in or out over the active site, depending on the whether substrates are bound to SN243.

**Figure 4:**
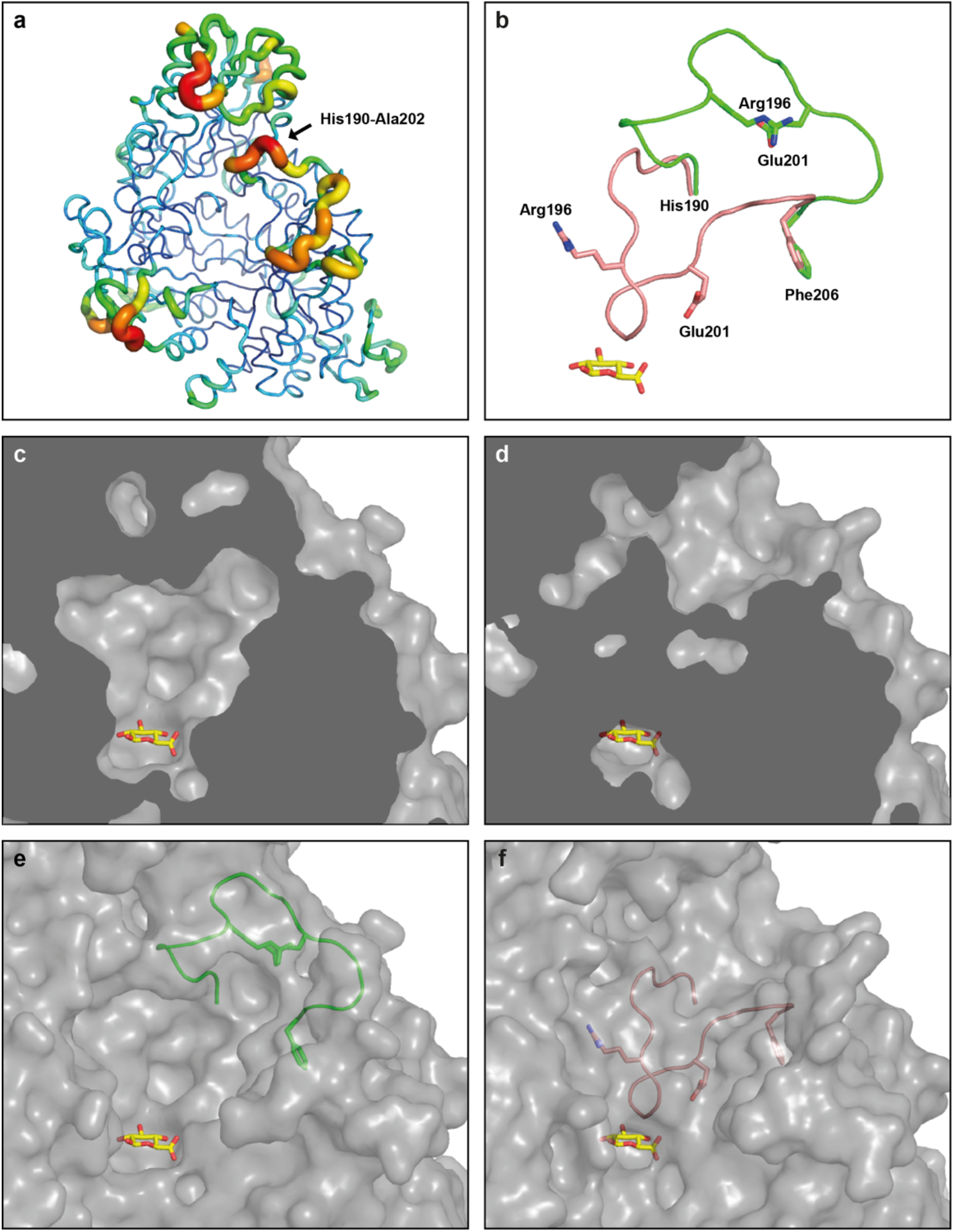
The differences between the experimentally derived crystal structure and the model generated by AlphaFold2. **(a)** The B-factors of SN243 show the disordered loop (His190-Ala202) in the original crystal structure (PDB: 7QE1). **(b)** The cartoon displays the difference in the loop His190-Ala202 between the *apo*-crystal structure of SN243 mutant D415N (green, PDB: 7QG4) and the AlphaFold2 model (salmon). The size of the active site in the experimentally derived model from the crystal structure **(c)** is much larger in comparison to the AlphaFold2 model **(d).** The position of the loop His190-Ala202 (green) in the experimentally derived model from the crystal structure **(e)** leaves the access to the active site wide-open, whereas this loop (salmon) folds over the active site in the AlphaFold2 model **(f)**. The reaction product GlcA is shown in yellow.

A comparison of crystal structures co-crystallized with substrates or product with the two *apo*-structures shows that the active site (and the cleft leading to it) narrows upon binding to a ligand (Figure 3, e-g). Comparison of the active site of the *apo*-structure to a structure co-crystallized with product GlcA point to a structural re-arrangement that leads to a tighter cleft in the co-crystallized structure. When GlcA is bound, Arg422, Tyr369 and His333 undergo large shifts and the active site access becomes narrower by more than 3 Å (from 14.4 to 11.3 Å, Figure 3). Such rearrangements are observed for all small molecules that were co-crystallized in this work (Extended Data Figure 3). While GHs are generally rigid around the active site, flexibility has been observed in the GH3 NagZ subfamily^44, 50^ and also features in GH2 β-glucuronidases,^51^ suggesting they might be intrinsic to enzymes catalysing this reaction. While in SN243 a tightening of the active site is observed upon substrate binding, Little *et al.* (2018) described a switch between an open and a closed conformation in the *Parabacteroides merdae* β-glucuronidase that is brought about by a 50 Å shift of a loop, placing a potential substrate binding residue into the active site.^52^

## DISCUSSION

Biochemical characterization of SN243 suggests that it is a genuine β-glucuronidase, devoid of the hallmark activities of GH3 and thus expanding the functional content of this family. Since SN243 was identified with a model substrate, we cannot infer the natural substrate (see Section 5.2, SI). As our metagenomic library contains only short environmental DNA inserts in a plasmid (rather than fosmid) library, the accompanying short fragments of the genomic context of its originating organism provide no additional clues. However, β-linked glucuronides are prominent in higher plants (Carbohydrate Structure Database: csdb.glycoscience.ru),^53^ consistent with SN243’s origin from a library derived from soil, where plant material is degraded. Namely, the decoration of rhamnogalacturonans,^54^ arabinogalactans of various gum species,^55, 56^ and glucuronomannan^57–59^ are well documented examples of cell wall polysaccharides bearing β-linked glucuronides. SN243 is likely to be an *exo*-acting GH, but the scope of possible natural substrates is not necessarily limited to terminal β-GlcA moieties, if the enzyme acts in a sequence of enzymes required to break down a polysaccharide. The C-terminal transmembrane domain of SN243 and its N-terminal signal peptide suggest that the enzyme may be bound to the bacterial surface, where it could target a large polysaccharide.^60, 61^ Alternatively, SN243 could be acting on oligosaccharides in the periplasmic space, which are imported following their depolymerization by membrane-bound and secreted CAZymes.

Relying on CAZy classification, overall sequence analysis and a conserved sequence motif (conserved catalytic nucleophile and the acid-base dyad of the GH3 β-GlcA NagZ subgroup), the function of SN243 would have been mis-assigned as one of the previously recorded activities. The GH3 is a large family comprising various GH activities, yet no current member had been known to be a β-glucuronidase at the outset of this study. The addition of SN243 as a β-glucuronidase to GH3 expands the catalogue of recorded activities, providing an addition of a reference point for future assignments. A discovery effort by function alone will unearth new catalysts empirically, avoiding dependence on the study design, presumed knowledge of activity type and taxonomic bias.^62^ Droplet microfluidics will be the technology of choice because its throughput allows rapid screening of much larger libraries than previously possible.^32, 63–65^

SN243 is as good as *any* wild-type β-glucuronidase on record (see Table S4, SI) but exhibits also significant substrate promiscuity with activities within a few orders of magnitude in k_cat_/K_M_ of the primary activity. First, we note that a high native activity (at least compared to known β-glucuronidases) does not preclude substantial promiscuous activities. Indeed, high intrinsic reactivity of the catalytic machinery in conjunction with a wide-open active site for easy access had been shown to allow both high promiscuity *and* high catalytic efficiency in the alkaline phosphatase superfamily.^66–68^ Second, SN243 breaks new ground in that there are no previous reports comprising all five promiscuous activities listed in Table 2 for a β-glucuronidase (Extended Data Figure 2), so the observed activities appear not to be intrinsic for an enzyme with this main activity (notwithstanding that promiscuity may not have been explored in previous studies or experimentally missed due to limits of the assay). The structural insight into SN243 highlights regions of the protein that may permit promiscuous interactions: comparisons of active site access diameter and volume (see the example shown in Extended Data Figure 4) to the closest homologous structures in the PDB show significant divergence from the previously known GH3 members and suggest fewer impediments for access of promiscuous substrates of various sizes.

Finally, the analysis of promiscuous activities is an important aspect of enzyme characterisation (even if often not recorded), because it reports on the versatility of the enzyme, suggesting, in an evolutionary context, ‘bridges’ between activities, when head-start side activities are exploited by adaptive laboratory or natural evolution^69^ for additional sugar classes as substrates.

A long-term benefit of recording multiple activities in databases (like CAZy^3^ or BRENDA^2^) is to assess the functional potential of a given enzyme and it homologs, enabling protein engineering, either rationally or by charting routes for directed evolution. While sequence information abounds and structural information may soon be accessible without experiment,^48, 70^ bridgeheads of functional annotation are the missing link to gain a comprehensive, system-wide understanding of the enzyme universe (see Section 4, SI). A network of activity correlations (as provided e.g. in Extended Data Figure 2) shows a partially reciprocal relationship of main and promiscuous activities, illustrating how network nodes^66, 71, 72^ can contribute to an understanding how function emerges.^73–75^ Even if such a characterisation is never complete, fractions of a network can be sufficient to understand it as a whole, as exemplified in the Pareto principle, embodied in the 80/20 rule.^76, 77^ The fraction necessary to understand the complex functional interrelationship in a network representation of *all* enzymes is yet to be defined, but the annotation of a ‘*vital few*’ ^76, 77^ by droplet microfluidics will undoubtedly propel such efforts.

## AUTHOR INFORMATION

### Present Addresses

^†^ Australian Synchrotron, 800 Blackburn Rd, Clayton VIC 3168, Australia.

## ACKNOWLEDGMENTS

The authors thank Philip Mair for help with FADS, D Janssen (Groningen University) for making metagenomic DNA libraries available for this work, Katherine Stott for help with CD spectra and Godwin Aleku, Simon Ladevèze and other members of the Hollfelder group for comments on the manuscript. We are grateful for access to and support of the Department of Biochemistry X-ray crystallographic facility. Data collected at Diamond Light Source on beamlines IO3, IO4 and I23 contributed to the data in this manuscript. (proposal numbers 18548 and 25402)

## Methods

### Microfluidic chip devices

Poly(dimethyl)siloxane (PDMS) chip devices for the emulsification and subsequent screening in microfluidic droplets were prepared as described previously.^79^ The corresponding chip designs (Figure S8) can be downloaded from http://openwetware.org/wiki/DropBase.

### Bacterial suspension for the microfluidic screening experiment

A pooled plasmid library, called the ‘SCV’ library,^79^ comprising ∼1.25 million environmental DNA inserts originating from cow rumen, different soils^80^ and vanilla pods^81^ with an average insert size of 3-5 kB, was used for this study.

The SCV library (20 ng) was transformed into electrocompetent *E. coli* (E. cloni 10G Elite, Agilent), plated on two lysogeny broth (LB) agar plates (14 cm diameter) supplemented with 50 mg/L kanamycin and incubated at 37 °C for 48 h. This yielded ∼10^8^ colonies, hence covering the library about 100 times. Colonies were washed off the plates with LB medium spun down at 6000 rpm for 6 min and the pellet was washed twice with 1 mL 100 mM Tris-HCl pH 8.0, 100 mM NaCl, 50 mg/L kanamycin. The bacterial library was resuspended in buffer containing 25% (v/v) Percoll to OD_600_ = 0.8. As the bacterial suspension is diluted 1:1 during droplet formation, an OD_600_ = 0.4 is expected in droplets. Assuming a Poisson distribution (P(k)= (λ^k^ *e^-λ^)/k!),^82, 83^ this corresponds to λ= 0.35 in 15 μm droplets. In this case, the actual occupancy values were: 70.5% empty, 24.7% one cell, 4.3% two cells, 0.5% three cells, <0.3% more than three cells.

### Generation of monodisperse water-in-oil emulsions

Monodisperse water-in-oil droplets were generated on flow-focusing chips with three inlets connected via PTFE tubing (0.38 mm ID, 1.09 mm OD, Portex) to glass syringes (SGE). These contained (a) 1.75% (v/v) 008-FluoroSurfactant (RAN Biotechnologies) in fluorinated oil HFE-7500, (b) 100 mM Tris-HCl pH 8.0, 100 mM NaCl, 50 μg/mL kanamycin supplemented with 0.8x CelLytic B and 30 kU/ml rLysozyme (Merck) as lysis agents, the fluorogenic substrate fluorescein-di-β-D-glucuronide (FD-β-GlcA, 10 μM), 100 nM fluorescein (as offset) and (c) the bacterial suspension. Flow control was achieved by low-pressure syringe pumps (neMESYS). The flow rates were set to 500 μL/h for oil and to 50 μL/h for the aqueous solutions, resulting in the generation of droplets with a volume of 4 pL at a rate of 6.6 kHz. The process was visually monitored under a microscope (Navitar) equipped with a camera (ALLIED Vision Technologies) and a high-speed Phantom camera (v7.2, Vision Research). A series of stills is shown in Figure S9a in the SI to illustrate droplet formation.

### Droplet incubation

Droplets were collected and incubated in a droplet container, which was self-made from an inverted Eppendorf tube connected to tubing through holes at the tip of the Eppendorf tube and on the side. The container was pre-filled with HFE-7500 supplemented with 0.5% (v/v) 008-FluoroSurfactant (Ran Biotechnologies, MA) and incubated at room temperature for one or two nights.

### Ultrahigh throughput sorting of droplets

The optical and electronic system for fluorescence-activated droplet sorting (FADS) was set up as previously described by Colin *et al.*^79^ A syringe containing 0.5% (v/v) 008-FluoroSurfactant in HFE-7500 was connected to the droplet storage device. Under the control of low-pressure syringe pumps, droplets were re-injected into a sorting chip at 15 μL/h. To create space between droplets, HFE-7500 was injected at 450 μL/h allowing droplet sorting above 1 kHz. Fluorophore excitation was achieved by focussing a 488 nm laser 80 μm upstream of the Y-shaped sorting junction under a 40x microscope objective (UPlan-FLN, Olympus). A photomultiplier tube (H8249, Hamamatsu Photonics) collected and amplified the emitted light signal. The processed signal (data acquisition card, National Instruments) was recorded by a peak detection algorithm (LabView 8.2, National Instruments). When the fluorescent peak recorded for a droplet exceeded a chosen threshold, a pulse was applied through the connected electrodes on the chip, causing the high-fluorescent droplet to exit through the narrower sorting channel instead of the default waste channel. Droplets were collected in PTFE tubing and pushed with an HFE-7500 filled syringe into a DNA LoBind tube (Eppendorf) at the end of the sorting. All sorting events were recorded by a high-speed camera, so that the quality of the sorting could be checked live and retrospectively (for an example, see Figure S9b, SI). An enrichment factor for β-glucuronidase expressing clones in this setup was determined to be 246, starting from a 1:10,000 dilution (Figure S6, SI).

### De-emulsification and DNA purification

Droplets collected from FADS were first de-emulsified to recover plasmid DNA: 200 μL 1H,1H,2H,2H-Perfluoro-1-octanol and 100 μL 2 ng/μL sheared salmon sperm DNA (ThermoFisher) were added to the droplet emulsion. After thorough vortexing and centrifugation (4000 rpm, 1 min) the aqueous supernatant was transferred to a fresh DNA LoBind tube and the procedure was repeated two more times. Subsequently, the DNA solution was purified using the DNA Clean & Concentrator-5 Kit (Zymo Research) following the manufacturer’s instructions.

### Metagenomic colony screening

DNA recovered from metagenomic screenings was transformed into *E. coli* (E. cloni 10G Elite, Agilent) and plated on 14 cm LB agar plates supplemented with kanamycin (50 μg/mL). Colonies were grown overnight at 37 °C and transferred onto a nitrocellulose membrane (BioTrace NT Pure Nitrocellulose Transfer Membrane 0.2 μm, PALL Life Sciences). Colonies on the original agar plate were regrown at 37 °C overnight. The nitrocellulose membrane was placed (colonies facing upwards) on a fresh LB agar plate supplemented with 50 μg/mL kanamycin and incubated at 37 °C overnight. Lysis was performed by freezing the membrane at −20 °C for at least 60 min followed by three cycles of heat shock (10 min at 37 °C, 10 min at −20 °C). The membrane was then transferred into an empty Petri dish and covered with agarose (0.5% w/v) in 100 mM Tris-HCl pH 8.0, 100 mM NaCl, 1 μM FD-β-GlcA. To detect active colonies, plates were immediately scanned for fluorescence using a Typhoon FLA 9500 (GE Healtcare) with a PMT of 300, an excitation wavelength λ_excitation_ of 473 nm and a 510 nm filter for detection.

### Metagenomic lysate screening in microtiter plates

Single colonies identified by their activity towards FD-β-GlcA in the colony screening assay were grown in 1 mL liquid LB cultures supplemented with kanamycin (50 μg/mL) in 96-well deep-well plates at 37 °C overnight. The OD_600_ of the cell culture suspension was measured in a 1:10 dilution using a plate reader (Tecan Infinite 200 PRO). 500 μL samples of each culture were transferred into a fresh 96-well deep well plate and centrifuged (at 4000 rpm and 4 °C for 60 min). Pellets were resuspended in lysis buffer (100 μL, 1x CelLytic B, 15 kU/mL rLysozyme (Merck), 10 U/mL benzonase (Merck), 100 mM Tris-HCl pH 8.0, 100 mM NaCl) and shaken (at 37 °C for 30 min). For the detection of activity towards the fluorogenic substrate, 10 μL lysate were combined with 90 μL substrate solution (1.1 μM FD-β-GlcA, 100 mM Tris-HCl pH 8.0, 100 mM NaCl) and the emerging fluorescence was monitored by a plate reader (λ_excitation_ = 480 nm, λ_emission_ = 530 nm). The fluorescence intensity recorded for each well was normalised to the OD_600_ of the corresponding culture. Hits were identified as such when lysate measurements satisfied the following criterion: a normalized fluorescence higher than the background *plus* 3x standard deviation (obtained from four negative controls). The plasmid DNA of hits was isolated (GeneJET Plasmid Miniprep Kit, ThermoFisher) and Sanger sequenced via primer walking. Open reading frames (ORFs) were detected by ORFfinder tool available on the NCBI website (https://www.ncbi.nlm.nih.gov/orffinder/).

### Production and purification of a truncated SN243 constructs

**Cloning primers for SN243 expression constructs.** Bases in capital letters anneal to the template.

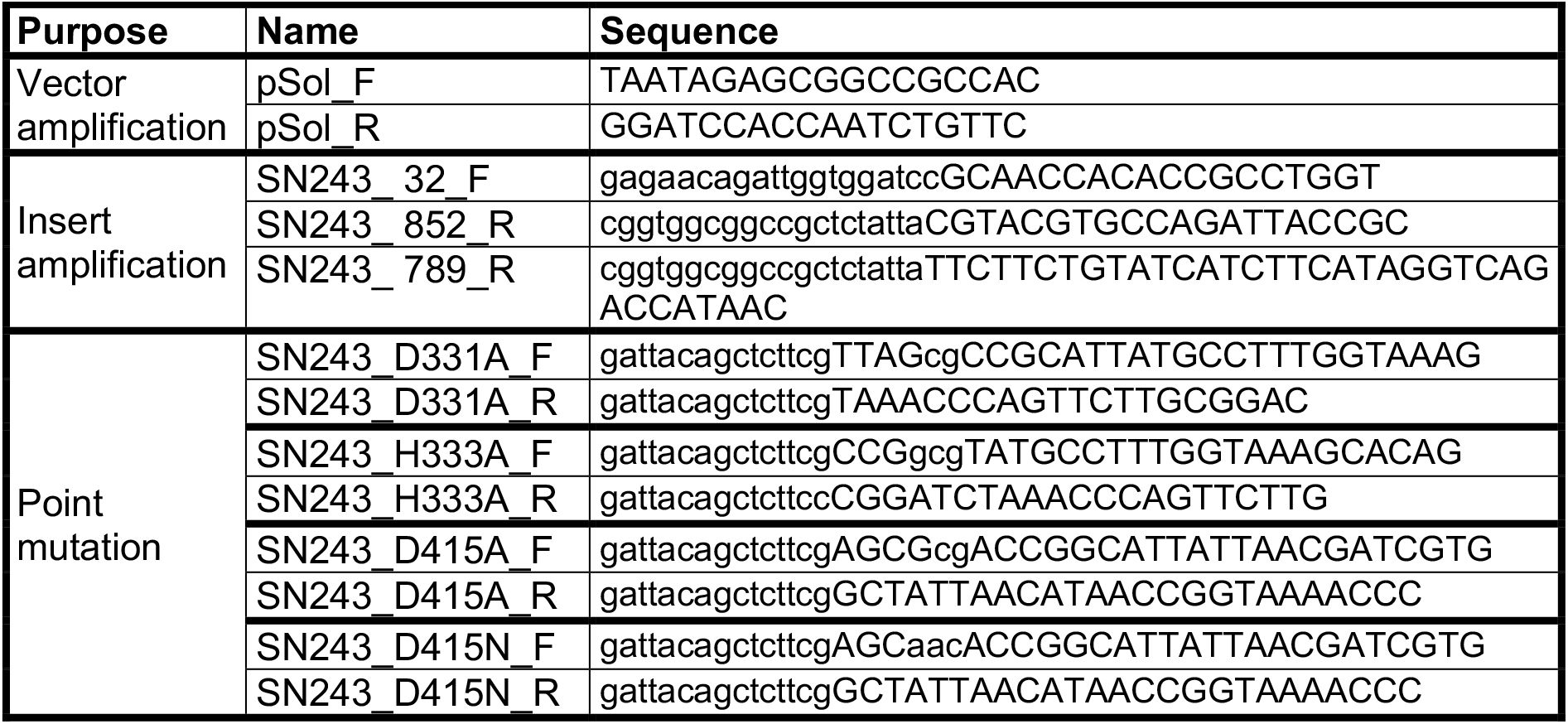

The ORF coding for SN243 (for sequences see section 2, SI) was cloned as codon-optimised synthetic gene (GeneArt Strings, ThermoFisher) using Gibson assembly^84^ into a pSol (Lucigen) expression vector (modified to include the tag from vector pC013, Addgene plasmid #90097), so that a fusion gene with N-terminal His6-Twin-Strep(II)-SUMO-tag was obtained. To remove the predicted N-terminal signalling tag and a C-terminal transmembrane domain (Figure S11, SI), SN243 constructs were truncated by 31 amino acids at the N-terminus and by 32 amino acids at the C-terminus (for activity tests and characterization) or by 95 amino acids at the C-terminus (for crystallization). Point mutations were introduced to the wild-type (wt) constructs by overlap extension PCR and SapI restriction digest. For protein expression, recombinant plasmids were transformed into *E. coli* BL21(DE3) and 1 L cultures were grown in LB with 50 mg/L kanamycin at 37 °C. Cultures were induced at OD_600_ = 0.6 with L-rhamnose (1 g/L, for 20 h at 20 °C). Bacteria were harvested by centrifugation and resuspended in buffer (100 mM Tris-HCl pH 8.0, 500 mM NaCl, 10 mM imidazole and 2.5 mM MgCl_2_). Egg white lysozyme (1 mg/mL, Merck), 1x BugBuster (Merck) and Benzonase (5 µL of the commercial solution of 2.5 kU, Merck) were added and cultures were lysed at room temperature for 30 min. Cell debris were spun down (12,000 rpm, 30 min, 4°C). IMAC purification (Ni-NTA agarose, Qiagen) was performed with the following elution buffer: 100 mM Tris-HCl pH 8.0, 500 mM NaCl, 300 mM imidazole. The protein was further purified by affinity chromatography with Strep-Tactin sepharose (IBA) equilibrated in 100 mM Tris pH 8.0, 150 mM NaCl and eluted with 2.5 mM D-desthiobiotin. After removal of D-desthiobiotin via PD-10 (Cytiva) in-house produced GST-tagged Ulp-1 was added and tag cleavage was completed overnight at 4 °C. The tag was removed via a second Strep-Tactin column and the Ulp-1 protease via affinity to SuperGlu agarose (Generon). Purified protein was concentrated to 10 mg/mL using an Amicon 50 K concentrator.

### Glycoside hydrolase substrates

**List of suppliers, from which fluorescein- and *para*-nitrophenol (*p*NP)-coupled substrates were purchased for activity tests.**

| Supplier | Substrates |
| --- | --- |
| Megazyme | pNP-β-D-galacturonide (pNP-βGalA)<br>pNP-β-D-galactoside (pNP-β-Gal)<br>pNP-β-D-mannopyranoside (pNP-β-Man)<br>pNP-β-lactoside (pNP-β-Lac)<br>pNP-α-D-galacturonide (pNP-α-GalA)<br>pNP-α-D-galactoside (pNP-α-Gal)<br>pNP-α-D-glucuronide (pNP-α-GlcA)<br>pNP-α-D-glucoside (pNP-α-Glc)<br>pNP-α-L-fucopyranoside (pNP-α-L-Fuc)<br>pNP-α-D-mannopyranoside (pNP-α-Man)<br>pNP-α-xylopyranoside (pNP-α-Xyl) |
| Merck | Fluorescein-di-β-D-glucuronide (FD-β-GlcA)<br>pNP-β-D-glucuronide (pNP-β-GlcA)<br>pNP-β-D-glucoside (pNP-β-Glc)<br>pNP-β-xylopyranoside (pNP-β-Xyl)<br>pNP-β-D-glucosaminide (pNP-β-GlcNAc)<br>pNP-α-L-rhamnopyranoside (pNP-α-L-Rha) |
| Glycosynth | pNP-α-L-arabinofuranoside (pNP-α-L-Araf) |
| Carbosynth | pNP-β-D-galactosaminide (pNP-β-GalNAc) |

### Enzyme assays

All experiments were conducted in 100 mM Tris pH 8.0, 150 mM NaCl and 100 μL final volume at 25°C unless otherwise stated. The melting temperature (T_m_) of SN243 was determined using 5 μM SN243 and 5x SyproOrange (Invitrogen) in 25 μl reaction volume by increasing the temperature (25-95 °C) in 0.5 °C steps and detecting the fluorescence signal in an RT-PCR cycler (CFX Connect, BioRad). Enzyme reactions were measured with an Infinite 200 PRO plate reader (Tecan). The activity of SN243 towards FD-β-GlcA (1 nM 243) was followed via fluorescence measurement (λ_excitation_ = 480 nm, λ_emission_ = 530 nm) and the hydrolysis of *p*NP-coupled substrates was detected via absorbance at 405 nm. Kinetic parameters were determined by fitting initial rates v_0_ to the Michaelis-Menten equation using GraphPad Prism. For analysis of the temperature dependence of the catalysed reaction, initial rates v_0_ of the hydrolysis of *p*NP-β-GlcA (50 μM) by SN243 (10 nM) were measured in steps of 2 °C (from 22 to 62 °C). The kinetics at the temperature optimum at 56 °C were measured in a Spectramax iD5 (Molecular Devices) in triplicate.

To determine the pH optimum of the β-glucuronidase reaction, v_0_ for the activity of SN243 (10 nM) towards *p*NP-β-GlcA (10 μM) was measured and normalised in various buffer conditions (100 mM citrate pH 4-6; sodium phosphate pH 6-8; Tris pH 7.5-9; glycine pH 9-10; CAPS pH 9.5-11; all buffers containing 150 mM NaCl). The IC_50_ for the product inhibition was determined with 10 μM *p*NP-β-GlcA with addition of varied D-glucuronide (GlcA) concentrations.

### Circular dichroism spectroscopy

Circular dichroism (CD) spectra were recorded on an Aviv 410 instrument in the range from 250 nm to 190 nm at 20 °C using a 1 mm quartz-cuvette. Protein samples were at 1 μM in 10 mM HEPES pH 7.4, 50 mM NaF. CD spectra were smoothened by correction for buffer noise and normalized to the number of peptide bonds in each sample expressed as mean residue ellipticity (Θ_mre_): Θ_mre_(λ) = smoothened signal [millidegrees] / ((number of amino acids – 1) * protein concentration [M] * cell pathlength [cm] * 10).

### Phylogenetic analysis

All 251 GenBank entries for characterized bacterial GH3 members were retrieved from the CAZy database (release date 22/02/2021); duplicate entries were removed and the fasta sequences for the remaining 165 proteins were retrieved from the NCBI database. SN243 was aligned with this set of sequences using ClustalW. Multiple sequence alignments were visualized with ESPript (http://espript.ibcp.fr/ESPript/cgi-bin/ESPript.cgi)^85^ and the phylogenetic tree (Figure S17) was displayed using Geneious.

### X-ray structure determination

SN243 (truncated variant: amino acids 32-789) was buffer exchanged to 100 mM MES pH 6.0, 500 mM NaCl using a PD-10 column (Cytiva) and concentrated to 10 mg/mL in an Amicon 50 K filter. Screening of crystallization conditions was performed at 20 °C in 96-well plates using the sitting-drop vapour diffusion method: drops were composed of 200 nL protein solution and 200 nL reservoir solution, with an 80 μL reservoir. Crystallisation experiments were set up using the Mosquito LV robotics system (sptlabtech) and imaged using Rockimager 1000 (formulatrix). Crystals of SN243 wt appeared after one week in 0.1 M MES pH 6.5, 0.01 M ZnSO_4_ and 25% PEG MME 550. All crystals were cryo-protected by addition of 25% glycerol. X-ray diffraction data were collected at the Diamond Light Source on beamlines IO3 and IO4. Data processing was performed using autoproc on the diamond autoprocessing pipeline.^86^

Phases were obtained via sulphur phasing of crystals grown in similar conditions (0.1 M MES pH 6.5, 0.016 ZnSO_4_ and 25% PEG MME, seeded): The crystals were harvested using LithoLoop sample mounts on specialised I23 copper sample assemblies. Data were collected on the long-wavelength beamline I23 at Diamond Light Source^87^ at an X-ray energy of 4.5 keV (λ = 2.755 Å) using the semi-cylindrical PILATUS 12M (Dectris, CH).

Sweeps of 360° with an exposure time of 0.1 s per 0.1° image were taken from three crystals grown in the same condition. As the space group was identified as *P*1, multiple datasets were collected from each crystal at varying κ and φ angles to ensure completeness. In total, 15 datasets were merged.

Data integration and scaling was performed with XDS and XSCALE^88^ in the space group *P*1 with unit cells 52.9 Å, 89.7 Å, 103.0 Å, 67.5°, 83.5°, 77.2°. The intensities were converted to amplitudes in Aimless^89^ and a resolution cut off of 2.4 Å was chosen corresponding to a CC_1/2_ of 0.68 and I/σI of 0.5. Phasing was performed using the pipeline CRANK2^90^ that built two partial SN243 molecules in the asymmetric unit. Manual model building in Coot^91^ and automated model building in Buccaneer^92^ improved the models but the experimental electron density maps quality limited the model building despite calculating 2-fold NCS average electron density maps. A higher 1.95 Å resolution dataset collected in the *P*2_1_2_1_2_1_ space group was used with one monomer of the partial SN243 model for molecular replacement in Phaser^93^. A TFZ score of ∼20 indicated a definitive solution with one molecule in the asymmetric unit, and electron density maps allowed the model to be built to completion. This model was used to solve the *apo*-structure of SN243 wt collected in the *P*1 space group via molecular replacement for comparisons with corresponding structures co-crystallized with substrates and product, which were collected in the same space group.

SN243 wt was co-crystallized with 2 mM glucuronic acid, and the mutants SN243^D415A^ and SN243^D415N^ with 1 mM *p*NP-β-GlcA or 200 μM FD-β-GlcA in 0.1 M Bis-Tris pH 7.5, 0.19-0.22 M LiSO_4_, 0.03-0.08 M ZnAcetate and 19.5-25% PEGSB. Data processing was performed on the diamond autoprocessing pipeline using autoproc^86^ and Dials^94^. Phases for holo-crystals were obtained using molecular replacement (Phaser, CCP4 suite)^95^ with the *apo-*crystal as model. One molecule was observed in the asymmetric unit of the *apo*-crystal and two molecules in the asymmetric unit of holo-crystals. Structures were iteratively refined using Refmac5 (CCP4 suite)^95^ and Coot.^91^ Ligand coordinates and restraints were generated from their SMILES strings using the Grade software package (https://www.globalphasing.com).^96^ All coordinates have been deposited to Protein Data Bank and accession numbers and data collection and refinement statistics are shown in Table S5, SI.

### Alphafold homology models

Homology models were produced using the ColabFold^97^ notebook and visualised in Pymol.

## Extended Data

**Extended Data Figure 1:**
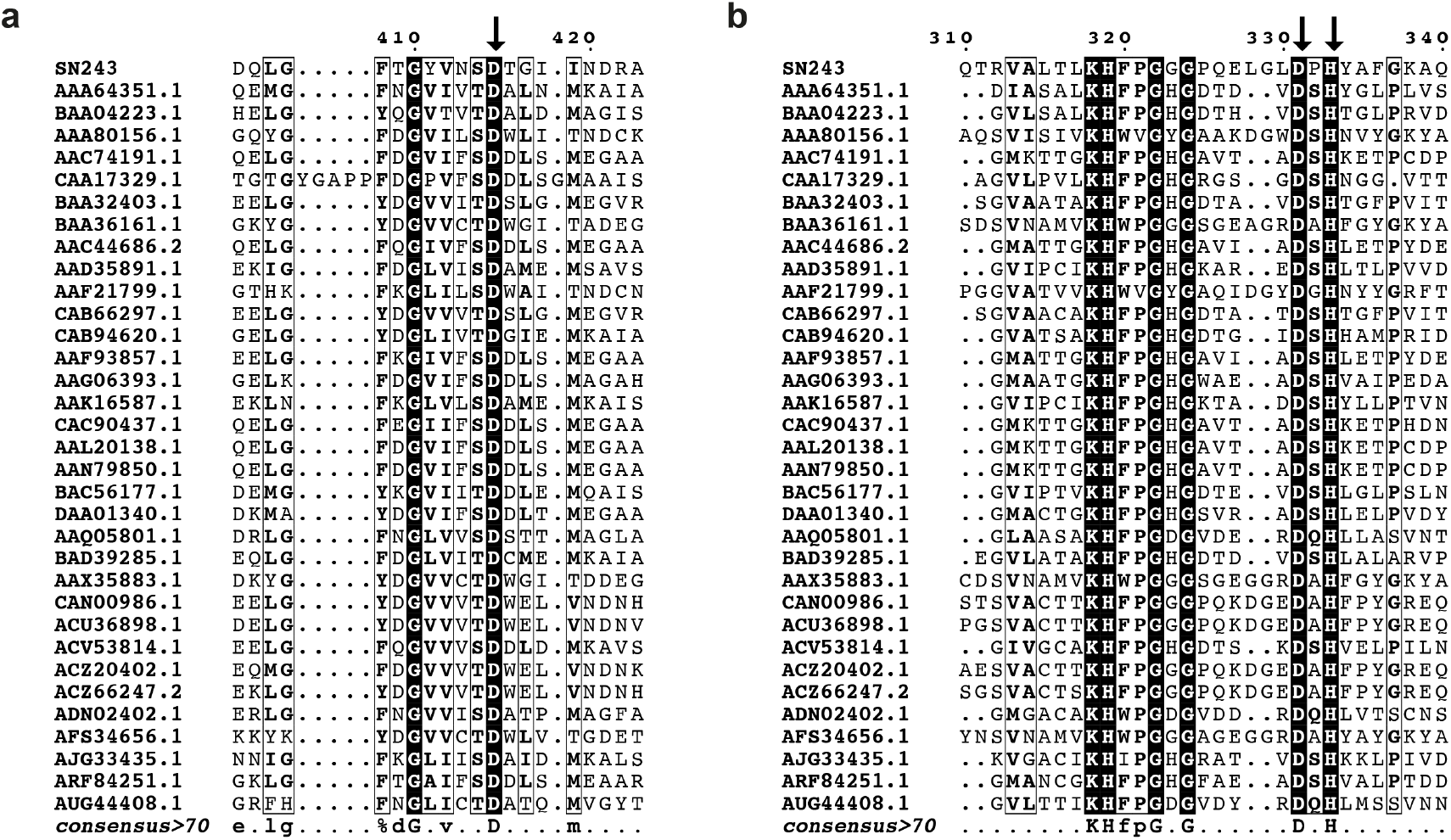
Multiple sequence alignment of SN243 with characterized bacterial GH3 family members. For this alignment the closest 33 sequences (accession codes indicated) from the phylogenetic tree generated with ClustalW were selected and aligned, and displayed with ESPript (http://espript.ibcp.fr/ESPript/cgi-bin/ESPript.cgi).^1^ Conservation is highlighted in black. A consensus marked by arrows identifies the catalytic nucleophile Asp415 **(a)** and the acid-base dyad Asp331/His333 **(b).**

**Extended Data Figure 2:**
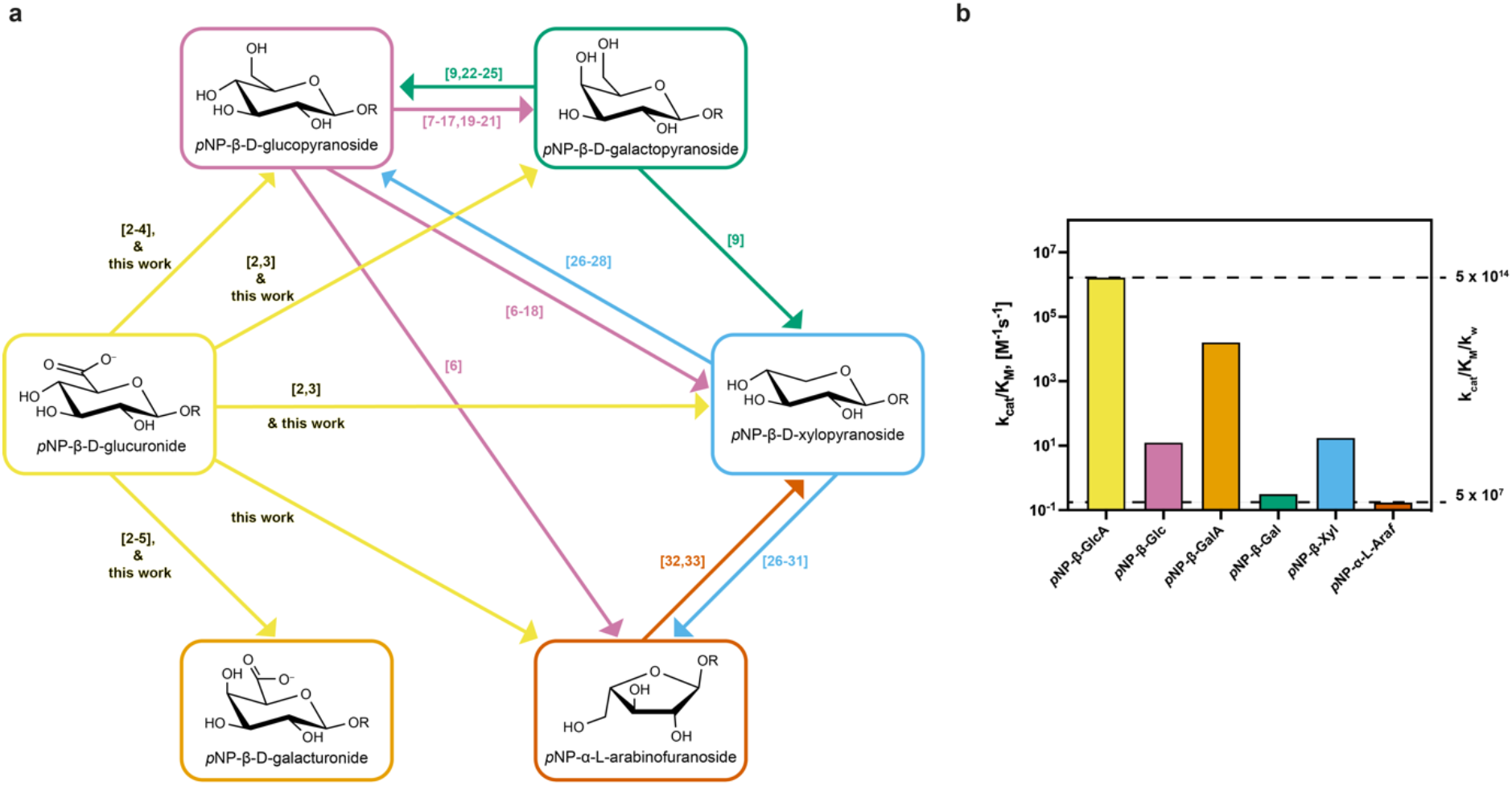
Promiscuous activities observed for SN243. **(a)** Promiscuity in glycoside hydrolase activities is still not investigated to satisfaction. Examples in the literature show that if one looks for it, side activities for other glycosides can often be found.^2–33^ Some of the promiscuous activities of SN243 had been observed before (see the literature references on the arrows), but there is no previous report of an enzyme that was active on *all* these six glycosides. Arrows start at the main activity of an enzyme and point to the observed promiscuous activity. Literature references are indicated for each arrow in square brackets. **(b)** Catalytic efficiencies (k_cat_/K_M_ – left y-axis) vary between 10^6^ M^-1^s^-1^ (*p*NP-β-GlcA) and 10^-1^ M^-1^s^-1^ (*p*NP-α-L-Ara*f*), which corresponds to second order rate enhancements (right y-axis) in the range of 10^14^ and 10^7^, respectively. Detailed kinetic information can be found in Table 2 and Extended Data Table 1.

**Extended Data Figure 3:**
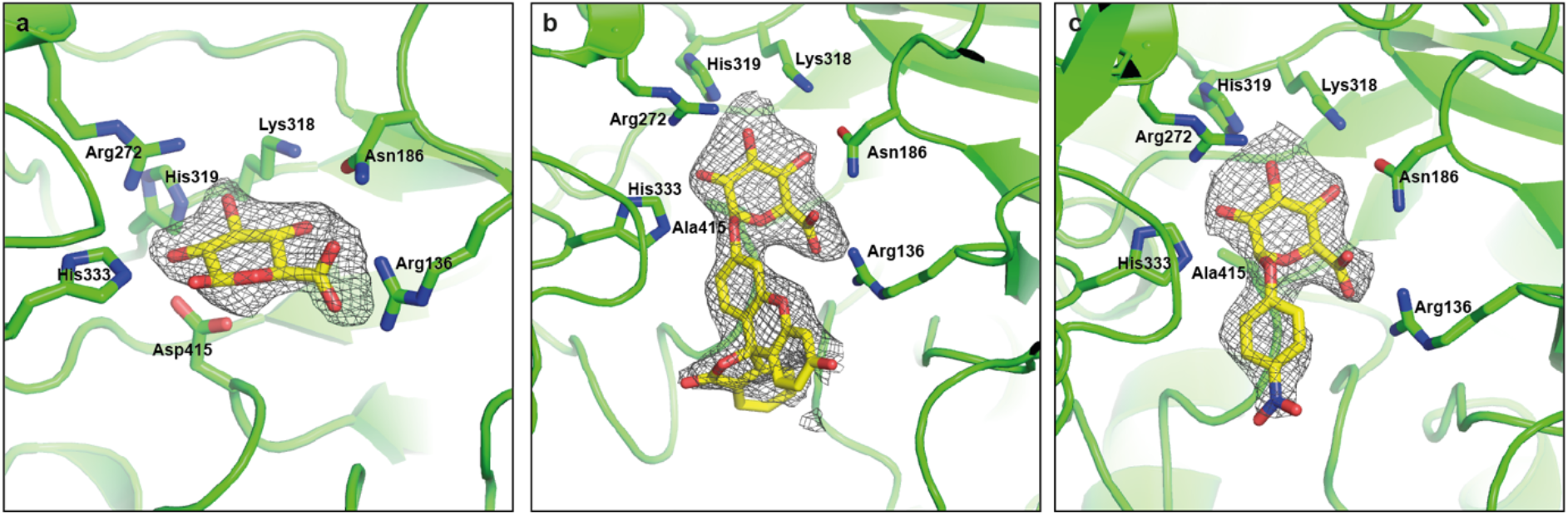
Active site of SN243 variants co-crystallized with reaction substrates and product. The structures of SN243 wt co-crystallized with GlcA (PDB: 7QE2) **(a)** and of the inactive mutant SN243^D415A^ with FD-β-GlcA (PDB: 7QEA) **(b)** and *p*NP-β-GlcA (PDB: 7QEF) **(c)** bound in the active site were obtained at 2.15 Å, 2.28 Å and 2.41 Å, respectively. The grey mesh shows the Fo-Fc map for GlcA **(a)**, FD-β-GlcA **(b)** and *p*NP-β-GlcA **(c)** contoured at 2 σ. As no electron density was observed for the second GlcA moiety of the FD-β-GlcA ligand in (b), the monoglycosylated substrate was modelled.

**Extended Data Figure 4:**
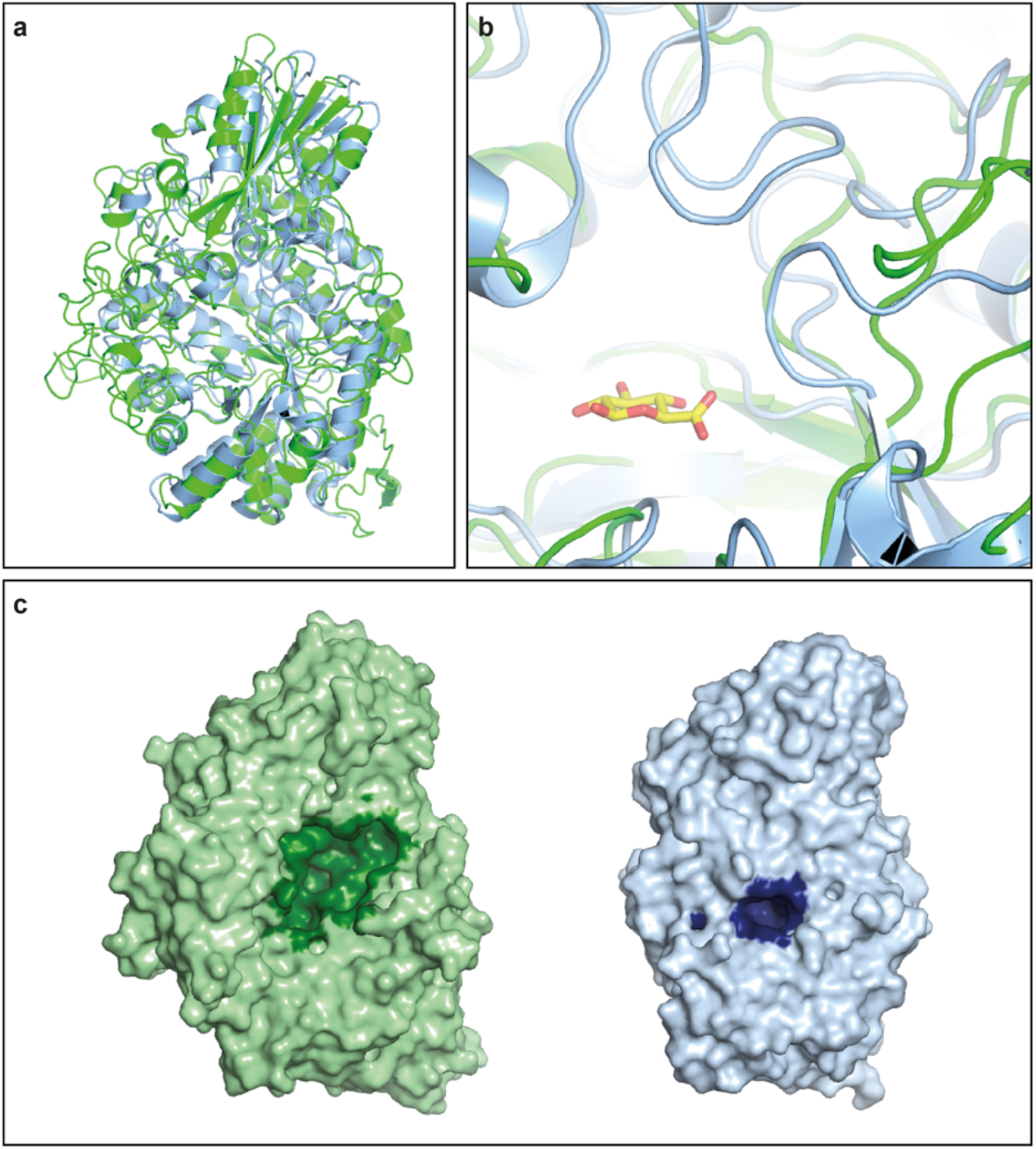
Comparison of the active site access between SN243 and the structure of a homologous GH3 member. **(a)** Cartoon representation of the structures of SN243 (crystal structure with density for loop His190-Ala202, PDB: 7QG4, green) and *Saccharopolyspora erythraea* glucan β-1,4-glucosidase (PDB: 5M6G, blue). **(b)** The zoom into the active site with the reaction product GlcA (yellow) shows loops folding over the binding pocket in the glucan β-1,4-glucosidase (blue), making the active site much smaller than in SN243 (green). **(c)** Surface display of SN243 (green) and the glucan β-1,4-glucosidase (blue). Active site binding pockets have been calculated by Castp 3.0^34^ and are highlighted. The active site volume of SN243 is with 1293 Å^3^ about twenty times as large as the active site volume (65 Å^3^) of the homologous GH3 structure.

## Extended Tables

**Extended Table 1:** Rate accelerations of the hydrolytic cleavage of *p*NP-glycosides by SN243.

| Substrate | $k_{\text{cat}}/k_{\text{uncat}}^b$ | $k_{\text{cat}}/K_M/k_{\text{uncat}}^c, [\text{M}^{-1}]$ | $k_{\text{cat}}/K_M/k_w^d$ |
| --- | --- | --- | --- |
| *p*NP-β-GlcA | $1.2 \times 10^8$ | $8.4 \times 10^{12}$ | $4.5 \times 10^{14}$ |
| *p*NP-β-Glc<sup>a</sup> | $3.7 \times 10^6$ | $6.5 \times 10^7$ | $3.5 \times 10^9$ |
| *p*NP-β-GalA | $8.3 \times 10^7$ | $8.0 \times 10^{10}$ | $4.3 \times 10^{12}$ |
| *p*NP-β-Gal | $6.0 \times 10^4$ | $1.6 \times 10^6$ | $8.4 \times 10^7$ |
| *p*NP-β-Xyl | $2.1 \times 10^6$ | $9.1 \times 10^7$ | $4.9 \times 10^9$ |
| *p*NP-α-L-Araf | $7.2 \times 10^4$ | $8.5 \times 10^5$ | $4.6 \times 10^7$ |

<sup>a</sup>The rate for the uncatalyzed hydrolysis of *p*NP-β-Glc was determined at 30 °C and 50 mM CAPS buffer, pH 10.5 as $k_{\text{uncat}} = 2.0 \times 10^{-7} \text{ s}^{-1}$ with $k_{\text{uncat}} = k_w \times c(\text{H}_2\text{O})$ and $c(\text{H}_2\text{O}) = 55 \text{ M}$ .<sup>35</sup> Assuming similar values at room temperature and pH 8 for different *p*NP-glycosides, $k_{\text{uncat}}$ was approximated as $\sim 2.0 \times 10^{-7} \text{ s}^{-1}$ and used for all substrates. Given the typical variation in the reactivity of *p*NP sugars, this should be sufficient as an order-of-magnitude estimate. As $\alpha$ -linked *p*NP-glycosides tend to have slightly lower uncatalyzed hydrolysis rates than their β-linked anomer analogs, and as the $k_{\text{uncat}}$ is higher for *p*NP-β-Glc than for *p*NP-β-Gal, using the $k_{\text{uncat}}$ of *p*NP-β-Glc for all *p*NP-glycoside substrates may thus under- (rather than over-) estimate the reaction enhancements. The data of Wolfenden et al. does not comprise a *p*NP-β-Glc substrate, but comparisons using related substrates would give similar rate enhancements.<sup>36</sup>
<sup>b</sup>First-order rate acceleration.
<sup>c</sup>Catalytic proficiencies.
<sup>d</sup>Second-order rate enhancement.

## References

1 Mitchell, A. L. et al. MGnify: the microbiome analysis resource in 2020. Nucleic Acids Res 48, D570–D578, doi:10.1093/nar/gkz1035 (2020).

2 Chang, A. et al. BRENDA, the ELIXIR core data resource in 2021: new developments and updates. Nucleic Acids Res 49, D498–D508, doi:10.1093/nar/gkaa1025 (2021).

3 Lombard, V., Golaconda Ramulu, H., Drula, E., Coutinho, P. M. & Henrissat, B. The carbohydrate-active enzymes database (CAZy) in 2013. Nucleic Acids Res 42, D490–495, doi:10.1093/nar/gkt1178 (2014).

4 Consortium, T. U. UniProt: the universal protein knowledgebase in 2021. Nucleic Acids Research 49, D480–D489, doi:10.1093/nar/gkaa1100 (2020).

5 Jensen, R. A. Enzyme recruitment in evolution of new function. Annu Rev Microbiol 30, 409–425, doi:10.1146/annurev.mi.30.100176.002205 (1976).

6 O’Brien, P. J. & Herschlag, D. Catalytic promiscuity and the evolution of new enzymatic activities. Chem Biol 6, R91–R105, doi:10.1016/S1074-5521(99)80033-7 (1999).

7 Khersonsky, O. & Tawfik, D. S. Enzyme promiscuity: a mechanistic and evolutionary perspective. Annu Rev Biochem 79, 471–505, doi:10.1146/annurev-biochem-030409-143718 (2010).

8 Miton, C., Jonas, S., Hyvonen, M., Tokuriki, N. & Hollfelder, F. Exploring Catalytic Promiscuity in the Alkaline Phosphatase Superfamily by Directed Evolution. Protein Science 21, 135–135 (2012).

9 Jonas, S. & Hollfelder, F. in The Handbook of Protein Engineering Vol. 1 (eds U. T. Bornscheuer & S. Lutz) 47–72 (Wiley-VCH, 2008).

10 Handelsman, J. Metagenomics: application of genomics to uncultured microorganisms. Microbiol Mol Biol Rev 68, 669–685, doi:10.1128/MMBR.68.4.669-685.2004 (2004).

11 Robinson, S. L., Piel, J. & Sunagawa, S. A roadmap for metagenomic enzyme discovery. Natural Product Reports 38, 1994–2023, doi:10.1039/D1NP00006C (2021).

12 Lorenz, P. & Eck, J. Metagenomics and industrial applications. Nat Rev Microbiol 3, 510–516, doi:10.1038/nrmicro1161 (2005).

13 Colin, P. Y., Zinchenko, A. & Hollfelder, F. Enzyme engineering in biomimetic compartments. Curr Opin Struct Biol 33, 42–51, doi:10.1016/j.sbi.2015.06.001 (2015).

14 Mair, P., Gielen, F. & Hollfelder, F. Exploring sequence space in search of functional enzymes using microfluidic droplets. Curr Opin Chem Biol 37, 137–144, doi:10.1016/j.cbpa.2017.02.018 (2017).

15 Leemhuis, H., Stein, V., Griffiths, A. D. & Hollfelder, F. New genotype-phenotype linkages for directed evolution of functional proteins. Current Opinion in Structural Biology 15, 472–478, doi:Doi 10.1016/J.Sbi.2005.07.006 (2005).

16 Schaerli, Y. & Hollfelder, F. The potential of microfluidic water-in-oil droplets in experimental biology. Mol Bio Sys 5, 1392–1404 (2009).

17 Ferrer, M., Beloqui, A., Timmis, K. N. & Golyshin, P. N. Metagenomics for mining new genetic resources of microbial communities. J Mol Microbiol Biotechnol 16, 109–123, doi:10.1159/000142898 (2009).

18 Neun, S., Zurek, P. J., Kaminski, T. S. & Hollfelder, F. Ultrahigh throughput screening for enzyme function in droplets. Methods Enzymol 643, 317–343, doi:10.1016/bs.mie.2020.06.002 (2020).

19 Stucki, A., Vallapurackal, J., Ward, T. R. & Dittrich, P. S. Droplet Microfluidics and Directed Evolution of Enzymes: An Intertwined Journey. Angewandte Chemie International Edition 60, 24368–24387, doi:https://doi.org/10.1002/anie.202016154 (2021).

20 Baret, J. C. et al. Fluorescence-activated droplet sorting (FADS): efficient microfluidic cell sorting based on enzymatic activity. Lab Chip 9, 1850–1858, doi:10.1039/b902504a (2009).

21 Gielen, F. et al. Ultrahigh-throughput-directed enzyme evolution by absorbance-activated droplet sorting (AADS). Proc Natl Acad Sci U S A 113, E7383–E7389, doi:10.1073/pnas.1606927113 (2016).

22 Zinchenko, A. et al. One in a Million: Flow Cytometric Sorting of Single Cell-Lysate Assays in Monodisperse Picolitre Double Emulsion Droplets for Directed Evolution. Anal Chem 86, 2526–2533, doi:10.1021/ac403585p (2014).

23 Tauzin, A. S. et al. Investigating host-microbiome interactions by droplet based microfluidics. Microbiome 8, 141, doi:10.1186/s40168-020-00911-z (2020).

24 Kintses, B. et al. Picoliter cell lysate assays in microfluidic droplet compartments for directed enzyme evolution. Chem Biol 19, 1001–1009, doi:S1074-5521(12)00216-5 [pii] 10.1016/j.chembiol.2012.06.009 (2012).

25 Colin, P. Y. et al. Ultrahigh-throughput discovery of promiscuous enzymes by picodroplet functional metagenomics. Nat Commun 6, 10008, doi:10.1038/ncomms10008 (2015).

26 Carnachan, S. M., Bell, T. J., Hinkley, S. F. R. & Sims, I. M. Polysaccharides from New Zealand Native Plants: A Review of Their Structure, Properties, and Potential Applications. Plants 8, 163 (2019).

27 Whitfield, C., Wear, S. S. & Sande, C. Assembly of Bacterial Capsular Polysaccharides and Exopolysaccharides. Annual Review of Microbiology 74, 521–543, doi:10.1146/annurev-micro-011420-075607 (2020).

28 Esko, J. D., Kimata, K. & Lindahl, U. Proteoglycans and sulfated glycosaminoglycans. Essentials of Glycobiology. 2nd edition (2009).

29 Dutton, G. J. Acceptor substrates of UDP glucuronosyltransferase and their assay. Glucuronidation of drugs and other compounds, 69–78 (1980).

30 Wenzl, P., Wong, L., Kwang-won, K. & Jefferson, R. A. A Functional Screen Identifies Lateral Transfer of β-Glucuronidase (gus) from Bacteria to Fungi. Molecular Biology and Evolution 22, 308–316, doi:10.1093/molbev/msi018 (2004).

31 Neun, S., Kaminski, T. S. & Hollfelder, F. in Enzyme Activity in Single Cells Vol. 628 Methods in Enzymology (eds N. L. Allbritton & M. L. Kovarik) 95–112 (Academic Press Ltd-Elsevier Science Ltd, 2019).

32 Macdonald, S. S. et al. Development and Application of a High-Throughput Functional Metagenomic Screen for Glycoside Phosphorylases. Cell Chemical Biology 26, 1001–1012.e1005, doi:10.1016/j.chembiol.2019.03.017 (2019).

33 Macdonald, S. S. et al. Structural and mechanistic analysis of a &#x3b2;-glycoside phosphorylase identified by screening a metagenomic library. Journal of Biological Chemistry 293, 3451–3467, doi:10.1074/jbc.RA117.000948 (2018).

34 Gabor, E. M., de Vries, E. J. & Janssen, D. B. Efficient recovery of environmental DNA for expression cloning by indirect extraction methods. FEMS Microbiol Ecol 44, 153–163, doi:10.1016/S0168-6496(02)00462-2 (2003).

35 Gabor, E. M., de Vries, E. J. & Janssen, D. B. Construction, characterization, and use of small-insert gene banks of DNA isolated from soil and enrichment cultures for the recovery of novel amidases. Environ Microbiol 6, 948–958, doi:10.1111/j.1462-2920.2004.00643.x (2004).

36 Wallace, B. D. et al. Structure and Inhibition of Microbiome beta-Glucuronidases Essential to the Alleviation of Cancer Drug Toxicity. Chem Biol 22, 1238–1249, doi:10.1016/j.chembiol.2015.08.005 (2015).

37 Biernat, K. A. et al. Structure, function, and inhibition of drug reactivating human gut microbial β-glucuronidases. Scientific Reports 9, 825, doi:10.1038/s41598-018-36069-w (2019).

38 Babtie, A. C., Tokuriki, N. & Hollfelder, F. What makes an enzyme promiscuous? Curr. Op. Chem. Biol 14, 200–207, doi:doi:10.1016/j.cbpa.2009.11.028 (2010).

39 Aharoni, A. et al. The ‘evolvability’ of promiscuous protein functions. Nat Genet 37, 73–76, doi:10.1038/ng1482 (2005).

40 Kaltenbach, M., Emond, S., Hollfelder, F. & Tokuriki, N. Functional Trade-Offs in Promiscuous Enzymes Cannot Be Explained by Intrinsic Mutational Robustness of the Native Activity. PLoS Genet 12, e1006305, doi:10.1371/journal.pgen.1006305 (2016).

41 Helbert, W. et al. Discovery of novel carbohydrate-active enzymes through the rational exploration of the protein sequences space. Proc Natl Acad Sci U S A 116, 6063–6068, doi:10.1073/pnas.1815791116 (2019).

42 Vocadlo, D. J., Mayer, C., He, S. & Withers, S. G. Mechanism of action and identification of Asp242 as the catalytic nucleophile of Vibrio furnisii N-acetyl-beta-D-glucosaminidase using 2-acetamido-2-deoxy-5-fluoro-alpha-L-idopyranosyl fluoride. Biochemistry 39, 117–126, doi:10.1021/bi991958d (2000).

43 Litzinger, S. et al. Structural and kinetic analysis of Bacillus subtilis N-acetylglucosaminidase reveals a unique Asp-His dyad mechanism. J Biol Chem 285, 35675–35684, doi:10.1074/jbc.M110.131037 (2010).

44 Bacik, J. P., Whitworth, G. E., Stubbs, K. A., Vocadlo, D. J. & Mark, B. L. Active site plasticity within the glycoside hydrolase NagZ underlies a dynamic mechanism of substrate distortion. Chem Biol 19, 1471–1482, doi:10.1016/j.chembiol.2012.09.016 (2012).

45 Pollet, R. M. et al. An Atlas of β-Glucuronidases in the Human Intestinal Microbiome. Structure 25, 967–977.e965, doi:https://doi.org/10.1016/j.str.2017.05.003 (2017).

46 Pellock, S. J. et al. Three structurally and functionally distinct β-glucuronidases from the human gut microbe Bacteroides uniformis. Journal of Biological Chemistry 293, 18559–18573, doi:https://doi.org/10.1074/jbc.RA118.005414 (2018).

47 Tian, W., Chen, C., Lei, X., Zhao, J. & Liang, J. CASTp 3.0: computed atlas of surface topography of proteins. Nucleic Acids Research 46, W363–W367, doi:10.1093/nar/gky473 (2018).

48 Jumper, J. et al. Highly accurate protein structure prediction with AlphaFold. Nature, doi:10.1038/s41586-021-03819-2 (2021).

49 Mirdita, M., Ovchinnikov, S. & Steinegger, M. ColabFold - Making protein folding accessible to all. bioRxiv, 2021.2008.2015.456425, doi:10.1101/2021.08.15.456425 (2021).

50 Vadlamani, G. et al. Conformational flexibility of the glycosidase NagZ allows it to bind structurally diverse inhibitors to suppress beta-lactam antibiotic resistance. Protein Sci 26, 1161–1170, doi:10.1002/pro.3166 (2017).

51 Dashnyam, P. et al. β-Glucuronidases of opportunistic bacteria are the major contributors to xenobiotic-induced toxicity in the gut. Scientific Reports 8, 16372, doi:10.1038/s41598-018-34678-z (2018).

52 Little, M. S. et al. Active site flexibility revealed in crystal structures of Parabacteroides merdae β-glucuronidase from the human gut microbiome. Protein Science 27, 2010–2022, doi:https://doi.org/10.1002/pro.3507 (2018).

53 Egorova, K. S. & Toukach, P. V. Expansion of coverage of Carbohydrate Structure Database (CSDB). Carbohydr Res 389, 112–114, doi:10.1016/j.carres.2013.10.009 (2014).

54 Renard, C. M., Crepeau, M. J. & Thibault, J. F. Glucuronic acid directly linked to galacturonic acid in the rhamnogalacturonan backbone of beet pectins. Eur J Biochem 266, 566–574, doi:10.1046/j.1432-1327.1999.00896.x (1999).

55 Fincher, G. B., Stone, B. A. & Clarke, A. E. Arabinogalactan-Proteins: Structure, Biosynthesis, and Function. Ann Rev PLan Physiol 34, 47–70, doi:https://doi.org/10.1146/annurev.pp.34.060183.000403 (1983).

56 Cartmell, A. et al. A surface endogalactanase in Bacteroides thetaiotaomicron confers keystone status for arabinogalactan degradation. Nat Microbiol 3, 1314–1326, doi:10.1038/s41564-018-0258-8 (2018).

57 Wee, M. S. M., Matia-Merino, L., Carnachan, S. M., Sims, I. M. & Goh, K. K. T. Structure of a shear-thickening polysaccharide extracted from the New Zealand black tree fern, Cyathea medullaris. International Journal of Biological Macromolecules 70, 86–91, doi:10.1016/j.ijbiomac.2014.06.032 (2014).

58 Wagner, R. et al. A high-viscosity glycoglucuronomannan from the gum exudate of Vochysia thyrsoidea: Comparison with those of other Vochysia spp. Carbohydrate Polymers 72, 382–389, doi:10.1016/j.carbpol.2007.09.005 (2008).

59 Mori, M. & Katō, K. An arabinoglucuronomannan from suspension-cultured cells of Nicotiana tabacum. Carbohydrate Research 91, 49–58, doi:https://doi.org/10.1016/S0008-6215(00)80990-8 (1981).

60 Larsbrink, J. et al. A complex gene locus enables xyloglucan utilization in the model saprophyte Cellvibrio japonicus. Molecular Microbiology 94, 418–433, doi:https://doi.org/10.1111/mmi.12776 (2014).

61 Grondin, J. M. et al. Polysaccharide Utilization Loci: Fueling Microbial Communities. Journal of Bacteriology 199, e00860–00816, doi:doi:10.1128/JB.00860-16 (2017).

62 Vickers, C. J., Fraga, D. & Patrick, W. M. Quantifying the taxonomic bias in enzymology. Protein Sci 30, 914–921, doi:10.1002/pro.4041 (2021).

63 Chuzel, L., Ganatra, M. B., Rapp, E., Henrissat, B. & Taron, C. H. Functional metagenomics identifies an exosialidase with an inverting catalytic mechanism that defines a new glycoside hydrolase family (GH156). Journal of Biological Chemistry 293, 18138–18150, doi:10.1074/jbc.RA118.003302 (2018).

64 Cheng, J. et al. Functional metagenomics reveals novel β-galactosidases not predictable from gene sequences. PLOS ONE 12, e0172545, doi:10.1371/journal.pone.0172545 (2017).

65 Armstrong, Z. et al. High-Throughput Recovery and Characterization of Metagenome-Derived Glycoside Hydrolase-Containing Clones as a Resource for Biocatalyst Development. mSystems 4, e00082–00019, doi:10.1128/mSystems.00082-19.

66 van Loo, B. et al. An efficient, multiply promiscuous hydrolase in the alkaline phosphatase superfamily. Proc. Natl Acad. Sci USA 107, 2740–2745, doi:doi/10.1073/pnas.0903951107 (2010).

67 Babtie, A. C., Bandyopadhyay, S., Olguin, L. F. & Hollfelder, F. Efficient Catalytic Promiscuity for Chemically Distinct Reactions. Angewandte Chemie-International Edition 48, 3692–3694, doi:Doi 10.1002/Anie.200805843 (2009).

68 van Loo, B. et al. Balancing Specificity and Promiscuity in Enzyme Evolution: Multidimensional Activity Transitions in the Alkaline Phosphatase Superfamily. J Am Chem Soc 141, 370–387, doi:10.1021/jacs.8b10290 (2019).

69 Copley, S. D. Evolution of new enzymes by gene duplication and divergence. FEBS J 287, 1262–1283, doi:10.1111/febs.15299 (2020).

70 Baek, M. et al. Accurate prediction of protein structures and interactions using a three-track neural network. Science, doi:10.1126/science.abj8754 (2021).

71 Roodveldt, C. & Tawfik, D. S. Shared promiscuous activities and evolutionary features in various members of the amidohydrolase superfamily. Biochemistry 44, 12728–12736, doi:10.1021/bi051021e (2005).

72 Glasner, M. E. et al. Evolution of structure and function in the o-succinylbenzoate synthase/N-acylamino acid racemase family of the enolase superfamily. J Mol Biol 360, 228–250, doi:10.1016/j.jmb.2006.04.055 (2006).

73 Furnham, N., Dawson, N. L., Rahman, S. A., Thornton, J. M. & Orengo, C. A. Large-Scale Analysis Exploring Evolution of Catalytic Machineries and Mechanisms in Enzyme Superfamilies. J Mol Biol 428, 253–267, doi:10.1016/j.jmb.2015.11.010 (2016).

74 Tyzack, J. D., Furnham, N., Sillitoe, I., Orengo, C. M. & Thornton, J. M. Understanding enzyme function evolution from a computational perspective. Curr Opin Struct Biol 47, 131–139, doi:10.1016/j.sbi.2017.08.003 (2017).

75 Tyzack, J. D., Furnham, N., Sillitoe, I., Orengo, C. M. & Thornton, J. M. Exploring Enzyme Evolution from Changes in Sequence, Structure, and Function. Methods Mol Biol 1851, 263–275, doi:10.1007/978-1-4939-8736-8_14 (2019).

76 Lappe, M. & Holm, L. Unraveling protein interaction networks with near-optimal efficiency. Nat Biotechnol 22, 98–103, doi:10.1038/nbt921 (2004).

77 Sullivan, W. Rockets, gauges, and pendulums: applying engineering principles to cell biology. Mol Biol Cell 30, 1635–1640, doi:10.1091/mbc.E19-02-0100 (2019).

78 Mistry, J. et al. Pfam: The protein families database in 2021. Nucleic Acids Research 49, D412–D419, doi:10.1093/nar/gkaa913 (2021).

## Methods References

79 Colin, P. Y. et al. Ultrahigh-throughput discovery of promiscuous enzymes by picodroplet functional metagenomics. Nat Commun 6, 10008, doi:10.1038/ncomms10008 (2015).

80 Gabor, E. M., de Vries, E. J. & Janssen, D. B. Construction, characterization, and use of small-insert gene banks of DNA isolated from soil and enrichment cultures for the recovery of novel amidases. Environ Microbiol 6, 948–958, doi:10.1111/j.1462-2920.2004.00643.x (2004).

81 Gabor, E. M., de Vries, E. J. & Janssen, D. B. Efficient recovery of environmental DNA for expression cloning by indirect extraction methods. FEMS Microbiol Ecol 44, 153–163, doi:10.1016/S0168-6496(02)00462-2 (2003).

82 Huebner, A. et al. Quantitative detection of protein expression in single cells using droplet microfluidics. Chemical Communications, 1218–1220, doi:Doi 10.1039/B618570c (2007).

83 Koster, S. et al. Drop-based microfluidic devices for encapsulation of single cells. Lab Chip 8, 1110–1115, doi:10.1039/b802941e (2008).

84 Gibson, D. G. et al. Enzymatic assembly of DNA molecules up to several hundred kilobases. Nat Methods 6, 343–345, doi:10.1038/nmeth.1318 (2009).

85 Robert, X. & Gouet, P. Deciphering key features in protein structures with the new ENDscript server. Nucleic Acids Res 42, W320–324, doi:10.1093/nar/gku316 (2014).

86 Vonrhein, C. et al. Data processing and analysis with the autoPROC toolbox. Acta Crystallographica Section D 67, 293–302, doi:doi:10.1107/S0907444911007773 (2011).

87 Wagner, A., Duman, R., Henderson, K. & Mykhaylyk, V. In-vacuum long-wavelength macromolecular crystallography. Acta Crystallographica Section D 72, 430–439, doi:doi:10.1107/S2059798316001078 (2016).

88 Kabsch, W. XDS. Acta Crystallographica Section D 66, 125–132, doi:doi:10.1107/S0907444909047337 (2010).

89 Evans, P. R. & Murshudov, G. N. How good are my data and what is the resolution? Acta Crystallographica Section D 69, 1204–1214, doi:doi:10.1107/S0907444913000061 (2013).

90 Skubak, P. et al. A new MR-SAD algorithm for the automatic building of protein models from low-resolution X-ray data and a poor starting model. IUCrJ 5, 166–171, doi:doi:10.1107/S2052252517017961 (2018).

91 Emsley, P., Lohkamp, B., Scott, W. G. & Cowtan, K. Features and development of Coot. Acta Crystallographica Section D 66, 486–501, doi:doi:10.1107/S0907444910007493 (2010).

92 Cowtan, K. The Buccaneer software for automated model building. 1. Tracing protein chains. Acta Crystallographica Section D 62, 1002–1011, doi:doi:10.1107/S0907444906022116 (2006).

93 McCoy, A. J., et al. Phaser crystallographic software. Journal of Applied Crystallography 40, 658–674 (2007).

94 Beilsten-Edmands, J. et al. Scaling diffraction data in the DIALS software package: algorithms and new approaches for multi-crystal scaling. Acta Crystallographica Section D 76, 385–399, doi:doi:10.1107/S2059798320003198 (2020).

95 Winn, M. D. et al. Overview of the CCP4 suite and current developments. Acta Crystallographica Section D 67, 235–242, doi:doi:10.1107/S0907444910045749 (2011).

96 Smart, O. S., et al. grade, version 1.2.20. (2021).

97 Mirdita, M., Ovchinnikov, S. & Steinegger, M. ColabFold - Making protein folding accessible to all. bioRxiv, 2021.2008.2015.456425, doi:10.1101/2021.08.15.456425 (2021).

## Extended Data References

1 Robert, X. & Gouet, P. Deciphering key features in protein structures with the new ENDscript server. Nucleic Acids Res 42, W320–324, doi:10.1093/nar/gku316 (2014).

2 Geddie, M. L. & Matsumura, I. Rapid Evolution of β-Glucuronidase Specificity by Saturation Mutagenesis of an Active Site Loop*. Journal of Biological Chemistry 279, 26462–26468, doi:https://doi.org/10.1074/jbc.M401447200 (2004).

3 Matsumura, I. & Ellington, A. D. In vitro Evolution of Beta-glucuronidase into a Beta-galactosidase Proceeds Through Non-specific Intermediates. Journal of Molecular Biology 305, 331–339, doi:https://doi.org/10.1006/jmbi.2000.4259 (2001).

4 Salleh, H. M. et al. Cloning and characterization of Thermotoga maritima β-glucuronidase. Carbohydrate Research 341, 49–59, doi:https://doi.org/10.1016/j.carres.2005.10.005 (2006).

5 Pellock, S. J., Walton, W. G. & Redinbo, M. R. Selecting a Single Stereocenter: The Molecular Nuances That Differentiate β-Hexuronidases in the Human Gut Microbiome. Biochemistry 58, 1311–1317, doi:10.1021/acs.biochem.8b01285 (2019).

6 Kim, B.-J., Singh, S. P. & Hayashi, K. Characteristics of chimeric enzymes constructed between Thermotoga maritima and Agrobacterium tumefaciens β-glucosidases: Role of C-terminal domain in catalytic activity. Enzyme and Microbial Technology 38, 952–959, doi:https://doi.org/10.1016/j.enzmictec.2005.08.038 (2006).

7 Nam, E. S., Kim, M. S., Lee, H. B. & Ahn, J. K. β-Glycosidase of Thermus thermophilus KNOUC202: Gene and biochemical properties of the enzyme expressed in Escherichia coli. Applied Biochemistry and Microbiology 46, 515–524, doi:10.1134/S0003683810050091 (2010).

8 Takase, M. & Horikoshi, K. Purification and Properties of a β-Glucosidase from Thermus sp. Z-1. Agricultural and Biological Chemistry 53, 559–560, doi:10.1080/00021369.1989.10869310 (1989).

9 Matsuzawa, T. & Yaoi, K. Screening, identification, and characterization of a novel saccharide-stimulated β-glycosidase from a soil metagenomic library. Applied Microbiology and Biotechnology 101, 633–646, doi:10.1007/s00253-016-7803-2 (2017).

10 Akram, F., Haq, I. u. & Mukhtar, H. Gene cloning, characterization and thermodynamic analysis of a novel multidomain hyperthermophilic GH family 3 β-glucosidase (TnBglB) from Thermotoga naphthophila RKU-10T. Process Biochemistry 66, 70–81, doi:https://doi.org/10.1016/j.procbio.2017.12.007 (2018).

11 Haq, I. U. et al. Cloning, characterization and molecular docking of a highly thermostable β-1,4-glucosidase from Thermotoga petrophila. Biotechnology Letters 34, 1703–1709, doi:10.1007/s10529-012-0953-0 (2012).

12 Hong, M.-R. et al. Characterization of a recombinant β-glucosidase from the thermophilic bacterium Caldicellulosiruptor saccharolyticus. Journal of Bioscience and Bioengineering 108, 36–40, doi:https://doi.org/10.1016/j.jbiosc.2009.02.014 (2009).

13 Nunoura, N. et al. Cloning and Nucleotide Sequence of the β-d-Glucosidase Gene from Bifidobacterium breve clb, and Expression of β-d-Glucosidase Activity in Escherichia coli. Bioscience, Biotechnology, and Biochemistry 60, 2011–2018, doi:10.1271/bbb.60.2011 (1996).

14 Plant, A. R., Oliver, J. E., Patchett, M. L., Daniel, R. M. & Morgan, H. W. Stability and substrate specificity of a β-glucosidase from the thermophilic bacterium Tp8 cloned into Escherichia coli. Archives of Biochemistry and Biophysics 262, 181–188, doi:https://doi.org/10.1016/0003-9861(88)90180-4 (1988).

15 Singh, A. & Hayashi, K. Construction of Chimeric β-Glucosidases with Improved Enzymatic Properties (∗). Journal of Biological Chemistry 270, 21928–21933, doi:https://doi.org/10.1074/jbc.270.37.21928 (1995).

16 Vallmitjana, M. et al. Mechanism of the Family 1 β-Glucosidase from Streptomyces sp: Catalytic Residues and Kinetic Studies. Biochemistry 40, 5975–5982, doi:10.1021/bi002947j (2001).

17 Michlmayr, H., Schümann, C., Barreira Braz da Silva, N. M., Kulbe, K. D. & Del Hierro, A. M. Isolation and basic characterization of a β-glucosidase from a strain of Lactobacillus brevis isolated from a malolactic starter culture. Journal of Applied Microbiology 108, 550–559, doi:https://doi.org/10.1111/j.1365-2672.2009.04461.x (2010).

18 Mattéotti, C. et al. Characterization of a new β-glucosidase/β-xylosidase from the gut microbiota of the termite (Reticulitermes santonensis). FEMS Microbiology Letters 314, 147–157, doi:10.1111/j.1574-6968.2010.02161.x (2011).

19 Kim, H.-J., Park, A.-R., Lee, J.-K. & Oh, D.-K. Characterization of an acid-labile, thermostable β-glycosidase from Thermoplasma acidophilum. Biotechnology Letters 31, 1457, doi:10.1007/s10529-009-0018-1 (2009).

20 Matsuzawa, T. et al. Crystal structure and identification of a key amino acid for glucose tolerance, substrate specificity, and transglycosylation activity of metagenomic β-glucosidase Td2F2. The FEBS Journal 283, 2340–2353, doi:https://doi.org/10.1111/febs.13743 (2016).

21 Kaper, T. et al. Comparative Structural Analysis and Substrate Specificity Engineering of the Hyperthermostable β-Glucosidase CelB from Pyrococcus furiosus. Biochemistry 39, 4963–4970, doi:10.1021/bi992463r (2000).

22 Dion, M., Fourage, L., Hallet, J.-N. & Colas, B. Cloning and expression of a β-glycosidase gene from Thermus thermophilus. Sequence and biochemical characterization of the encoded enzyme. Glycoconjugate Journal 16, 27–37, doi:10.1023/A:1006997602727 (1999).

23 Benešová, E., Lipovová, P., Dvořáková, H. & Králová, B. β-d-Galactosidase from Paenibacillus thiaminolyticus catalyzing transfucosylation reactions. Glycobiology 20, 442–451, doi:10.1093/glycob/cwp196 (2009).

24 Parikh, M. R. & Matsumura, I. Site-saturation Mutagenesis is more Efficient than DNA Shuffling for the Directed Evolution of β-Fucosidase from β-Galactosidase. Journal of Molecular Biology 352, 621–628, doi:https://doi.org/10.1016/j.jmb.2005.07.020 (2005).

25 Zhang, J.-H., Dawes, G. & Stemmer, W. P. C. Directed evolution of a fucosidase from a galactosidase by DNA shuffling and screening. Proceedings of the National Academy of Sciences 94, 4504–4509, doi:10.1073/pnas.94.9.4504 (1997).

26 Vocadlo, D. J., Wicki, J., Rupitz, K. & Withers, S. G. Mechanism of Thermoanaerobacterium saccharolyticum β-Xylosidase: Kinetic Studies. Biochemistry 41, 9727–9735, doi:10.1021/bi020077v (2002).

27 Zhang, S. et al. Cloning, overexpression and characterization of a thermostable β-xylosidase from Thermotoga petrophila and cooperated transformation of ginsenoside extract to ginsenoside 20(S)-Rg3 with a β-glucosidase. Bioorganic Chemistry 85, 159–167, doi:https://doi.org/10.1016/j.bioorg.2018.12.026 (2019).

28 Yin, Y.-R. et al. Expression and characterisation of a pH and salt tolerant, thermostable and xylose tolerant recombinant GH43 β-xylosidase from Thermobifida halotolerans YIM 90462T for promoting hemicellulose degradation. Antonie van Leeuwenhoek 112, 339–350, doi:10.1007/s10482-018-1161-2 (2019).

29 Shao, W. & Wiegel, J. Purification and characterization of a thermostable beta-xylosidase from Thermoanaerobacter ethanolicus. Journal of Bacteriology 174, 5848–5853, doi:doi:10.1128/jb.174.18.5848-5853.1992 (1992).

30 Jordan, D. B. & Braker, J. D. β-d-Xylosidase from Selenomonas ruminantium: Role of Glutamate 186 in Catalysis Revealed by Site-Directed Mutagenesis, Alternate Substrates, and Active-Site Inhibitor. Applied Biochemistry and Biotechnology 161, 395–410, doi:10.1007/s12010-009-8874-7 (2010).

31 Wagschal, K. et al. Purification and Characterization of a Glycoside Hydrolase Family 43 β-xylosidase from Geobacillus thermoleovorans IT-08. Applied Biochemistry and Biotechnology 155, 1–10, doi:10.1007/s12010-008-8362-5 (2009).

32 Bouraoui, H. et al. The GH51 α-l-arabinofuranosidase from Paenibacillus sp. THS1 is multifunctional, hydrolyzing main-chain and side-chain glycosidic bonds in heteroxylans. Biotechnology for Biofuels 9, 140, doi:10.1186/s13068-016-0550-x (2016).

33 Shallom, D. et al. The identification of the acid–base catalyst of α-arabinofuranosidase from Geobacillus stearothermophilus T-6, a family 51 glycoside hydrolase. FEBS Letters 514, 163–167, doi:https://doi.org/10.1016/S0014-5793(02)02343-8 (2002).

34 Tian, W., Chen, C., Lei, X., Zhao, J. & Liang, J. CASTp 3.0: computed atlas of surface topography of proteins. Nucleic Acids Research 46, W363–W367, doi:10.1093/nar/gky473 (2018).

35 Striegler, S., Fan, Q. H. & Rath, N. P. Binuclear copper(II) complexes discriminating epimeric glycosides and alpha- and beta-glycosidic bonds in aqueous solution. J Catal 338, 349–364, doi:10.1016/j.jcat.2015.12.026 (2016).

36 Wolfenden, R. Benchmark Reaction Rates, the Stability of Biological Molecules in Water, and the Evolution of Catalytic Power in Enzymes. Annual Review of Biochemistry 80, 645–667, doi:10.1146/annurev-biochem-060409-093051 (2011).

