## Supplementary Information for "Functional metagenomic screening in microfluidic droplets identifies a β-glucuronidase in an unprecedented sequence neighbourhood"

#### Table of Contents

1. Nucleotide and amino acid sequences for hit SN243 ...................................... S2
2. Nucleotide and amino acid sequences for hit SN268 ...................................... S5
3. Evidence for activity of hit SN268 ...................................... S7
4. Bioinformatic exploration of the sequence neighbourhood of SN243 ............. S11
5. Supplementary Notes ...................................... S12
   5.1 False positive sorting hits ...................................... S12
   5.2 Activity tests of SN243 with natural carbohydrate substrates.................... S12
6. Supporting Figures ...................................... S14
7. Supporting Tables ...................................... S29
8. Supporting References ...................................... S33

#### 1. Nucleotide and amino acid sequences for hit SN243

##### Genomic insert of library member SN243

Highlighted: ORF SN243

AACCTCACCACGCTGCTCGAGGCACCCTCGGTCGCCGAGGGGCTCAAGGAGAGCCAGGAGGGGTG
GCGGGCGGCCGTCTCCCACGCCGTCGCCAACGGCGTCCCGGTGCCCGGGTTCTCGGCGGCGCTGG
CCTACTACGACACCGTGCGCGCCGAGCGCCTGCCCGCCGCCCTGGTCCAGGGCCAGCGCGACTTCT
TCGGCGCCCACACCTACCACCGCGTGGACCGCGAGGGCGTCTTCCACACCGAGTGGTCCGGCGACC
GCAGCGAGACCGAGCAGGGCTGACCCCGGCGCCACCTCGCTCCAGAACCCGGCTCCCCGTCCGCG
GGGAGCCGGGCTCTCGCCGTCCGGGGGTACCCGTGTCCGTGCGATAGCACGGTCCGGCGCCTCCG
TGCGCTCCGGTGGCAACCGACGGCAACTTCTTCGGTCGTGACCCGCGACACGGCCAGCATCTGGCT
CACCGGCCCCCGCCACGGTGACGCGGCGCGGAGGAACCGGTCGAGGCACGTGCACGTGCACGTGC
CGCGGACGACCTGCGTCAGCGACGACGACGATCGAGGAGGAAGAATGTCACGTACCCCACGTGTGG
GGCTGCGCACCGCGGTGGCGACCGCCGCCGCTGCGCCACTGGTGCTGACGCTCGCCGTCGGCGCC
GGGGCGGCGACCACGCCGCCCGGTGACCTCGAGCAGCCCGAGCTCGAGGCTCGGGTGAAGGAGAT
CATCGAGGTCGACGGGTACCAGTTCCGCGACCTCAACGACAACGGCGAGCTGGACCCGTACGAGGA
CTGGCGCCTGCCGACCCCCGAGCGGGTCGCCGACCTCGTCGGGCAGATGTCGCTGGTCGAGAAGTC
GGGCCTCATGCTCATCAACACGCTCAACGCGGCGTGCGACCCGCAGACCGGCGAGTTCGGGGTGCT
GCCCGCGCAGGCGGACAACTACATCAACACCCAGCACATGCACCGGTTCGTCCAGCGCCGTTTCGC
CCGCACCCAGGGCTAGCTTCCGCAACGTCGTCGACGTGCGTGCGGAGGGGGTGGAGTGCACCGGG
ACGGGGACCCCCGTGGTGAGCCCGGCCGAGGCCGCGACGTTCACCAACGCCGTGCAGGAGATGAG
CGAGGCCACCCGCCTGGGGATCCCCTCGCTGTTCAAGTCCAACGCGCGCAACCACATCGACCCCGA
CGCGCGGGTCGGCATCAACGAGGCCGCCGGTGCGTTCAGCGCCTTCCCGAAGGAGGCGGGCATCG
CGGCGGCCGCGCTGGGTGAGCAGGCGCGCCGCACCGGCGAGGCCACCACGGGTGACATGTCCGTC
GTCGCGGACTTCGCCGACGTCATGGGCGAGGAGTGGGCCTCGATCGGCCTGCGCGGCATGTACGG
CTACATGGCCGACCTGTCCACCGAGCCGCGCTGGTACCGCACCCACGAGACCTTCACCGAGGACGC
CTACCTCGCCGCCGAGATCATGGAGACCCTCGTCCAGACCCTCCAGGGCGAGGAGCTCACCGACAA
CGGCCTGGCGCTGAGCCCGCAGACCCGTGTCGCCCTCACCCTCAAGCACTTCCCGGGCGGCGGCC
CGCAGGAGCTGGGGCTGGACCCGCACTACGCCTTCGGCAAGGCGCAGGTCTACCCGGCCGGGCGC
TTCGAGGAGCACTTCCTGCCCTTCCAGGCGGCCATCGACGCCGGCGTCTCCTCGATCATGCCGTACT
ACGGCGTCCCGGTGGACGTCCCGGTCGTCGGCGGGGAACCGGGGGAGACCTACCCGCACACCGGC
TTCGCCTTCTCCGACTCGATCGTCAACGGTCTGCTGCGTGACCAGCTGGGCTTCACCGGGTACGTCA
ACTCCGACACCGGGATCATCAACGACCGCGCGTGGGGCCTGGAGGGCAACACCGTGCCCGAGCGG
GTGGCAGCCGCGATCAACGGCGGCACCGACACCCTCTCCGGCTTCAGCGACGTCTCGGTCATCACC
GACCTGTACGAGGCCGACCTCATCTCCGAGGAGCGGATCGACCTGGCCGCCGAGCGTCTGCTCGAG
CCGCTGTTCGACATGGGGCTCTTCGAGAACCCCTACGTCGACCCGGACGTGGCCACGGCGACGGTG
GGCGCCGACGACCACCGCGCCGTCGGGCTCGACCTCCAGCGCAAGTCGCTCGTCCTGCTCCAGAAC
GAGGAGACCGACGAGGGCCCGGTGCTGCCCCTGAAGGAGGGCGGCGACGTCTACATCCTCGGTGA
CTTCACCGAGGAGACCGTGGAGAGCTACGGCTACGAGGTGACCAACGGCAACGTCGCGGAGGGGGA
GGAGCGGCCCAGCGCCGCGGGCAGCGACTACGTCCTCATCTCGATGACTGCCAAGACCAACGCCGG
GGACTACGTGAGCGACGACCCGAGCCTCGGCCTCAACCCGGACCACGGCACGAACCCGAGCGTCAT
CATCGGTGACGACGGCGAGCCGCTGCCCGGCCTGGACGGCCAGAGCCTGTGGGGCGCGGCCGACG
TGTGTGTCCACAAGGAGGGACACGAGGAGAACCCCTCGTGCACCGACAACCGGCTGCGCTTCGGCG
GCGCCTACCCGTGGGAGTCGAGCATCCTCGACTTCACCGGCATGGAGGCGGCCGAGTCGTGGGAGG
TCGTCCCCTCGCTGGAGACCATCCAGGAGGTCATGGCCGAGGTCGAGGACCCGAGCAAGGTCATCC
TCCACGTCTACTTCCGCCAGCCCTACGTGCTCGACGAGGAGAGCGGCCTGCGTGACGCCGGCGCGA
TCCTCGCCGGCTTCGGGATGACCGACACCGCCCTCATGGACGTCCTCACCGGTGCCTACGCGCCAC
AGGGCAAGCTGCCCTTCGCGCTCGCCGGCACCCGCGAGGCGATCATCGAGCAGGACTCCGACCGCC
CCGGCTACGACGAGACCGAGGACGGTGCACTCTACCCCTTCGGCTACGGCCTCACCTACGAGGACG
ACACCGAGGAGCCGACGACCCCGGAGCCCGAGCCGACCACGTCCCCGGACGACGAGACGCCGGGT
GACGACCGGACGCCGGGCGACGACCAGACGCCGGGCGACGACGAGACCCCGGCCGACGACGACG
AGACGCCGGGTGACACCACCGACGACGAGTCGCCGGCCGCCGGTGCCGACTCGGGCAACCTCGCC
CGGACCGGCGCACCGGTCACCGCGCTCGCACTGGTCGCGGTCGCCGCACTCGTGCTCGGCACGGC
ACTGCTGATCCAGCGCCGTTTCGCCCGCACCCAGGGCTAGCCGACACCGCACCGCCGAGGGCGGGT
GGGGACGCCGACGGCGTCCCTGCCCGCCCTCGGTCCGTCCGCGGGTCGCCGCGCGTGGTCGCGCC
GGCGGTCCCGGAGCGGTCAGGGGGACGCGCGGTCCTCGCCCGCCGGGGTGCCGGCGGTCCCGGA
GCGGCCAGGGGGACGCGCGGCCCTCGCCCGCCGGGGTGTGGGCCGTGCCGGTGGACCCGGAGCG
GTCAGGGGGACGCGCAGTTCTCGCCGGCCGGGGTGCCGGCCGTGCCGGCGGTCCCGGCTCAGGC
GGACTCGCGGTCCTCGCCCGGCGTGGGGTCGCC

#### Codon-optimized SN243 gene

ATGAGCCGTACACCGCGTGTTGGTCTGCGTACCGCAGTTGCAACCGCAGCAGCAGCACCGCTGGTTC
TGACCCTGGCAGTTGGTGCCGGTGCAGCAACCACACCGCCTGGTGATCTGGAACAGCCGGAACTGG
AAGCACGTGTTAAAGAAATTATTGAGGTGGATGGTTATCAGTTCCGCGATCTGAATGATAATGGTGAAC
TGGATCCGTATGAAGATTGGCGTCTGCCGACACCGGAACGTGTTGCAGATCTGGTTGGTCAGATGAG
CCTGGTTGAAAAAAGCGGTCTGATGCTGATTAATACCCTGAATGCAGCATGTGATCCGCAGACCGGTG
AATTTGGTGTTCTGCCTGCACAGGCCGATAATTACATTAATACACAGCATATGCACCGCTTTGTGTTTC
GTAATGTTGTTGATGTTCGTGCCGAAGGTGTTGAATGCACCGGTACAGGTACACCGGTTGTTAGTCCG
GCAGAAGCAGCAACCTTTACCAATGCAGTTCAAGAAATGAGCGAAGCAACCCGTCTGGGTATTCCGA
GCCTGTTTAAAAGCAATGCACGCAATCATATTGATCCGGATGCACGTGTTGGTATTAATGAAGCAGCG
GGTGCATTTAGCGCATTTCCGAAAGAAGCAGGTATTGCAGCAGCCGCACTGGGTGAACAGGCACGTC
GTACCGGTGAAGCCACCACCGGTGATATGAGCGTTGTTGCCGATTTTGCAGATGTTATGGGTGAAGAA
TGGGCAAGCATTGGCCTGCGTGGTATGTATGGTTATATGGCAGATCTGAGCACCGAACCGCGTTGGT
ATCGTACCCATGAAACCTTTACAGAAGATGCATATCTGGCAGCCGAAATTATGGAAACCCTGGTGCAG
ACACTGCAGGGCGAAGAACTGACCGATAATGGTCTGGCACTGAGTCCGCAGACACGTGTGGCACTGA
CCCTGAAACATTTTCCTGGTGGTGGTCCGCAAGAACTGGGTTTAGATCCGCATTATGCCTTTGGTAAA
GCACAGGTTTATCCGGCAGGTCGTTTTGAAGAACATTTTCTGCCGTTTCAGGCAGCAATTGATGCCGG
TGTTAGCAGTATTATGCCGTATTATGGTGTTCCGGTTGATGTGCCGGTTGTTGGTGGTGAACCGGGTG
AAACCTATCCGCATACCGGCTTTGCCTTTAGCGATAGCATTGTGAATGGTCTGCTGCGTGATCAGCTG
GGTTTTACCGGTTATGTTAATAGCGATACCGGCATTATTAACGATCGTGCATGGGGTTTAGAAGGTAAT
ACCGTGCCGGAACGCGTGGCAGCAGCAATTAATGGTGGTACAGATACCCTGAGCGGTTTTAGTGATG
TTAGCGTTATTACCGATCTGTATGAAGCCGATCTGATTAGCGAAGAACGTATTGACCTGGCAGCGGAA
CGTCTGCTGGAACCGCTGTTTGATATGGGTCTGTTTGAAAATCCGTATGTTGATCCTGATGTTGCCAC
CGCAACCGTTGGTGCGGATGATCACCGTGCAGTTGGTCTGGATCTGCAGCGTAAAAGCCTGGTTCTG
CTGCAGAATGAAGAAACCGATGAAGGTCCGGTTCTGCCGCTGAAAGAAGGTGGCGACGTTTATATTCT
GGGTGATTTTACCGAAGAAACGGTGGAAAGCTATGGTTATGAAGTGACCAATGGTAATGTTGCGGAAG
GTGAAGAACGTCCGAGCGCAGCAGGTAGCGATTATGTGCTGATTAGCATGACCGCAAAAACCAATGC
CGGTGATTATGTTAGTGATGATCCGAGTCTGGGTCTGAACCCGGATCATGGCACCAATCCGAGCGTG
ATTATTGGTGATGATGGCGAACCGCTGCCTGGCCTGGATGGTCAGAGCCTGTGGGGTGCAGCAGATG
TTTGTGTTCATAAAGAAGGTCACGAAGAAAATCCGAGCTGTACCGATAATCGCCTGCGTTTTGGTGGT
GCATATCCGTGGGAAAGCAGCATTCTGGATTTTACAGGTATGGAAGCAGCAGAAAGCTGGGAAGTTGT
GCCGAGCCTGGAAACCATTCAAGAAGTTATGGCAGAAGTTGAAGATCCGAGCAAAGTTATTCTGCATG
TGTATTTTCGTCAGCCGTATGTTCTGGATGAAGAAAGCGGACTGCGTGATGCGGGTGCAATTCTGGCA
GGTTTTGGTATGACCGATACCGCACTGATGGATGTTCTGACCGGTGCCTATGCTCCGCAGGGTAAACT
GCCGTTTGCACTGGCAGGCACCCGTGAAGCAATTATTGAACAGGATAGCGATCGTCCGGGTTATGAT
GAAACCGAAGATGGTGCCCTGTATCCGTTTGGTTATGGTCTGACCTATGAAGATGATACAGAAGAACC
GACCACTCCGGAACCGGAACCGACAACCAGTCCGGATGATGAGACACCGGGTGATGATCGTACCCCT
GGTGATGACCAGACGCCAGGTGATGACGAAACTCCGGCAGATGACGATGAAACCCCAGGTGATACCA
CCGATGATGAAAGCCCTGCAGCCGGTGCCGATAGCGGTAATCTGGCACGTACGGGTGCACCGGTGA
CCGCACTGGCACTGGTTGCAGTTGCAGCACTGGTTCTGGGCACCGCACTGCTGATTCAGCGTCGTTT
TGCACGTACCCAGGGCTAA

**Amino acid sequence of SN243**

Highlighted: truncated construct (residues 32-852) used for biochemical characterization

**Bold green**: truncated construct (residues 32-789) for crystallization

MSRTPRVGLRTAVATAAAAPLVLTLAVGAGA**ATTPPGDLEQPELEARVKEIIEVDGYQFRDLNDNGELDPY
EDWRLPTPERVADLVGQMSLVEKSGLMLINTLNAACDPQTGEFGVLPAQADNYINTQHMHRFVFRNVVD
VRAEGVECTGTGTPVVSPAEAATFTNAVQEMSEATRLGIPSLFKSNARNHIDPDARVGINEAAGAFSAFP
KEAGIAAAALGEQARRTGEATTGDMSVVADFADVMGEEWASIGLRGMYGYMADLSTEPRWYRTHETFT
EDAYLAAEIMETLVQTLQGEELTDNGLALSPQTRVALTLKHFPGGGPQELGLDPHYAFGKAQVYPAGRF
EEHFLPFQAAIDAGVSSIMPYYGVPVDVPVVGGEPGETYPHTGFAFSDSIVNGLLRDQLGFTGYVNSDTGI
INDRAWGLEGNTVPERVAAAINGGTDTLSGFSDVSVITDLYEADLISEERIDLAAERLLEPLFDMGLFENP
YVDPDVATATVGADDHRAVGLDLQRKSLVLLQNEETDEGPVLPLKEGGDVYILGDFTEETVESYGYEVT
NGNVAEGEERPSAAGSDYVLISMTAKTNAGDYVSDDPSLGLNPDHGTNPSVIIGDDGEPLPGLDGQSLW
GAADVCVHKEGHEENPSCTDNRLRFGGAYPWESSILDFTGMEAAESWEVVPSLETIQEVMAEVEDPSKV
ILHVYFRQPYVLDEESGLRDAGAILAGFGMTDTALMDVLTGAYAPQGKLPFALAGTREAIIEQDSDRPGY
DETEDGALYPFGYGLTYEDDTEE**PTTPEPEPTTSPDDETPGDDRTPGDDQTPGDDETPADDDETPGDTT
DDESPAAGADSGNLARTGAPVTALALVAVAALVLGTALLIQRRFARTQG

#### 2. Nucleotide and amino acid sequences for hit SN268

##### Genomic insert of library member SN268

Highlighted: ORF SN268

ATGACGTGGTCCCTTTACCTCGCCAAATAGAAATGAAGAAAGGACCTGGTTTCCAGATGAATTCCACC
ATTTACATTGTGGTGAAGGATGCTTCCTTGGCGAGAGATGCAAAATTCCTCCGTGACTATCTTTTGGGA
GAGAATCAGATGAGTGCTGTTATTCAGGAGAAGGGCATCAGCGGCCATACCATCGTGCTGCAACTGA
ATCCCAAGATGACGGAAAAGGAGGGCTACCGCATCAGCATAAGTTCGAAACTGGTGACGATAGAGGG
ACTAACGCCACAGGGTGTATTTTATGGCATTCAGACTCTGCGGAAATCATTGGCAGGTTTTAAGAAGTA
TGCCATTCTTCCTGCGGCCGTGATTGAGGATGCCCCACGCTTTGCTTACCGAGGCATGCATCTCGAC
GTGGCCCGCCATTTCTTTGGTGTGGAGTTTGTGAAGAAGTATCTCGACGCGATGGCGATGCACAATAT
GAACCGCTTCCACTTTCATCTGACCGACGATCAGGGCTGGCGCGTGGGTATTGACCGTTATCCGAAA
CTGATAGAGGAGGGCTCGAAGCGCAACTACACGGTCTTGGGTCGCAATTCCCCGATTGATGACGGTA
CGCCCTACGGCGGTAGTTTTACCAAGGAGGAGCTGAAGGAAATCGTGCGATACGCGCAGGAACGCTT
TATCACGGTGATTCCGGAGATTGATATGCCGGGCCACATGCTGGGGGCACTGAAGGCCTATCCTGAA
TTGGGATGCACGGGCGGACCTTACGAAGTGGAGGGCCACTGGGGTGTCTTCGAGGATATTCTATGTG
CTGGAAAGGAAGAGACCTTTAAATTCATAGAAGGCGTGTTGACGGAACTGATGGAGATATTCCCATCG
GAATATATCCATATTGGCGGTGATGAGGCTCCGAAAGTGAGGTGGGAGCAATGTCCACGTTGCCAGC
AGCGTATTAAGGATTTAGGGCTAAAAGCCGATGAAGGGCATACGGCGGAAGCCAAACTGCAGGGCTA
TTTCACCAAACGTATAGAGGCATTCTTGAATGCTCATGGACGAAAATTGATCGGTTGGGATGAACTGAT
GGAAAGTGATGTCAATGAGAGTGCTACGATCATGAGCTGGCGTGGTGTGAAAGGTGGTTTGGCAGCT
TCACAAAAAGGGCATGACGTCATCATGTCACCAACAAGCCATTGCTATTTCGATTATTACCAGACGAAA
GATCATGAGTGGGGAGAACCATTCTCCTTTGCCAATATCATTACTGTGGAGACTACTTACGGTCTTGAT
CCTGCTCCTAAGACGTTATCTCCTGAAGCGCAAGCTCATATTCTCGGTGCTCAGGCAAACCTTTGGAC
GGAATATATTGCCCATGAAAGTCAGGCATTCTATCAGGTGCTGCCCCGTATGGGAGCTTTGTGTGAAG
TGCAGTGGGTGATGCCTGAGAAAAAAGATTTCAAGCAGTTCGTGGAAAGGGAGCGCCGCCTTACCTT
ATTATATAAACACTATGGCTGGCCATTTGCCCCACACATTTTTAAAGTGAAGAGTGAAGAATGAAGAGT
GTAGAATTATGATTACTCTTTATGTTTGAAAACTGAAAAACTGTCAGAGTTTACGGGCTTTGGCATAGG
CATTTTGAGCCATGCGGTTGATGATGTCGAGGGCAAGTTCCTCGTCCATAAGGCCAGCTTCTGCACAA
GGTGTGTAGCGGTTGGCCTCTTCCGTTGTGGCCAGTCTTCCCTCCAGTAAAAACATGAAATGTCGCTT
ATATTCTTCCACCATTGCATTGTAGAAAGGTTCCGTGCGGCAGGCAATGCGCACTAGTTGCATCAAGT
CGTTAGTGGGATTCTTCCGAAGGGTGTTTTCTATTTCCTCGGCTTCATAATTCATACGTGCAATAGCAT
AGAAAGCCTCCTGATTGTAGATCTTCGAAATATGTTCATAGTATTCCAGCAGCCTCTGCTTTACGGCTT
TGGTATCGTTGAGGTCCTTACCTTCCATCATACCTGTCAGGAAGTATTTGAAAAAGGTGCGCACGAGG
GCGGCAAGTACACGGTCTTTACTTATCTTGTTCATGTTTGATGGCTTTAATATTTCTCATAAGTCCTTCG
TATTTGGCCCGTTTTACAGCAGAACCTTTGAATAACTGCTGATATTGTTCCACAGACAACTGTTGCCAG
TCTTCTTCTTTCATAGAAAGTAGTTCGTCGGTAGGTTGAAGCAAAGGTTCATCAGTAGGTTGTGCCTTA
GCATTCCAAGGACAAACCTGTTGGCAACGGTCACATCCATAGAAGGTTTCACCCATCTTCTGGGCTGC
TTCCTCCGGAATGTCACCCCGATTCTCGATGGTGAGGTAGGAAAGGCATAAGGAGGAAGAAAACGAG
GTCCCCCTTTCGGCCCCCGAGGGGGCTAGGCTTAATGCACCTGTAGGACATGCATCTAAACATCGGT
GGCAACTACCGCAACGCTGTTGGCAAGGGGCATCGGACTCCAGTTCAATATCTACAAAAAGTTCTCCT
AAGAAGAACTGACTGCCCATGCCGGGGATAATGAGCTGATGGTTCTTGCCGATAAAACCGAGCCCTG
CCTTCATGGCCCAATAGCGTTCCAGAACGGGGGCACTGTCGGTGAAGGCACGGAAGGAAGTCCCCT
CAGGGGAAGATGAGAGGGAGACTATGAAGGCAGCCAATTTATGGAGTTTCTGCTTCACGATGTCGTG
ATAGTCCTTGCCAATGGCATAGTTGGCAAATCCCTTTACGGGCTGAGAGGGGGCATAATTCAAGGCTA
CGCTGATGATACTTTTCACACCAGGCATCAGTATACGGGGATCAAGACGCTTATCCTCATAATTCTCCA
TATACTGCATGGTGGCATGGCCTCCCCGCTGAATCCAGCTTTGGAAAGCCTTGGCTGCCTCGGTATC
CACGGGTTCCGCTTTGGCAATACCGCAGGCAGAAAAGCCGAGGCGTAGTGCCTCGGCTTTTATTTCA
CGGGATAACTTTGAATTCATAGCCTTGTTCGATAAGCCAGTCTAAGGATTGCGGCAGAGCCGTTTGCA
GTTTGTCGATGCTTTTCAGCGAGTCGTGGAAGGTAATAATAGAGCCGTTGCGCGTATATGTCTTCACG
TTTTCCACTACATCCTCCGCGTTAAGCCGGTTAGAGTAGTCGCGGGTAACCACATCCCACATCACTAC
GCGATAATATTTCTTTATCCAGTAGTATACGCTATGGCGCATCACACCGTGAGGTGGACGGAAGAGGT
TGCTATGTAGGTACTCATTGGCCTGATAGGTATTGATGACATACGTGGTGGTCCAATGTTTGAACGAC
CCCAGATGGTTCATCGTATGGTTACCTATCCGATGTCCGGCGGCCTTCACCCTTTCGAAAAGTTCTGG
GTATTTACGTACATTGTCGCCCACCATGAAGAAGGTGGCCTTAATATCGTAACGTTCCAGCGTCTCAA
GGATGAAAGGCGTAGCCTCGGGAATAGGGCCGTCGTCGAACGTCAGATAGACCGAATGTTCGTTCGG
GTCCATTCTCCAAGTTGCGTGCGGATAAAACAACCGGAGCCATCGTGCTGGCTGTTC

#### Codon-optimized SN268 gene

ATGAAAAAAGGTCCGGGTTTTCAGATGAATAGCACCATTTATATCGTGGTGAAAGATGCAAGCCTGGC
ACGTGATGCAAAATTTCTGCGTGATTATCTGCTGGGTGAAAATCAGATGAGCGCAGTGATTCAAGAAA
AAGGTATTAGCGGTCATACCATTGTTCTGCAGCTGAATCCGAAAATGACCGAAAAAGAAGGTTATCGC
ATTAGCATTAGCAGCAAACTGGTTACCATTGAAGGTCTGACACCGCAGGGTGTTTTTTATGGTATTCAG
ACCCTGCGTAAAAGTCTGGCAGGTTTTAAGAAATATGCAATTCTGCCTGCAGCCGTGATTGAAGATGC
ACCGCGTTTTGCCTATCGTGGTATGCATCTGGATGTTGCACGTCATTTTTTTGGTGTGGAATTTGTGAA
AAAATACCTGGATGCAATGGCCATGCACAATATGAATCGCTTTCATTTTCATCTGACCGATGATCAAGG
TTGGCGTGTTGGTATTGATCGTTATCCGAAACTGATCGAAGAAGGTAGCAAACGTAATTATACCGTTCT
GGGTCGTAATAGCCCGATTGATGATGGCACCCCGTATGGTGGTAGCTTTACCAAAGAAGAACTGAAAG
AAATTGTGCGCTATGCCCAAGAACGTTTTATTACCGTTATTCCGGAAATTGATATGCCTGGTCATATGC
TGGGTGCACTGAAAGCATATCCGGAACTGGGTTGTACCGGTGGTCCGTATGAAGTTGAAGGTCATTG
GGGTGTGTTTGAAGATATTCTGTGTGCAGGTAAAGAGGAAACGTTCAAATTTATCGAAGGTGTTCTGA
CCGAACTGATGGAAATTTTTCCGAGCGAATATATCCATATTGGCGGTGATGAAGCACCGAAAGTTCGT
TGGGAACAGTGTCCGCGTTGTCAGCAGCGTATTAAAGATCTGGGTCTGAAAGCAGATGAAGGCCATA
CCGCAGAAGCAAAACTGCAGGGTTATTTTACCAAACGCATTGAAGCATTTCTGAATGCCCATGGTCGT
AAACTGATTGGTTGGGATGAGCTGATGGAATCAGATGTTAATGAAAGTGCAACCATTATGAGCTGGCG
TGGTGTTAAAGGTGGTCTGGCAGCAAGCCAGAAAGGTCATGATGTTATTATGAGCCCGACCAGCCATT
GCTATTTCGATTATTATCAGACCAAAGATCACGAATGGGGTGAACCGTTTAGCTTTGCAAATATCATTA
CCGTGGAAACCACCTATGGTCTGGATCCGGCACCGAAAACACTGAGTCCGGAAGCACAGGCACACAT
TCTGGGTGCCCAGGCAAATCTGTGGACCGAATATATTGCACATGAAAGCCAGGCATTTTATCAGGTTC
TGCCTCGTATGGGTGCCCTGTGTGAAGTTCAGTGGGTTATGCCGGAAAAAAAAGACTTTAAACAGTTT
GTGGAACGTGAACGTCGTCTGACCCTGCTGTATAAACATTATGGTTGGCCGTTTGCACCGCACATCTT
TAAAGTTAAAAGCGAAGAGTAA

#### Amino acid sequence of SN268

MKKGPGFQMNSTIYIVVKDASLARDAKFLRDYLLGENQMSAVIQEKGISGHTIVLQLNPKMTEKEGYRISISS
KLVTIEGLTPQGVFYGIQTLRKSLAGFKKYAILPAAVIEDAPRFAYRGMHLDVARHFFGVEFVKKYLDAMAM
HNMNRFHFHLTDDQGWRVGIDRYPKLIEEGSKRNYTVLGRNSPIDDGTPYGGSFTKEELKEIVRYAQERFI
TVIPEIDMPGHMLGALKAYPELGCTGGPYEVEGHWGVFEDILCAGKEETFKFIEGVLTELMEIFPSEYIHIGG
DEAPKVRWEQCPRCQQRIKDLGLKADEGHTAEAKLQGYFTKRIEAFLNAHGRKLIGWDELMESDVNESATI
MSWRGVKGGLAASQKGHDVIMSPTSHCYFDYYQTKDHEWGEPFSFANIITVETTYGLDPAPKTLSPEAQA
HILGAQANLWTEYIAHESQAFYQVLPRMGALCEVQWVMPEKKDFKQFVERERRLTLLYKHYGWPFAPHIF
KVKSEE

#### 3. Evidence for activity of hit SN268

The lysate of the original library member SN268 was tested for activity towards the bait substrate FD-β-GlcA. At least 20-fold activity enhancement (fluorescence intensity normalized to the $OD_{600}$ of the culture) in comparison to negative controls (random clones of the SCV library not encoding any β-glucuronidase) was detected (Figure S1a).

The ORF of hit SN268 (sequence see Section 2, SI) encoding a glycoside hydrolase of the GH20 family was cloned into the expression vectors pBat4[1] and pSol (see Methods) to recombinantly express the new β-glucuronidase without tag, with a C-terminal $His_6$-tag and with a N-terminal $His_6$-Twin-Strep(II)-SUMO-tag, respectively (cloning primers, Table S1). Expressed protein was mainly found in the insoluble fraction and various purification attempts did not lead to sufficient amounts of pure enzyme for characterization purposes. Nevertheless, the lysate of SN268 expressed as a fusion protein with a C-terminal $His_6$-tag showed clear activity towards *p*NP-β-GlcA and the trace amounts of protein that could be purified via a Ni-NTA column retained this activity (Figure S1b).

**Table S1: Cloning primers for SN268 expression constructs.** Bases in capital letters anneal to the template.

| Purpose | Name | Sequence |
|---|---|---|
| Vector amplification | pSol_F | TAATAGAGCGGCCGCCAC |
| | pSol_R | GGATCCACCAATCTGTTC |
| | pBat4_F | CTTGAGTATTCTATAGTGTCACCTAAATC |
| | pBat4_His_F | aagcttgcggccgcactccaccaccaccaccaccactaaCTTGAGTATTCTATAGTGTCACCTAAATC |
| | pBat4_R | GGATATATCTCCTTCTTAAAGTTAAAC |
| Insert amplification | SN268_ pSol_F | gagaacagattggtggatccATGAAAAAAGGTCCGGGTTTTC |
| | SN268_ pSol_R | cggtggcggccgctctattaTTACTCTTCGCTTTTAACTTTAAAGATG |
| | SN268_ pBat4_F | tttaagaaggagatatatccATGAAAAAAGGTCCGGGTTTTC |
| | SN268_ pBat4_R | gacactatagaatactcaagTTACTCTTCGCTTTTAACTTTAAAGATG |
| | SN268-His_ pBat4_R | gacactatagaatactcaagttagtggtggtggtggtggtggagtgcggccgcaagcttCTCTTCGCTTTTAACTTTAAAGATG |

While SN268 was established as a *bona fide* β-glucuronidase, the difficult purification of SN268 made us focus our attention on hit SN243 instead.

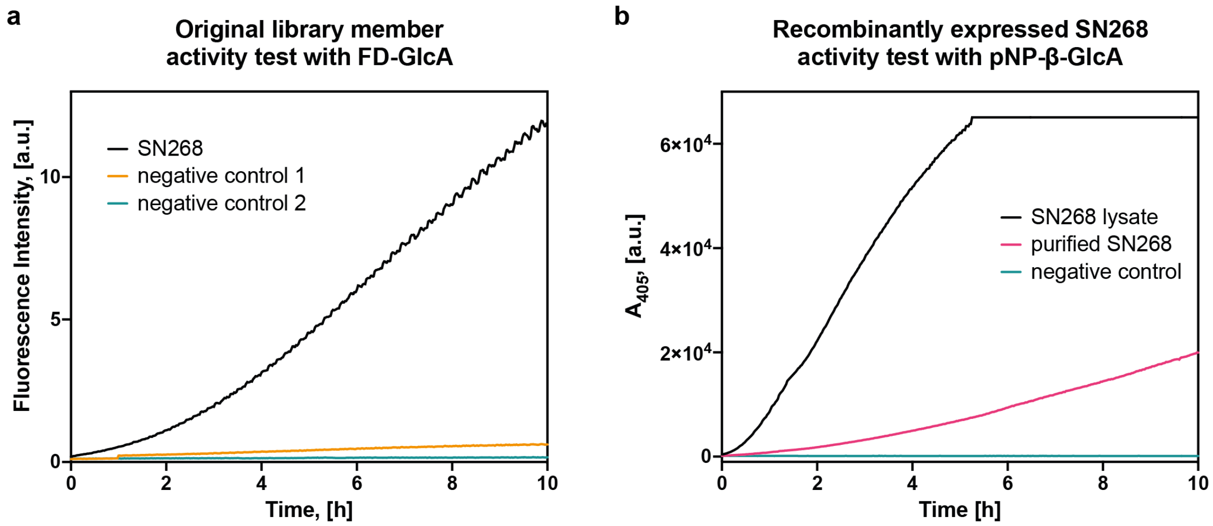


**Figure S1: β-glucuronidase activity of SN268. (a) Lysate activity test of the original library member.** Hit SN268 showed clear activity towards the bait substrate FD-β-GlcA. Negative controls: randomly selected SCV library members not encoding any β-glucuronidase gene. **(b) Activity test of recombinantly expressed protein**. Activity towards *p*NP-β-GlcA was detected for lysate of recombinant expression cultures and purified protein solutions of SN268. Negative control: elution buffer of the Ni-NTA column without enzyme.

When compared to other GH20 members, SN268 displays the typical conserved HxGGDE motif, as shown below, classifying the enzyme as a member of this family. Moreover, a phylogenetic tree with the characterized, bacterial GH20 members highlights that the closest homologs are all N-acetyl-β-D-glucosaminidases (Figure S2). In a second analysis (in Figure S3), we compared the structure of SN268 predicted by Alphafold2 to the structure of the β-N-acetylhexosaminidase from *Akkermansia muciniphila* co-crystallized with GlcNAc (PDB: 7CBO). While the overall sequence homology between the two structures is 40.5%, their structures align almost perfectly (RMSD: 0.768 Å, over 2239 atoms), and the active sites share an identical sequence (Figure S3). Therefore, it seems likely that SN268 is an N-acetyl-β-D-glucosaminidase that was discovered in our screens because of a *promiscuous* activity towards β-glucuronidase substrates.

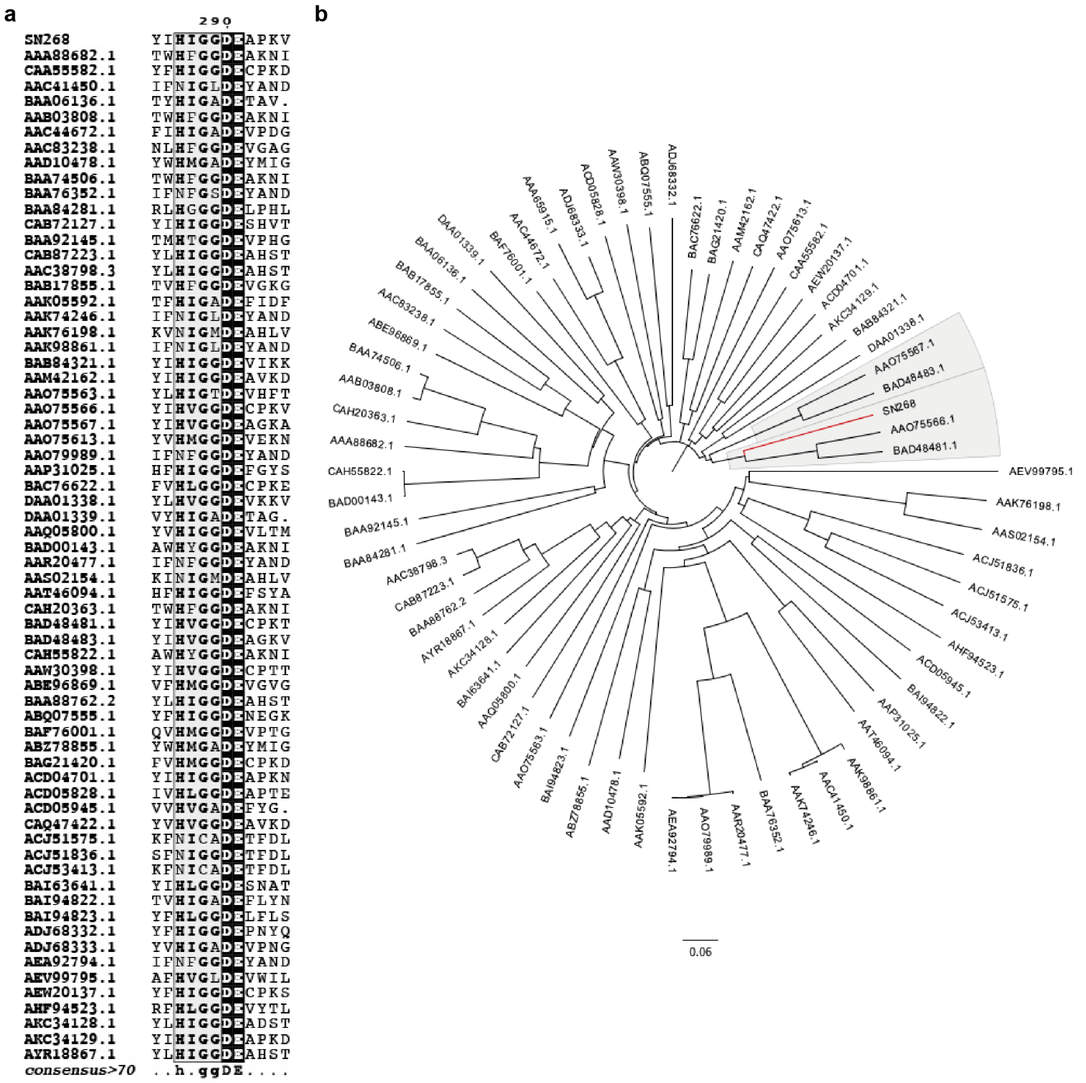


**Figure S2: Homology analysis of SN268 and characterized bacterial GH20 members**. **(a)** The multiple sequence alignment shows the conservation of the HxGGDE motif, with Asp and Glu (in positions 290 and 291 in SN268, respectively) acting as catalytic residues. **(b)** The closest homologs identified in the phylogenetic tree are all N-acetyl-β-D-glucosaminidases. GenBank accession codes are indicated.

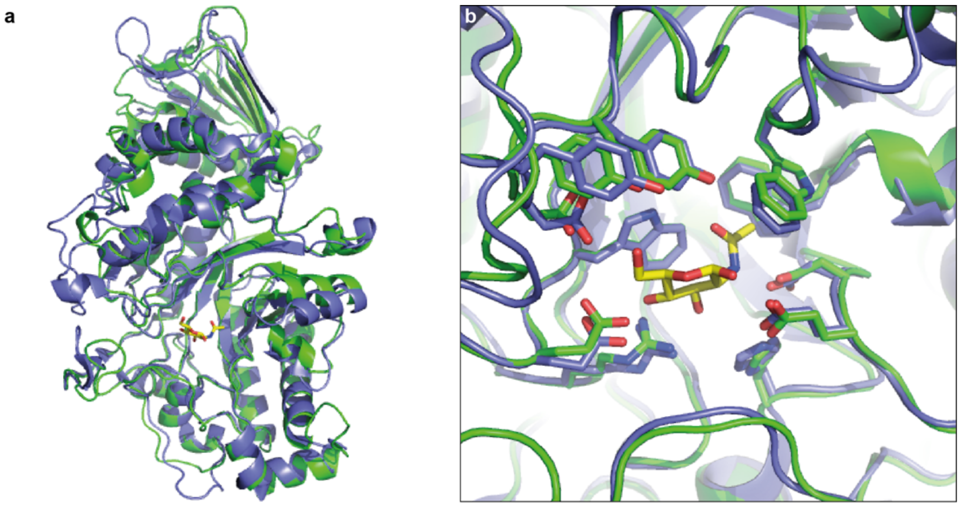


**Figure S3: Structural comparison of SN268 (based on an Alphafold2 prediction) with its closest homolog in the PDB**. **(a)** Overall alignment of SN268 (green) and the β-N-acetylhexosaminidase from *Akkermansia muciniphila* co-crystallized with GlcNAc (PDB: 7CBO, purple). **(b)** The focus on the active site shows an almost perfect alignment of the binding sites. A reasonable conjecture based on the perfect active site similarity is that they catalyse turnover of the same main substrate, but that weaker promiscuous activities may exist, and we have detected one of them in our droplet screens.

#### 4. Bioinformatic exploration of the sequence neighbourhood of SN243

The sequence neighbourhood of SN243 was explored by searching the MGNify database.[2] A search for SN243 results in 11,862 sequences with significant homology (as defined by MGnify) to the query sequence. The closest search hit (MGYP000481601007) aligned with an E-value of 1.1e-229 to SN243 (corresponding to a 55% identity and 87% query coverage). Homology analysis of MGYP000481601007 with characterized sequences (UniProt/SwissProt)[3] returns β-glucosidase/β-xylosidase hits with much lower homology to the query (E-value of the closest hit: 3e-28; Q46684.1). In total, our MGNify search identified 512 sequences with an E-value lower than e-40.
To visualise the sequence space around SN243, we constructed a sequence-similarity network (SSN)[4] and displayed it with Cytoscape.[5] The 11,862 significant hits from the MGNify search, were clustered together (based on more than 95% identity) using CD-HIT[6] giving 9,083 sequences as input data for the SSN. The SSN was constructed with a threshold of e-40. After removing unconnected nodes, 8,354 nodes were left and are shown in Figure S4. SN243 (red) is directly connected to 427 nodes (yellow) with an E-value < e-40. The furthest relative of SN243 in this group has an e-value of 9.4e-41 (MGnify), with a sequence homology of 36% and a query coverage of 36%.
The discovery and functional characterization of SN243 (and closely related sequences in its perimeter) add a bridgehead in sequence space that can help with the annotation of related sequences. Even though sequence similarity does not strictly predict functional similarity, as single residue changes can alter function, the characterization of SN243 illuminates the functional potential in this sequence neighbourhood (i.e. similar sequences are likely to have β-glucuronidase activity as their main or promiscuous function).

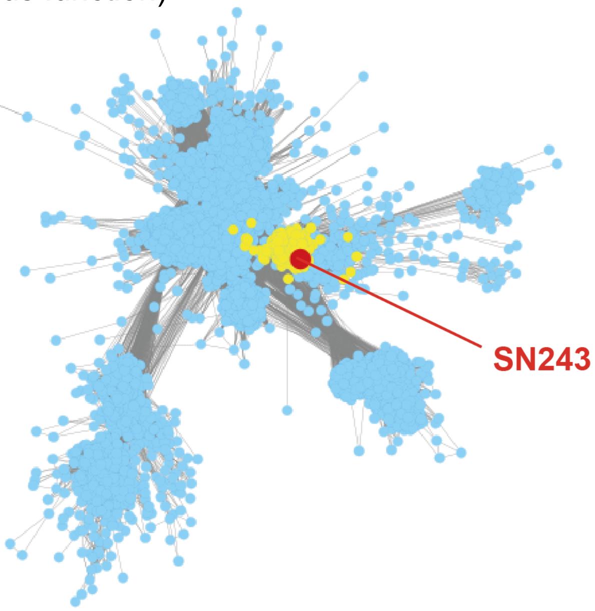


**Figure S4: Sequence-similarity network of sequences related to SN243 (red), starting with a list of significant hits from a MGNify search.** Shown in yellow are 427 nodes directly connected to SN243 with an e-value < e-40.

### 5. Supplementary Notes

#### 5.1 On the origin of false positive hits after FADS

The sequences of false positive hits of the screening campaign were analysed and all of them contained short metagenomic inserts or vector deletions (without inserts). The activity of these clones (in lysate activity assay in microtiter plates) was higher than that of randomly picked library members (with normal insert length), validating their selection in the droplet sorter and the positive confirmation by an assay in microplate format. Only when sequenced, were they recognised as false positive hits.

The selection of these clones with apparent increased activity can be explained by a frequently made observation: the β-glucuronidase background in *E. coli* is much reduced in clones that carry a plasmid library member with an insert that promotes the expression of a protein encoded in the insert. By contrast, clones without a functional metagenomic insert channel their energy for protein expression into all other cellular proteins, leading to higher background activity of the host lysate. This background activity of β-glucuronide hydrolysis is brought about by (i) GusA from *E. coli*, (ii) chemical (non-enzymatic) background, and (iii) any other (unknown) enzymes in *E. coli* that have a promiscuous β-glucuronidase activity.

In addition, smaller vectors are more transformed and are more efficiently amplified. Hence, amongst a population of initial hits, smaller constructs from the library will be favoured in DNA recovery and transformation steps. There are thus good reasons to expect short metagenomic inserts or partly deleted vector.

It should be noted that the frequency of constructs with small or no inserts in the source libraries was small (i.e. was not apparent in quality control by Sanger sequencing): the enormous screening throughput has the capacity to enrich marginal events, especially when the selection threshold was deliberately set close to background activity, in order not to miss rare events. Despite this impediment, we managed to extract information on intact proteins, and discovered genuine hits.

#### 5.2 Activity tests of SN243 with natural carbohydrate substrates

The question of the natural substrate(s) of SN243 was addressed by activity tests on plant cell wall polysaccharides containing β-glucuronides. In the collection of polysaccharides for testing, we focussed on arabinogalactan proteins (AGPs),[7-10] because they carry terminal β-glucuronides or β-glucuronides decorated with only one further sugar moiety. We tested the activity of SN243 (1 μM) on AGPs (1 mg) from *Gum arabic*, *Gum odina*, *Arabidopsis thaliana* and *Larch wood* (included as a negative control, as it does *not* contain a β-glucuronide). These substrates were pre-treated with an arabinofuranosidase[11] to remove terminal Ara*f* decorations and an exo-β-1,3-galactanase[12] to hydrolyse the backbone and release side chains. Samples were prepared, analysed and cleaned up according to Tryfona et al. (2012).[10] Activity tests were carried out by polysaccharide analysis using carbohydrate gel electrophoresis (PACE), which has a limit of detection of 100 fmol for the corresponding monosaccharides,[13] and MALDI-TOF. Despite using very sensitive analysis methods, we could not identify any activity towards the tested substrates, ruling them out as the main and also promiscuous substrates.

It is likely that SN243 is specific for a natural substrate in its original environment that was not included in this survey. The number of substrates and enzymes (and combinations of enzymes) required for further exploration of the role of SN243 in the cleavage of different natural polysaccharides exceeds a simple matrix: enzymes for all individual sugar bonds that need to be cleaved to let an exo-acting β-glucuronidase access its target linkage would need to be included and there is no high-throughput system available that would allow a straightforward definition of the natural substrate.

#### 6. Supporting Figures

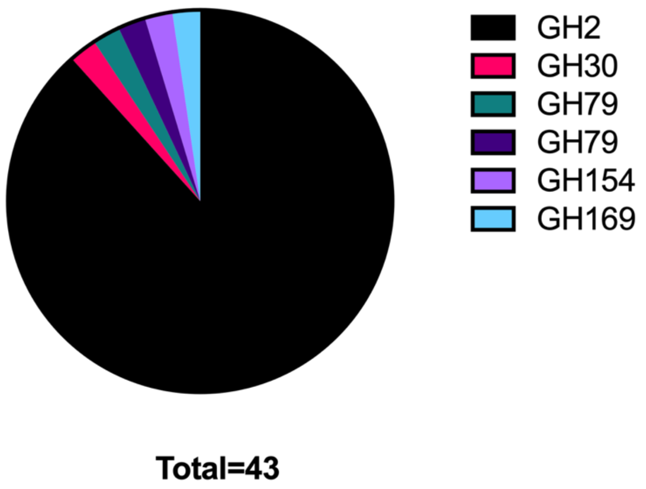


**Figure S5: Pie-chart representing characterized bacterial β-glucuronidases included in the CAZy database (release date 22/01/21) sorted by families.**

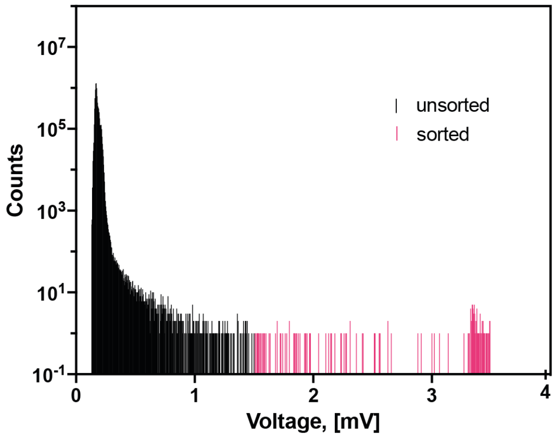


**Figure S6: Enrichment experiment for the FADS assay for β-glucuronidase activity in droplets**. The displayed histogram was recorded for the sorting of *E. coli* clones overexpressing *E. coli* GusA from a $10^{-4}$ dilution with *E. coli* expressing a negative control. Quantification of the selection output plasmid DNA by qPCR gave a 246-fold enrichment.

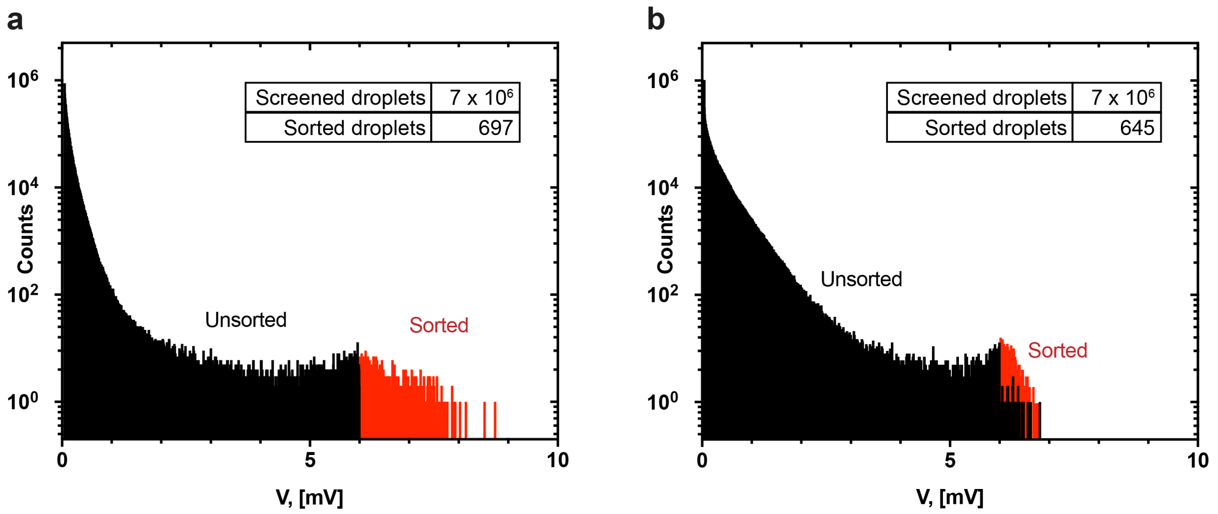


**Figure S7: Sorting histograms of the functional metagenomic screening for β-glucuronidases in microfluidic droplets.** After one **(a)** and two nights **(b)** incubation respectively, the droplets containing lysed cells and FD-β-GlcA were reinjected into a sorting chip and screened for high fluorescence (indirectly measured as Voltage). Droplets with a signal above 6 V and with the chosen size range were sorted.

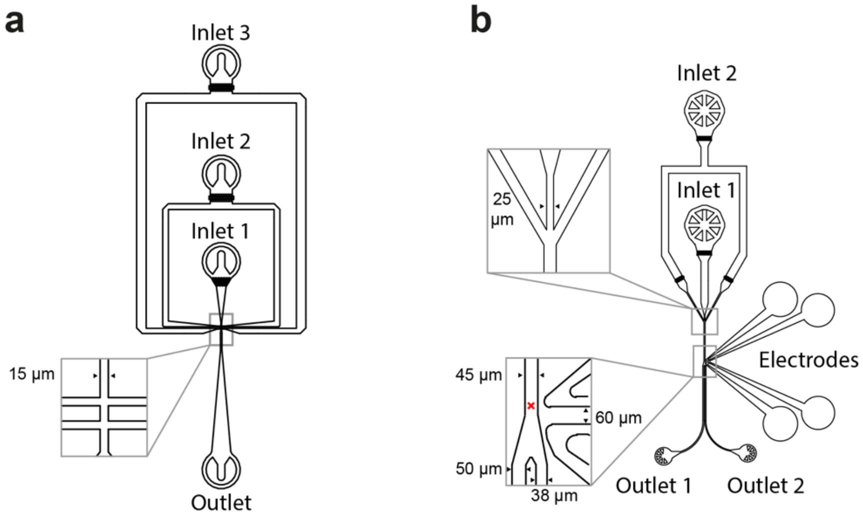


**Figure S8: Design of the microfluidic chip devices used for the functional metagenomic screening in droplets. (a) Flow focusing chip.** Bacterial suspension (inlet 1) and lysis substrate solution (inlet 2) are emulsified together in fluorous oil containing surfactant (inlet 3 resulting in water-in-oil droplets of 4 pL volume. The height of the channels is 16 µm. **(b) FADS chip.** After incubation droplets are reinjected (inlet 1) into a sorting chip and spaced out by additional fluorous oil (inlet 2). The fluorescence of droplets is monitored. If a fluorescence of a droplet is higher than the user defined threshold, an electric pulse is applied and the droplets is sorted into a collection tube via outlet 2, while the remainder goes via outlet 1 into the waste. The small red cross indicates the position of the laser. This chip design has been used previously[14] and the CAD file can be found at our open access repository DropBase (https://openwetware.org/wiki/DropBase:Contact).

a

b

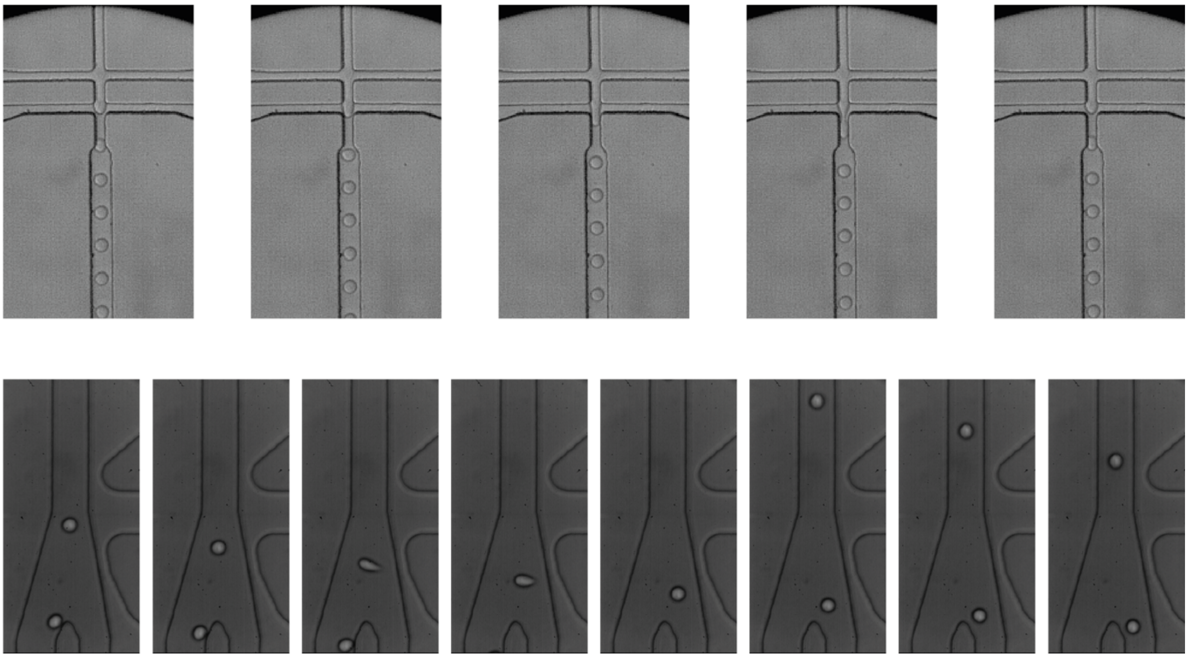

**Figure S9: Series of stills for microfluidic procedures. (a) Flow focusing.** The SCV library (in *E. coli*, vertical channel from top) was encapsulated together with substrate and lysis agent (top horizontal channels) into monodisperse 15 µm droplets (oil: bottom horizontal channels). **(b) FADS**. Droplets with a higher fluorescence intensity than the selected threshold were sorted. This example shows the video footage of sorting event 417 on screening day 1.

a

b

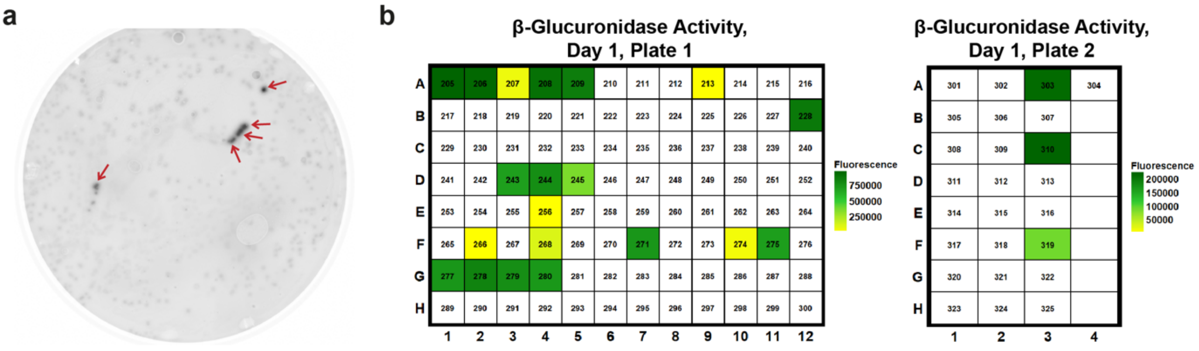


c

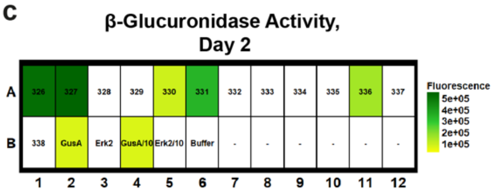


d

$$avg(fluorescence_{m,end}) = \frac{\sum_{t=n-9}^{n} fluorescence_{m,t}}{10}$$

$$threshold_{negatives} = \frac{\sum_{m=1}^{4} avg(fluorescence_{m,end})}{4} + 3 \times SD(avg(fluorescence_{m,end}))$$

**Figure S10: Colony and plate screening of the enriched SCV library for β-glucuronidase activity. (a) Example of a nitrocellulose membrane with partially lysed colonies and covered with the FD-β-GlcA agarose buffer solution.** Fluorescence imaging was done in a Typhoon plate scanner. Red arrows point to positive spots. Both enriched SCV libraries resulting from FADS after one- and two-nights incubation, respectively, were separately screened on ten plates. Only one of twenty plates is shown here as an example. **(b)** and **(c) Lysate activity screening in microtiter plates.** Small cultures in deep-well plates were inoculated from single colonies in the area of fluorescent spots in the colony screening assay (of both screened sub-libraries resulting from the two FADS days), grown overnight and their lysates were tested for β-glucuronidase activity. Lysates of cultures over-expressing *E. coli* GusA (and its 10-fold dilution; GusA/10) and Erk2 (and its 10-fold dilution; Erk2/10) served as positive and negative controls, respectively. The reaction was followed via fluorescence in a plate reader over 10 h. The normalised average fluorescence end value *avg(fluorescence$_{end}$)* is shown in a heat plot. The detected β-glucuronidase activate in the lysates of 28 library members was above the threshold calculated from the negative controls. **(d) Formula used to calculate the fluorescence threshold for the selection of clones for subsequent sequencing.** SD: standard deviation.

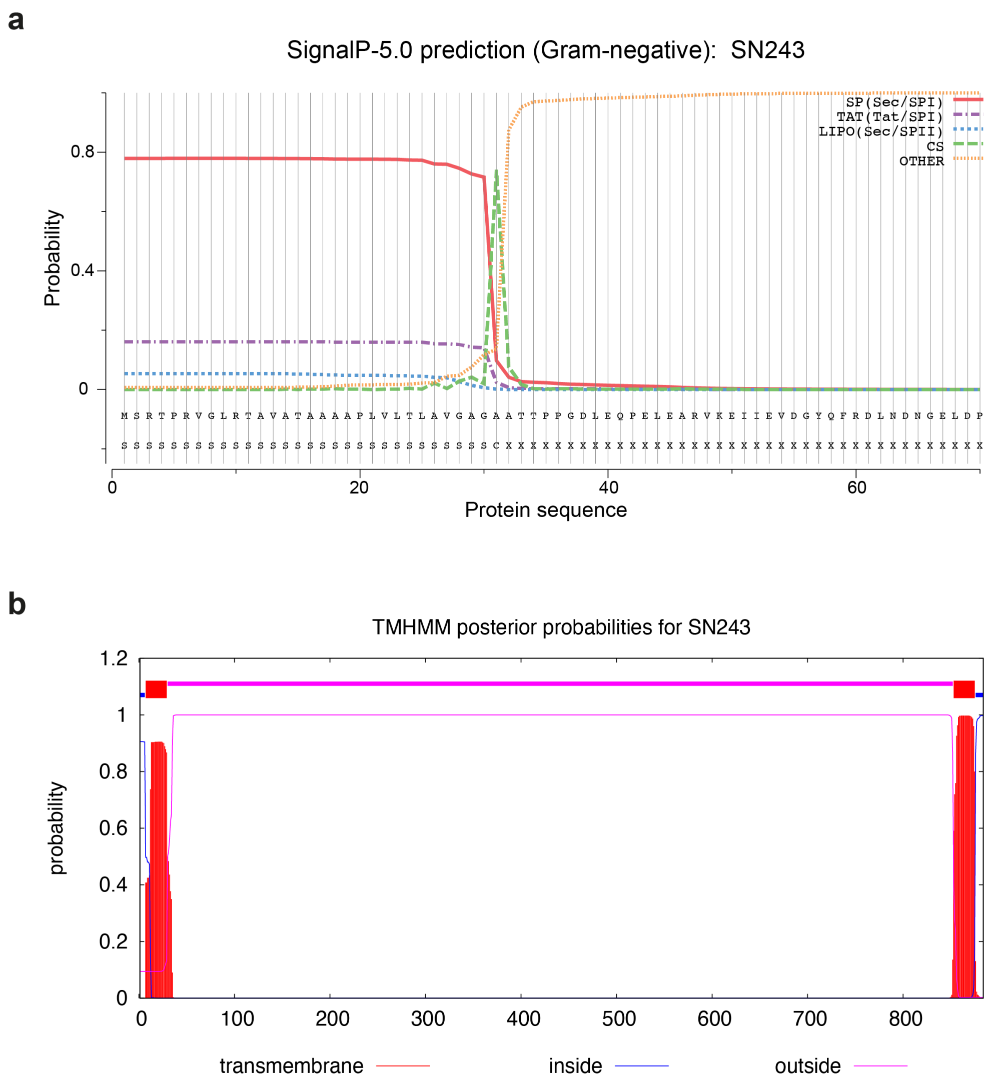


**Figure S11: Protein sequence analysis for the metagenomic hit SN243. (a) Signal peptide prediction.** Using the SignalP-5.0 server (http://www.cbs.dtu.dk/services/SignalP-5.0/)[15] with standard settings for gram negative bacteria, an N-terminal signal peptide with a cleavage site between Ala31 and Ala32 was predicted with high probability. **(b) Transmembrane domain prediction.** N- and C-terminal transmembrane domains ranging from residues 7 to 29 and 853 to 875 were predicted by TMHMM (http://www.cbs.dtu.dk/services/TMHMM/)[16]. Based on these results an N- and C-terminally truncated construct ranging from residues 32 to 852 was designed and used for the characterisation of the protein.

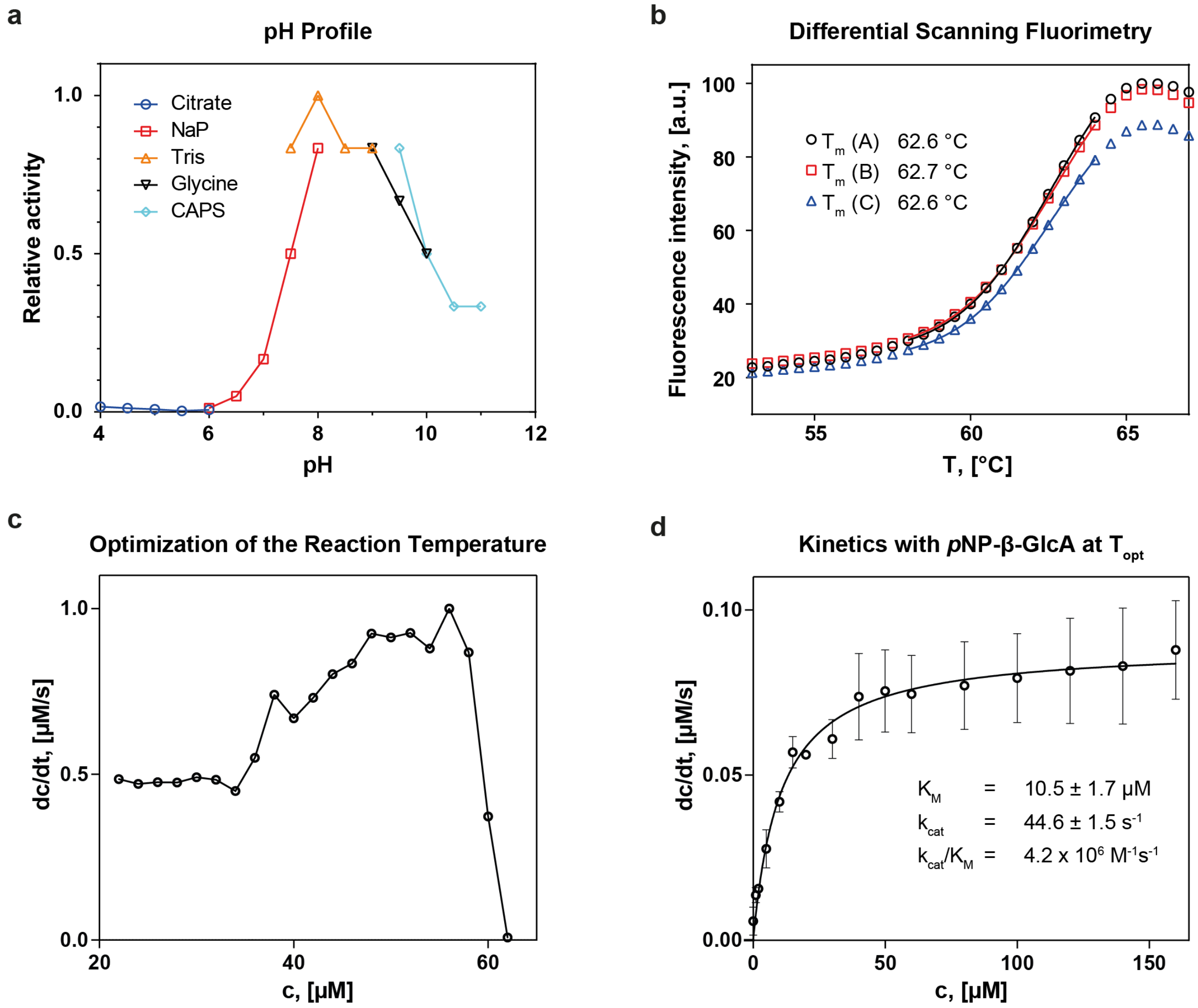


**Figure S12: Biochemical characterisation of the metagenomic β-glucuronidase SN243. (a) pH profile.** SN243 was most active on *p*NP-β-GlcA between pH 7.5 and 9.5. The buffer pH was varied between pH 4 and 11 using 100 mM citrate (pH 4-6), sodium phosphate (NaP, pH 6-8), Tris (pH 7.5-9), glycine (pH 9-10) and CAPS (pH 9.5-11) buffer, respectively, while keeping the salt concentration at 150 mM NaCl. **(b) Thermostability of SN243.** In a differential scanning fluorimetry experiment, the melting temperature of SN243 was determined at 62.6 °C. The assay was performed in triplicate. **(c) Optimization of the reaction temperature.** The temperature was increased in 2 °C increments and the initial reaction velocity with 10 nM SN243 and 50 µM *p*NP-β-GlcA recorded and normalised to the highest observed value (56 °C). (**d) Determination of Michaelis-Menten kinetics at the optimal reaction temperature.** Using 2 nM SN243, kinetic parameters for the reaction with *p*NP-β-GlcA were determined at 56 °C. The mean of three independent datasets and the standard errors with a 95% confidence level are shown.

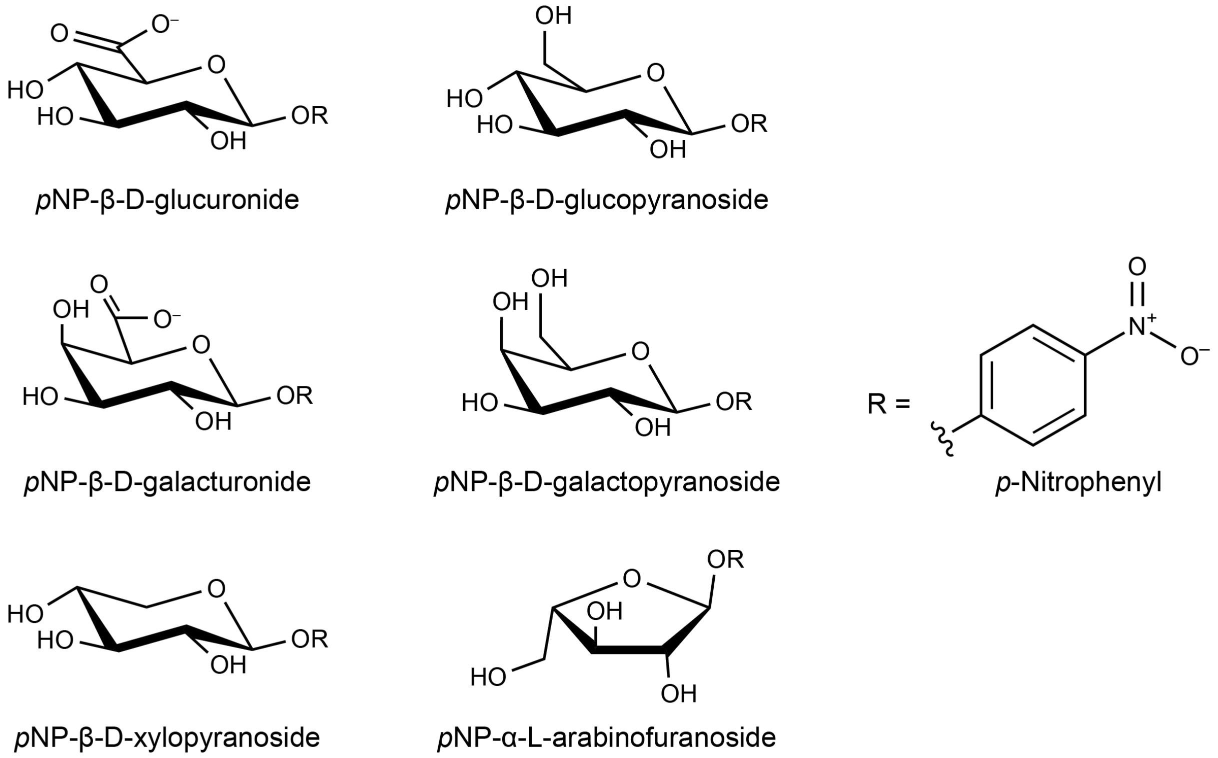


**Figure S13: Structures of model substrates (with 4-nitrophenyl leaving groups; R= $C_6H_4$-$NO_2$) that have been shown to be hydrolysed by SN243.**

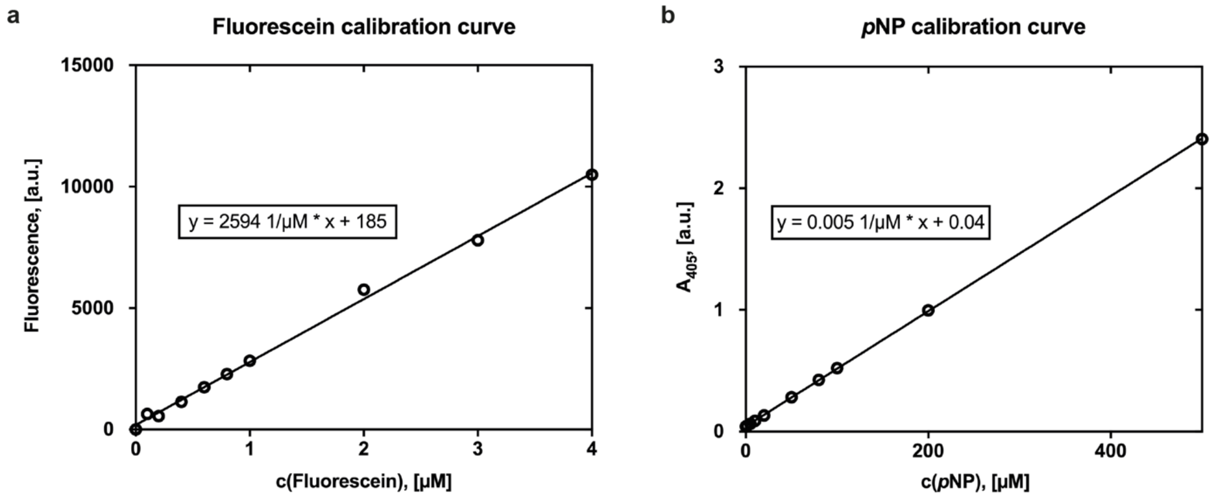


**Figure S14: Calibration curves for kinetic measurements.** Fluorescein **(a)** and *p*NP **(b)** in 100 mM Tris-HCl pH 8.0, 150 mM NaCl.

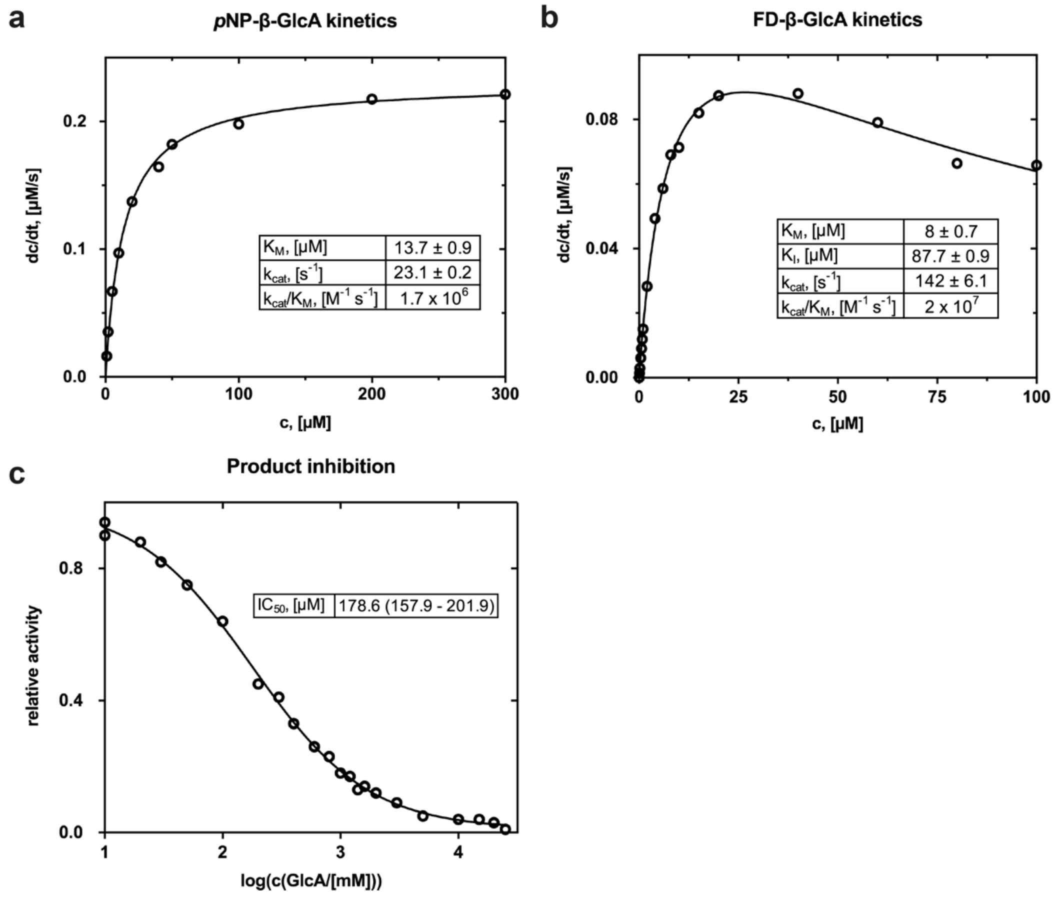


**Figure S15: Michaelis-Menten kinetics for the β-glucuronidase activity of SN243 at 25 °C.** Kinetic parameters were determined using *p*NP-β-GlcA **(a)** and FD-β-GlcA **(b)** as substrate and 10 nM and 1 nM enzyme, respectively. No inhibition was detected for the reaction with *p*NP-β-GlcA, while an inhibition constant of one order in magnitude higher than the $K_M$ was measured for FD-β-GlcA. **(c) Investigation of inhibitory effects of the reaction product GlcA on the β-glucuronidase activity of SN243.** Using 10 µM *p*NP-β-GlcA and 10 nM enzyme, the $IC_{50}$ was determined to be 179 µM. Fluorescein and *p*NP concentrations were determined using the respective calibration curves (Figure S14).

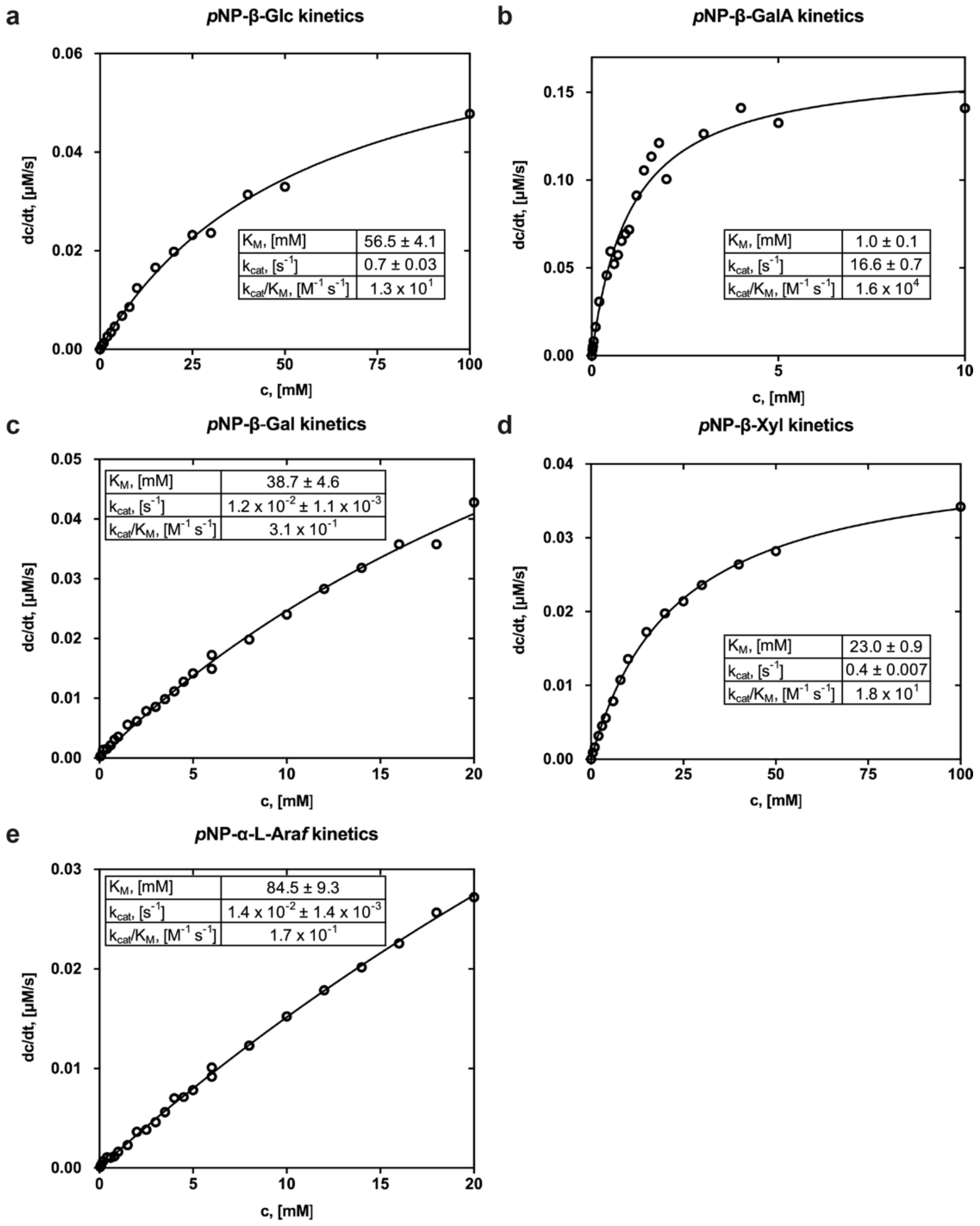


**Figure S16: Michaelis-Menten kinetics for promiscuous activities of SN243 at 25 °C.** In order to determine the Michaelis-Menten parameters for the hydrolysis of *p*NP-β-Glc **(a)**, *p*NP-β-GalA **(b),** *p*NP-β-Gal **(c)**, *p*NP-β-Xyl **(d)**, and *p*NP-α-L-Ara*f* **(e)** respectively, either 10 nM (*p*NP-β-GalA), 100 nM (*p*NP-β-Glc and *p*NP-β-Xyl) or 10 µM (*p*NP-β-Gal and *p*NP-α-L-Araf) SN243 were used. Limited substrate solubility in buffer prevented measurement of the full Michaelis Menten curve for *p*NP-β-Gal and *p*NP-α-L-Ara*f*. In cases where the saturation was not reached, the Michaelis-Menten parameters were extrapolations and thus carry larger errors. *p*NP concentrations were determined using the respective calibration curves (Figure S14).

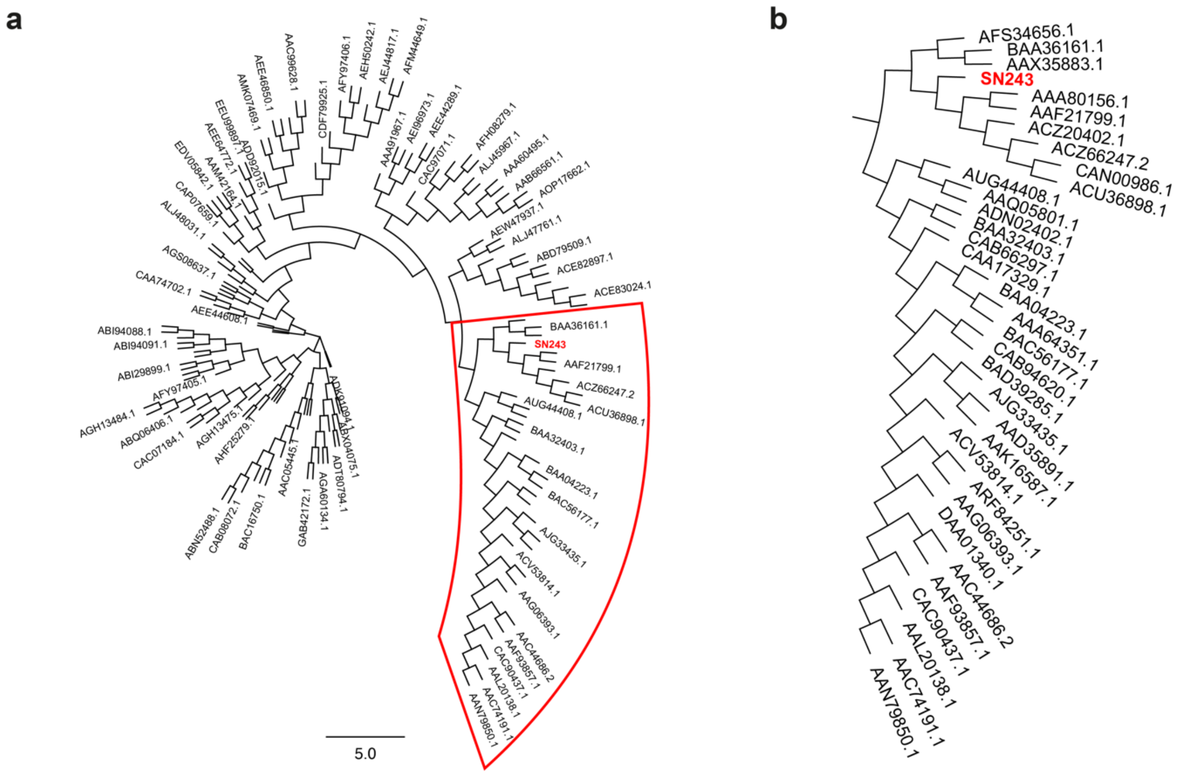


**Figure S17: Phylogenetic tree.** A multiple sequence alignment of SN243 and the characterized bacterial members of the GH3 family (CAZy database) was generated using ClustalW and displayed as a phylogenetic tree **(a)**. The read box highlights the 33 closest hits to SN243 and a close-up of this region is displayed in **(b)**. The multiple sequence alignment of SN243 with these 33 sequences served to identify the active side residues of SN243. Accession codes of genes are indicated.

a

| SN243 variant | $K_M$, [µM] | $k_{cat}$, [$s^{-1}$] | $k_{cat}/K_M$, [$M^{-1}s^{-1}$] |
|---|---|---|---|
| **wt** | 13.7 ± 0.9 | 23.1 ± 0.2 | 1.7 x $10^6$ |
| **D331A** | 1.1 ± 0.1 | 1.4 x $10^{-1}$ ± 0.3 x $10^{-1}$ | 1.3 x $10^5$ |
| **H333A** | 4.0 ± 0.2 | 3.1 x $10^{-1}$ ± 0.4 x $10^{-2}$ | 7.4 x $10^4$ |
| **D415A** | n.a | n.a | n.a |
| **H319A** | 61.1 x $10^1$ ± 3.2 x $10^1$ | 18.5 ± 0.4 | 3.0 x $10^4$ |
| **R136A** | 3.9 x $10^3$ ± 0.3 x $10^3$ | 45.3 ± 1.4 | 1.2 x $10^4$ |

b

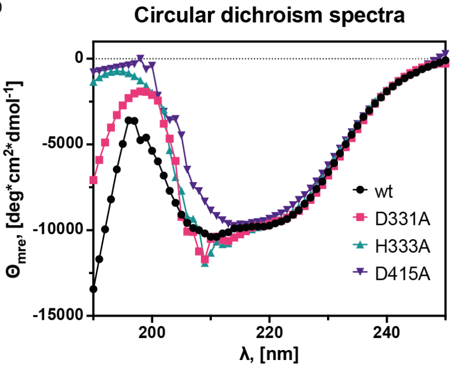


**Figure S18: Alanine scanning of the active site of SN243. (a) Michaelis-Menten kinetics for the β-glucuronidase activity of SN243 Ala mutants at 25 °C.** The role of the identified active site residues in catalysis was experimentally probed by testing the activity of x→Ala mutants towards *p*NP-β-GlcA. The catalytic nucleophile mutant D415A resulted in a complete loss of activity. The SN243 Ala variants for the acid-base dyad D331A and H333A have a 180-fold and 80-fold reduced $k_{cat}$, respectively, when compared to the wt enzyme. Mutation of residues involved in substrate binding (H319, R136) resulted in an increase in $K_M$. The corresponding Michaelis-Menten plots are given in Figure S19. **(b) Circular dichroism spectra of SN243 wt and Ala mutants of catalytic residues.** To ensure that reduced activity is caused by the point mutations and not an overall change in secondary structure CD spectra were recorded. All recorded spectra are very similar suggesting that the reduced or abolished activities of the SN243 Ala mutants are due to the exchange of an active site residues to Ala.

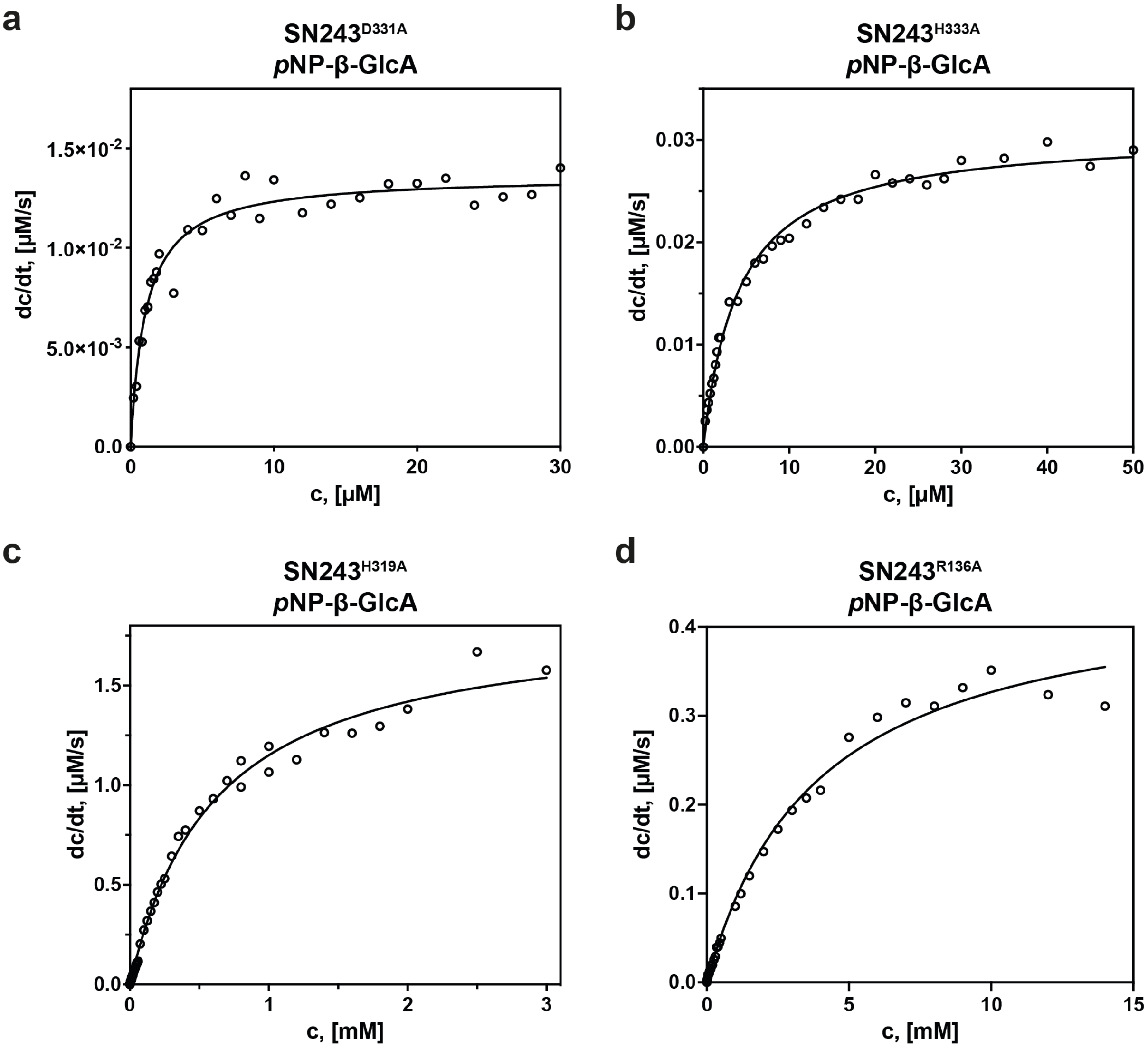


**Figure S19: Michaelis-Menten plots for the β-glucuronidase activity of SN243 active site mutants at 25 °C.** Kinetics were determined for the hydrolysis of *p*NP-β-GlcA using 100 nM enzyme (D331A, H333A) or 10 nM enzyme (H319A, R136A). Kinetic parameters are indicated in Figure S18a.

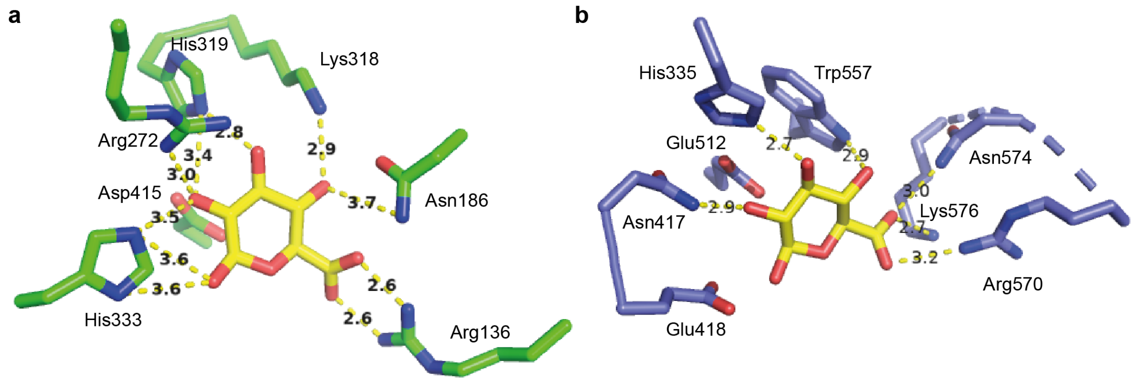


**Figure S20: Comparison of the binding of GlcA into the active site between of SN243 (green, panel a) and the GH2 β-glucuronidase from *Ruminococcus gnavus* (PDB: 6JZ5, purple, panel b).** While in SN243 Arg136 interacts via two short H-bonds with the C6-carboxylate of GlcA, this binding is achieved by three residues in GH2, Arg570, Asn574 and Lys576. Interestingly, the neighbouring O(C4) binds to Asn186 and Lys318 in SN243. However, these residues are far apart in the primary protein sequence and thus, do not mirror the conserved NxK motif of the GH2 family, which interact with the C6-carboxylate.

### 7. Supporting Tables

#### Table S2: Characterized bacterial β-glucuronidases contained in the CAZy database (release 22/01/21).

| Family | Protein name | Organism |
|---|---|---|
| GH2 | β-glucuronidase (Gus) | *Arthrobacter* sp. RP10 |
| GH2 | β-glucuronidase (Gus) | *Arthrobacter* sp. RP40.7 |
| GH2 | β-glucuronidase (BcellWH2_04226; BACWH2_3980; Lacz_22) | *Bacteroides cellulosilyticus* WH2 |
| GH2 | β-glucuronidase (BF0328) | *Bacteroides fragilis* NCTC 9343 |
| GH2 | β-glucuronidase (Bovatus_02309;BACOVA_03100;Uida_1) | *Bacteroides ovatus* ATCC 8483 |
| GH2 | β-glucuronidase (BMSP;HMPREF2532_04191) | *Bacteroides ovatus* KLE1656 |
| GH2 | bimodular β-L-arabinofuranosidase / β-glucuronidase (BT0996;BT_0996) | *Bacteroides thetaiotaomicron* VPI-5482 |
| GH2 | β-glucuronidase (BT4181;BT_4181) | *Bacteroides thetaiotaomicron* VPI-5482 |
| GH2 | β-glucuronidase (BuGUS-2; M094_3283) | *Bacteroides uniformis* str. 3978 T3 i |
| GH2 | β-glucuronidase (BuGUS-1; ERS417307_01040; Uida_4) | *Bacteroides uniformis* str. 3978 T3 i |
| GH2 | HMPREF0168_2111 (UidA) | *Bifidobacterium dentium* ATCC 27679 |
| GH2 | β-glucuronidase (BglR;CPE0147) | *Clostridium perfringens* str. 13 |
| GH2 | β-glucuronidase (UidA) | *Clostridium* sp. Marseille-P299 |
| GH2 | DICTH_1429 | *Dictyoglomus thermophilum* H-6-12 |
| GH2 | β-glucuronidase (UidA;GusA;GurA;b1617) | *Escherichia coli* str. K-12 substr. MG1655 |
| GH2 | β-glucuronidase (EUBELI_20590) | *[Eubacterium] eligens* ATCC 27750 |
| GH2 | β-glucuronidase (CG447_12395) | *Faecalibacterium prausnitzii* A2165 |
| GH2 | β-glucuronidase (C4Q21_10165) | *Faecalibacterium prausnitzii* APC918/95b |
| GH2 | β-glucuronidase (FP2_04120) | *Faecalibacterium prausnitzii* L2-6 L2/6 |
| GH2 | β-1,2-glucuronidase (BN863_22060) | *Formosa agariphila* KMM 3901 |
| GH2 | FsGUS (β-galacturonidase/β-glucuronidase) | *Fusicatenibacter saccharivorans* |
| GH2 | β-glucuronidase (HMPREF0539_0313; UidA) | *Lacticaseibacillus rhamnosus* LMS2-1 |
| GH2 | β-glucuronidase (GusA) | *Lactobacillus gasseri* ADH |
| GH2 | β-glucuronidase (Gus) | *Levilactobacillus brevis* RO1 |
| GH2 | β-glucuronidase (B160DRAFT_01711) | *Niabella aurantiaca* DSM 17617 |
| GH2 | β-glucuronidase (ZP_02032394.1;PARMER_02407)(26901) | *Parabacteroides merdae* ATCC 43184 |
| GH2 | β-glucuronidase (BSEG_01974) | *Phocaeicola dorei* 5_1_36/D4 |
| GH2 | β-glucuronidase | *Roseburia hominis* |
| GH2 | ERS852391_01800 (Uida_2) | *Roseburia hominis* 2789STDY5608834 |
| GH2 | β-glucuronidase | *Ruminococcus gnavus* |
| GH2 | β-glucuronidase (UidA) | *Ruminococcus gnavus* E1 |
| GH2 | β-glucuronidase (Gus) | *Staphylococcus sp.* RLH1 |
| GH2 | β-glucuronidase (SAG0698) | *Streptococcus agalactiae* 2603V/R |
| GH2 | β-glucuronidase (Cst_c06190;Lacz2;Uid2A) | *Thermoclostridium stercorarium* subsp. *stercorarium* DSM 8532 |
| GH2 | β-glucuronidase (Gus;TmGUS;TM1062;Tmari_1066) | *Thermotoga maritima* MSB8 |
| GH2 | β-glucuronidase (H11G11-BG) | uncultured bacterium |
| GH2 | β-glucuronidase (C7D2-BG) | uncultured bacterium |
| GH2 | Betα-glucuronidase (uidA_2) | uncultured *Clostridium* sp. |
| GH30 | β-glucuronidase (LcGUS30) | *Levilactobacillus brevis* subsp. *coagulans* FERM BP-4693 |
| GH79 | β-glucuronidase (ACP_2665;AcGlcA79A) (GlcA79A) | *Acidobacterium capsulatum* ATCC 51196 |
| GH79 | heparanase / endo-β-glucuronidase (BpHep;DP46_278) | *Burkholderia pseudomallei* BEK |
| GH154 | Arabinogalactan β-1,6-D-glucuronidase (BT3677; BT_3677) | *Bacteroides thetaiotaomicron* VPI-5482 |
| GH169 | β-1,4-glucuronidase (Pn3Pase; Pbac_3551) | *Paenibacillus* sp. 32352 |

**Table S3: Promiscuity survey for glycoside hydrolase activity of SN243 towards various *p*NP-coupled sugar substrates.** Activity could reliably be detected and quantified if at least a two-fold enhancement[a] compared to a no-enzyme control was observed using 2 mM substrate and 1 µM enzyme. Standard errors for Michaelis-Menten parameters are indicated with a 95% confidence level.

| Substrate[b] | Activity | $K_M$, [mM] | $k_{cat}$, [$s^{-1}$] | $k_{cat}/K_M$, [$M^{-1}$ $s^{-1}$] |
|---|---|---|---|---|
| *p*NP-β-GlcA | ✓ | $13.7 \times 10^{-3} \pm 0.9 \times 10^{-3}$ | 23.1 ± 0.2 | $1.7 \times 10^{6}$ |
| *p*NP-β-Glc | ✓ | 56.5 ± 4.1 | 0.7 ± 0.03 | $1.3 \times 10^{1}$ |
| *p*NP-β-GalA | ✓ | 1.0 ± 0.1 | 16.6 ± 0.7 | $1.6 \times 10^{4}$ |
| *p*NP-β-Gal[c] | ✓ | 38.7 ± 4.6 | $1.2 \times 10^{-2} \pm 1.1 \times 10^{-3}$ | $3.1 \times 10^{-1}$ |
| *p*NP-β-Xyl | ✓ | 23.0 ± 0.9 | 0.4 ± 0.007 | $1.8 \times 10^{1}$ |
| *p*NP-β-Man | - | - | - | - |
| *p*NP-β-GlcNAc | - | - | - | - |
| *p*NP-β-GalNAc | - | - | - | - |
| *p*NP-β-Lac | - | - | - | - |
| *p*NP-α-GlcA | - | - | - | - |
| *p*NP-α-Glc | - | - | - | - |
| *p*NP-α-GalA | - | - | - | - |
| *p*NP-α-Gal | - | - | - | - |
| *p*NP-α-Xyl | - | - | - | - |
| *p*NP-α-Man | - | - | - | - |
| *p*NP-α-L-Ara*f*[c] | ✓ | 84.5 ± 9.3 | $1.4 \times 10^{-2} \pm 1.4 \times 10^{-3}$ | $1.7 \times 10^{-1}$ |
| *p*NP-α-L-Rha | - | - | - | - |
| *p*NP-α-L-Fuc | - | - | - | - |

[a]A two-fold enhancement corresponds to $A_{405}$ at least twice as high as for the uncatalysed reaction. An extrapolation using a calibration curve of absorbance as a function of 4-nitrophenolate (Figure S14B) suggests that > 8 µM product of the reaction of *p*NP-β-GlcA are required for detection of activity. After 10 h reaction time (where uncatalyzed product formation is undetectable) and 1 µM enzyme, activity with a minimmal $k_{cat}$ of $2.2 \times 10^{-4}$ $s^{-1}$ could be detected. The highest $K_M$ experimentally measured in this setup was 85 mM. Therefore, we assume that the assay is sufficiently sensitive to detect a $k_{cat}/K_M$ that is $10^{8}$-fold lower than the corresponding values for SN243 for the hydrolysis of *p*NP-β-GlcA.
[b]Chemical structures of substrates, for which activity was detected, are shown in Figure S13.
[c]Limited solubility of the substrates in buffer prevented reliable determination of the Michaelis Menten parameters. The indicated values ($K_M$ and $k_{cat}$) for the reaction with *p*NP-β-Gal and *p*NP-α-L-Ara*f* are only estimates resulting from measurements plotting the start of the Michaelis Menten curve.

**Table S4: Comparison of catalytic parameters for the hydrolysis of pNP-β-GlcA by SN243 and other bacterial β-glucuronidases.**

| Enzyme | $K_M$, [μM] | $k_{cat}$, [$s^{-1}$] | $k_{cat}/K_M$, [$M^{-1}s^{-1}$] | Ref. |
|---|---|---|---|---|
| SN243 | 10.5 ± 1.7 | 44.6 ± 1.5 | $4.2 \times 10^6$ | this work |
| *Thermotoga maritima* GUS | 150 ± 10 | 68 ± 2 | $4.5 \times 10^5$ | [17] |
| *Escherichia coli* GUS | 130 ± 10 | 120 ± 12 | $9.2 \times 10^5$ | [18] |
| *Streptococcus agalactiae* GUS | 360 ± 30 | 80 ± 2 | $2.2 \times 10^5$ | [18] |
| | 170 ± 10 | 122 ± 3 | $7.2 \times 10^5$ | [20] |
| *Bacteroides uniformis* GUS-1 | 57 ± 9 | 12 ± 1 | $2.2 \times 10^5$ | [19] |
| *Eubacterium eligens* GUS | 223 ± 8 | 120 ± 8 | $5.4 \times 10^5$ | [20] |
| *Clostridium perfringens* GUS | 160 ± 20 | 57 ± 2 | $3.6 \times 10^5$ | [20] |
| *Faecalibacterium prausnitzii* GUS | 2200 ± 300 | 58 ± 4 | $2.7 \times 10^4$ | [20] |
| *Lactobacillus rhamnosus* GUS | 1500 ± 100 | 10 ± 0.7 | $6.7 \times 10^3$ | [20] |
| *Ruminococcus gnavus* GUS | 1400 ± 0.2 | 2.8 ± 0.3 | $2.0 \times 10^3$ | [20] |
| *Bacteroides fragilis* GUS | 580 ± 90 | 22 ± 2 | $3.9 \times 10^4$ | [20] |
| *Parabacteroides merdae* GUS | 2400 ± 70 | $8.8 \times 10^{-2} \pm 0.1 \times 10^{-2}$ | $3.7 \times 10^1$ | [20] |
| *Bacteroides ovatus* GUS | 1380 ± 10 | $4.8 \times 10^{-1} \pm 0.3 \times 10^{-1}$ | $3.4 \times 10^2$ | [20] |
| *Bacteroides dorei* GUS | 1400 ± 100 | 7.5 ± 0.4 | $5.2 \times 10^3$ | [20] |
| *Fusicatenibacter saccharivorans* GUS | 30 ± 6 | 15 ± 1 | $5.0 \times 10^5$ | [21] |
| *Bifidobacterium dentium* GUS | 1359 ± 228 | 113 ± 8 | $8.3 \times 10^4$ | [22] |
| *Dictyoglomus thermophilum* GUS | 125 ± 20 | 9.3 ± 0.8 | $7.4 \times 10^4$ | [23] |

**Table S5: Data collection, processing and refinement statistics for SN243 crystal structures.**

| **PDB code** | **7QE1** | **7QE2** | **7QEF** | **7QEA** | **7QG4** | **7QEE** |
|---|---|---|---|---|---|---|
| SN243 variant | wt | wt | D415A | D415A | D415N | D415N |
| Ligand | *apo* | GlcA | *p*NP-β-GlcA | FD-β-GlcA | *apo* | *p*NP-β-GlcA |
| **Data collection** | | | | | | |
| Collection Date | 24/02/2021 | 23/04/2021 | 22/06/2021 | 22/06/2021 | 22/06/2021 | 22/06/2021 |
| Synchrotron | Diamond | Diamond | Diamond | Diamond | Diamond | Diamond |
| Beamline | i24 | i04 | i24 | i24 | i24 | i24 |
| X-ray wavelength | 0.9999 | 0.9795 | 0.9999 | 0.9999 | 0.9999 | 0.9999 |
| **Data processing*** | | | | | | |
| Space group | P1 | P1 | P1 | P1 | P1 | P1 |
| Unit cell (a, b, c [Å]) | 53.28, 90.76, 103.29 | 63.23, 82.18, 93.18 | 63.38, 81.68, 93.55 | 63.27, 81.57, 93.39 | 62.60, 82.77, 92.44 | 62.45, 82.17, 93.04 |
| Unit cell (α, β, γ [°]) | 67.22, 84.03, 76.91 | 66.74, 89.26, 89.34 | 66.57, 89.39, 89.17 | 66.64, 89.38, 89.57 | 113.33, 90.29, 91.04 | 113.69, 90.65, 90.31 |
| Resolution limits, [Å] | 51.89-1.95 (2.00-1.95) | 48.31-2.15 (2.21-2.15) | 85.83-2.41 (2.54-2.41) | 74.88-2.28 (2.40-2.28) | 75.98-2.08 (2.15-1.89) | 75.24-2.37 (2.50-2.37) |
| Total/unique reflections | 735719/123601 | 325221/91233 | 130173/62881 | 171343/75860 | 347587/100349 | 181699/65955 |
| Multiplicity | 6.0 (5.7) | 3.6 (3.7) | 2.1 (1.9) | 2.3 (2.3) | 3.5 (3.5) | 2.8 (1.6) |
| $R_{merge}$ | 0.073 (2.886) | 0.107 (1.650) | 0.122 (0.881) | 0.095 (0.876) | 0.094 (1.670) | 0.136 (0.762) |
| $R_{meas}$ | 0.080 (3.172) | 0.126 (1.930) | 0.160 (1.188) | 0.124 (1.151) | 0.130 (2.360) | 0.165 (1.078) |
| I/σI | 9.0 (0.6) | 7.1 (0.9) | 6.5 (1.8) | 3.8 (0.8) | 5.3 (0.6) | 2.5 (0.7) |
| $CC_{1/2}$ | 0.998 (0.371) | 0.995 (0.353) | 0.986 (0.412) | 0.992 (0.452) | 0.995 (0.404) | 0.988 (0.457) |
| Completeness, [%] | 97.6 (95.9) | 97.3 (97.3) | 94.5 (94.6) | 97.0 (97.2) | 97.8 (97.0) | 96.3 (92.3) |
| **Refinement** | | | | | | |
| $R_{work}$/$R_{free}$, [%] | 0.2403/0.2904 | 0.1885/0.2463 | 0.2150/0.2855 | 0.1977/0.2530 | 0.2488/0.2508 | 0.2168/0.2729 |
| Unique/free reflections used | 123265/ 6070 | 325221/91233 | 61589/3061 | 75285/3978 | 91984/4934 | 65533/3410 |
| R.m.s deviations: | | | | | | |
| bond lengths, [Å] | 0.009 | 0.015 | 0.009 | 0.009 | 0.009 | 0.009 |
| bond angles, [°] | 1.239 | 1.580 | 1.179 | 1.305 | 1.119 | 1.240 |
| Ramachandran analysis: | | | | | | |
| Most favoured | 1416 (96.5%) | 1434 (97.7%) | 1432 (97.5%) | 1430 (97.4%) | 1433 (96.3%) | 1414 (96.3%) |
| Allowed | 44 (3.0%) | 34 (2.3%) | 34 (2.3%) | 38 (2.6%) | 55 (3.7%) | 52 (3.5%) |
| Outliers | 7 (0.5%) | 0 (0.0%) | 2 (0.1%) | 0 (0.0%) | 0 (0.0%) | 2 (0.1%) |
| Number of atoms: | | | | | | |
| Protein | 11218 | 11256 | 11250 | 11250 | 11401 | 11247 |
| Solvent atoms | 92 | 376 | 170 | 328 | 451 | 110 |
| Ligand atoms | 27 | 87 | 63 | 39 | 31 | 68 |
| Heterogen atoms | 11337 | 11719 | 11527 | 11739 | 11881 | 11459 |
| Mean B-factor, [$Å^2$] | 72.83 | 61.77 | 50.42 | 53.43 | 42.99 | 57.96 |

*Values for the high-resolution shell are given in parenthesis.

#### 8. Supporting References


1 Peränen, J., Rikkonen, M., Hyvönen, M. & Kääriäinen, L. T7 Vectors with a Modified T7lacPromoter for Expression of Proteins inEscherichia coli. *Analytical Biochemistry* **236**, 371-373, doi:https://doi.org/10.1006/abio.1996.0187 (1996).

2 Mitchell, A. L. *et al.* MGnify: the microbiome analysis resource in 2020. *Nucleic Acids Res* **48**, D570-D578, doi:10.1093/nar/gkz1035 (2020).

3 Consortium, T. U. UniProt: the universal protein knowledgebase in 2021. *Nucleic Acids Research* **49**, D480-D489, doi:10.1093/nar/gkaa1100 (2020).

4 Atkinson, H. J., Morris, J. H., Ferrin, T. E. & Babbitt, P. C. Using Sequence Similarity Networks for Visualization of Relationships Across Diverse Protein Superfamilies. *PLOS ONE* **4**, e4345, doi:10.1371/journal.pone.0004345 (2009).

5 Shannon, P. *et al.* Cytoscape: A Software Environment for Integrated Models of Biomolecular Interaction Networks. *Genome Research* **13**, 2498-2504, doi:10.1101/gr.1239303 (2003).

6 Huang, Y., Niu, B., Gao, Y., Fu, L. & Li, W. CD-HIT Suite: a web server for clustering and comparing biological sequences. *Bioinformatics* **26**, 680-682, doi:10.1093/bioinformatics/btq003 (2010).

7 Fincher, G. B., Stone, B. A. & Clarke, A. E. Arabinogalactan-Proteins: Structure, Biosynthesis, and Function. *Ann Rev PLan Physiol* **34**, 47-70, doi:https://doi.org/10.1146/annurev.pp.34.060183.000403 (1983).

8 Cartmell, A. *et al.* A surface endogalactanase in Bacteroides thetaiotaomicron confers keystone status for arabinogalactan degradation. *Nat Microbiol* **3**, 1314-1326, doi:10.1038/s41564-018-0258-8 (2018).

9 Williams, P. A. & Phillips, G. O. in *Handbook of Hydrocolloids (Third Edition)* (eds Glyn O. Phillips & Peter A. Williams) 627-652 (Woodhead Publishing, 2021).

10 Tryfona, T. *et al.* Structural Characterization of Arabidopsis Leaf Arabinogalactan Polysaccharides *Plant Physiology* **160**, 653-666, doi:10.1104/pp.112.202309 (2012).

11 Takata, R. *et al.* Degradation of carbohydrate moieties of arabinogalactan-proteins by glycoside hydrolases from Neurospora crassa. *Carbohydrate Research* **345**, 2516-2522, doi:https://doi.org/10.1016/j.carres.2010.09.006 (2010).

12 Tsumuraya, Y., Mochizuki, N., Hashimoto, Y. & Kovác, P. Purification of an exo-beta-(1----3)-D-galactanase of Irpex lacteus (Polyporus tulipiferae) and its action on arabinogalactan-proteins. *Journal of Biological Chemistry* **265**, 7207-7215, doi:https://doi.org/10.1016/S0021-9258(19)39100-8 (1990).

13 Goubet, F., Jackson, P., Deery, M. J. & Dupree, P. Polysaccharide Analysis Using Carbohydrate Gel Electrophoresis: A Method to Study Plant Cell Wall Polysaccharides and Polysaccharide Hydrolases. *Analytical Biochemistry* **300**, 53-68, doi:https://doi.org/10.1006/abio.2001.5444 (2002).

14 Colin, P. Y. *et al.* Ultrahigh-throughput discovery of promiscuous enzymes by picodroplet functional metagenomics. *Nat Commun* **6**, 10008, doi:10.1038/ncomms10008 (2015).

15 Almagro Armenteros, J. J. *et al.* SignalP 5.0 improves signal peptide predictions using deep neural networks. *Nat Biotechnol* **37**, 420-423, doi:10.1038/s41587-019-0036-z (2019).

16 Krogh, A., Larsson, B., von Heijne, G. & Sonnhammer, E. L. Predicting transmembrane protein topology with a hidden Markov model: application to complete genomes. *J Mol Biol* **305**, 567-580, doi:10.1006/jmbi.2000.4315 (2001).

17 Salleh, H. M. *et al.* Cloning and characterization of Thermotoga maritima β-glucuronidase. *Carbohydrate Research* **341**, 49-59, doi:https://doi.org/10.1016/j.carres.2005.10.005 (2006).

18 Wallace, B. D. *et al.* Structure and Inhibition of Microbiome beta-Glucuronidases Essential to the Alleviation of Cancer Drug Toxicity. *Chem Biol* **22**, 1238-1249, doi:10.1016/j.chembiol.2015.08.005 (2015).

19 Pellock, S. J. *et al.* Three structurally and functionally distinct β-glucuronidases from the human gut microbe Bacteroides uniformis. *Journal of Biological Chemistry* **293**, 18559-18573, doi:https://doi.org/10.1074/jbc.RA118.005414 (2018).

20 Biernat, K. A. *et al.* Structure, function, and inhibition of drug reactivating human gut microbial β-glucuronidases. *Scientific Reports* **9**, 825, doi:10.1038/s41598-018-36069-w (2019).

21 Pellock, S. J., Walton, W. G. & Redinbo, M. R. Selecting a Single Stereocenter: The Molecular Nuances That Differentiate β-Hexuronidases in the Human Gut Microbiome. *Biochemistry* **58**, 1311-1317, doi:10.1021/acs.biochem.8b01285 (2019).

22 Dashnyam, P. *et al.* β-Glucuronidases of opportunistic bacteria are the major contributors to xenobiotic-induced toxicity in the gut. *Scientific Reports* **8**, 16372, doi:10.1038/s41598-018-34678-z (2018).

23 Kurdziel, M. *et al.* Thioglycoligation of aromatic thiols using a natural glucuronide donor. *Organic & Biomolecular Chemistry* **18**, 5582-5585, doi:10.1039/D0OB00226G (2020).
